# SubCell: Proteome-aware vision foundation models for microscopy capture single-cell biology

**DOI:** 10.1101/2024.12.06.627299

**Authors:** Ankit Gupta, Zoe Wefers, Konstantin Kahnert, Jan N. Hansen, Mohini K. Misra, Will Leineweber, Anthony Cesnik, Dan Lu, Ulrika Axelsson, Frederic Ballllosera, Russ B. Altman, Theofanis Karaletsos, Emma Lundberg

## Abstract

Cell morphology and subcellular protein organization provide important insights into cellular function and behavior. These cellular features can be studied using large-scale fluorescence microscopy, and machine learning has become a powerful tool to interpret the resulting images for biological insights. Here, we introduce SubCell, a deep learning model for fluorescence microscopy designed to accurately capture cellular morphology, protein localization, cellular organization, and biological function beyond what humans can readily perceive. SubCell was trained on the proteome-wide image collection from the Human Protein Atlas with a novel proteome-aware learning objective. SubCell outperforms state-of-the-art methods across a variety of tasks relevant to single-cell biology and generalizes to other fluorescence microscopy datasets without any fine-tuning. Additionally, we construct the first proteome-wide hierarchical map of proteome organization that is directly learned from image data. This vision-based multiscale cell map defines cellular subsystems down to protein complex resolution, reveals proteins with similar functions, and distinguishes dynamic and stable behaviors within cellular compartments. Finally, combining SubCell with a protein sequence model enables a rich multimodal approach to capture gene function better than either vision-only or sequence-only models alone. In conclusion, SubCell creates deep, image-driven representations of cellular architecture that are applicable across diverse biological contexts and datasets.

## Main

Proper protein localization is essential for cells to function. Changes in protein localization induce cells to adopt diverse morphologies that orchestrate complex biological processes, and mislocalization is highly associated with disease states [1–6]. Therefore, understanding cell behavior requires comprehensive spatial mapping of cell morphology, subcellular protein organization, and dynamics, which can currently only be captured at scale by microscopy [7]. Consequently, fluorescence microscopy has become an indispensable tool for systematically characterizing protein localization [8–10] and for profiling morphological responses to chemical and genetic perturbations at scale [11–17].

The massive scale of high-throughput fluorescence microscopy datasets requires automated analysis methods. Profiling tools [3] and machine learning approaches [18–22] that create single-cell feature embeddings have demonstrated faster and deeper analysis of large image datasets beyond human capability. Self-supervised learning has emerged as a particularly powerful paradigm for general feature representation in biology [23–25]; however, bespoke models are typically trained separately for each dataset and for different biological tasks. The proliferation of machine learning models with narrow applicability across different datasets has created a critical need for an easy-to-use feature extractor that matches the performance of task-specific models across different imaging modalities.

Here, we present SubCell, a vision model for representing single cells in fluorescence microscopy images. Trained on the Human Protein Atlas (HPA) proteome-wide dataset that provides unmatched cellular pattern diversity [8], SubCell utilizes a multitask learning framework to efficiently encode both cellular phenotypes and subcellular protein organization (Fig. 1A, B). Without fine-tuning, SubCell produces robust representations of cell morphology and protein localization across diverse, independent datasets that vary widely in image resolution, reference marker stainings, cell types, and species. The single-cell representations enable a wide range of downstream tasks, including protein localization and cell line classification, cell-cycle modeling, drug-response prediction, and mechanism-of-action identification (Fig. 1C-E). Further, by capturing subtle, continuous patterns of protein localization, SubCell enables the creation of the first image-based multiscale map of subcellular protein organization. Finally, combining SubCell with protein sequence representations yields a latent space enriched with signals for functional genomics, facilitating a multimodal systems-level understanding of proteins and cells.

**Fig. 1.**
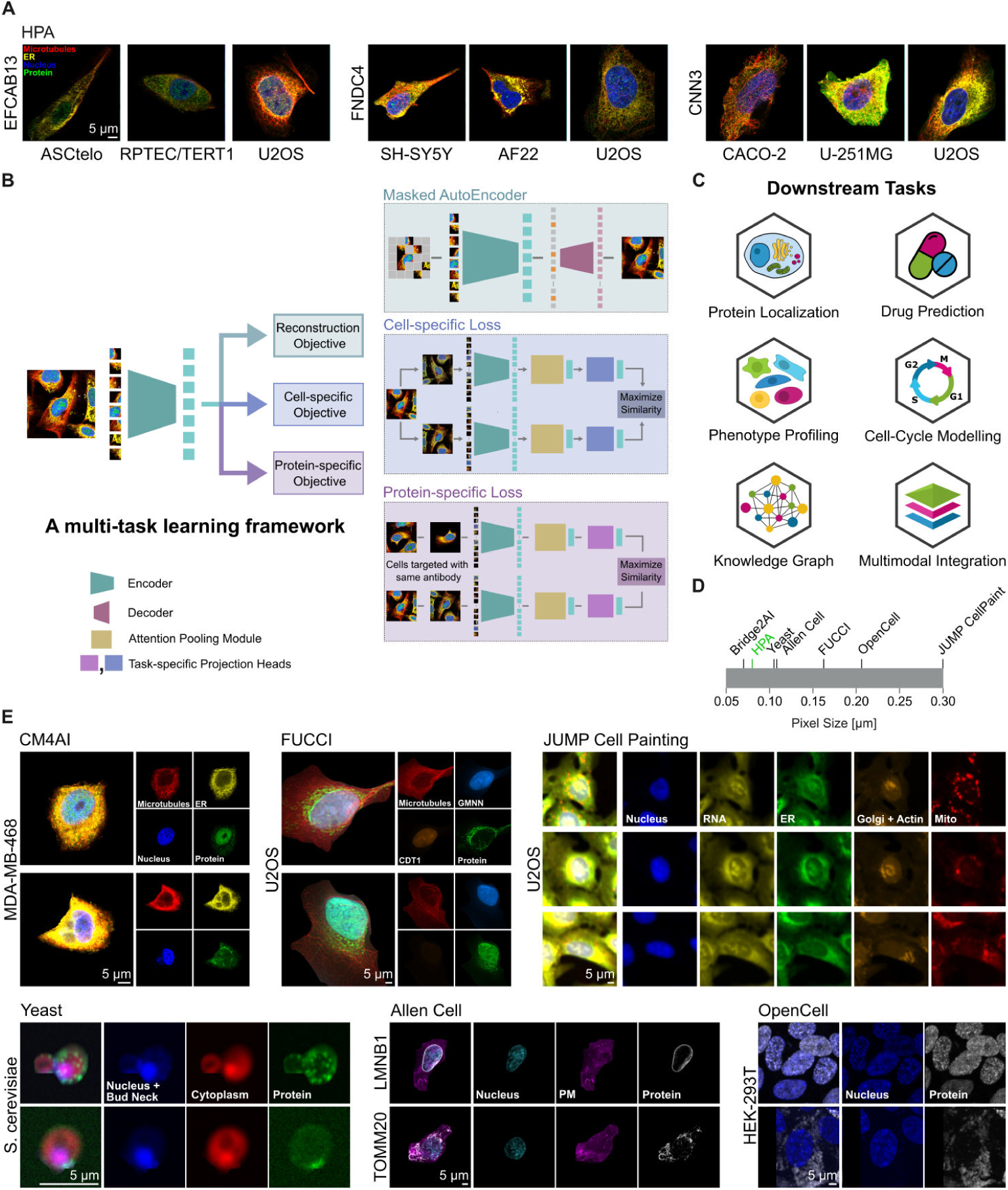
Overview of the SubCell learning framework. (A) Representative images of single-cell crops from the Human Protein Atlas dataset demonstrating the diverse cell morphologies and protein subcellular localizations present in a wide range of human cells (blue: nucleus; red: microtubules; yellow: endoplasmic reticulum; green: protein of interest). The images show three different proteins (EFCAB13, FNDC4 and CNN3) in a variety of human primary and cancer cell lines derived from different organs of origin in donors of different age and sex (ASC52telo: mesenchyme, adipose tissue; RPTEC/TERT1: epithelium, kidney; U2OS: osteosarcoma; SH-SY5Y: brain; AF22: neuroepithelial stem cells; CACO-2: adenocarcinoma; U-251MG: glioblastoma (more details see Materials and Methods)) (B) Illustration depicting our multi-task learning approach to train vision transformer models. We use three objectives to train our models: reconstruction, cell-specific, and protein-specific objectives. (C) Examples of the downstream tasks that are enabled with SubCell shown in this work. (D) The pixel size of each of the datasets used for evaluating SubCell (HPA, highlighted in green, was used for training). (E) Representative images from the datasets used in the evaluations to showcase diversity in resolution and imaging channel: (top, left to right) CM4AI, FUCCI U2OS, and JUMP cell paint datasets, and (bottom, left to right) yeast cell cycle, Allen Cell, and OpenCell datasets.

### Proteome-aware single-cell vision model

We developed generalizable vision models for fluorescence microscopy images to capture a broad range of cellular characteristics, including protein localization patterns and cell morphology. Since proteins are key structural and functional molecules in cells, we reasoned that a proteome-based training framework would enable the model to learn attention patterns relevant to cellular function. For this purpose, we used the HPA (v23) dataset, which contains images of 1.13 million single cells, encoding the expression and subcellular distribution of 13,141 genes across 37 human cell lines representing cells of different developmental and tissue origins with vastly different phenotypes (e.g., suspension cells, epithelial cells, fibroblasts). The HPA was chosen for its broad coverage of cell lines and protein subcellular localization patterns, as well as its high image resolution (Fig. 1A).

We examined various state-of-the-art unsupervised learning methods, including masked autoencoders (MAEs), and self-supervised learning techniques such as contrastive learning and self-distillation. We found that these techniques learned morphology well but fell short in encoding protein subcellular localization patterns (See “Model Development” in Methods). In addition to the standard reconstruction objective, we introduced the concepts of “cell-specific” and “protein-specific” objectives (Fig. 1B), which guide the models with biologically relevant information about the images (see Methods). The cell-specific objective enabled the model to learn cell-identity features, whereas the protein-specific objective helped the model understand protein localization. We combined the three objectives in a multitask learning framework (Fig. 1B, detailed in Supplementary Fig. S1) to train a vision transformer (ViT) model [26] with 86.4 million parameters, thus achieving a generalized representation of single-cell images. We added an attention-pooling module to the encoder’s output to prevent the model from learning spurious background features and optimized the multitask learning framework by evaluating combinations of learning objectives.

After experimenting with different learning objectives (see “Model Development” in Methods), we found that the best-performing models resulted from using the multitask training objective of combining reconstruction, cell-specific, and protein-specific losses, which we coined “SubCell”, and from using only the protein-specific objective, which we coined “SubCell-P”.

### Performance on Benchmarks

We benchmarked the protein localization performance of SubCell (and SubCell-P) against current state-of-the-art models trained on HPA images, including the supervised “bestfitting” model that won a Kaggle competition for protein localization prediction [27] and the self-supervised DINO4Cells-HPA model [19]. We also included a weakly supervised model with a ViT architecture identical to SubCell (ViT-Weak-Supervised), allowing us to explore the contribution of SubCell’s training objective to performance. We evaluated performance on a held-out HPA test set and the hidden HPA Kaggle challenge test set [27] (Fig. 2A). On both datasets, SubCell outperformed the state-of-the-art self-supervised model (DINO4Cells-HPA) and the baseline ViT-Weak-Supervised model, highlighting the advantage of SubCell’s new model architecture. SubCell fell slightly short compared to the supervised “bestfitting” model on the Kaggle dataset. A breakdown by protein localization category revealed that this difference was mainly due to rare classes (Fig. 2B), such as the mitotic spindle and centrosome. These classes are particularly challenging because they represent patterns visible in only a few cells, resulting in sparse training data. Centrosomes, for example, are small punctate structures that may not be present in the image because they lie outside the z-plane of the image or exhibit low signal-to-noise ratios. As a consequence, both human annotators and models perform with lower confidence on these classes. The bestfitting model achieves better performance on rare classes by boosting rare-class representation using specialized techniques, whereas SubCell classifiers did not employ any such techniques.

**Fig. 2.**
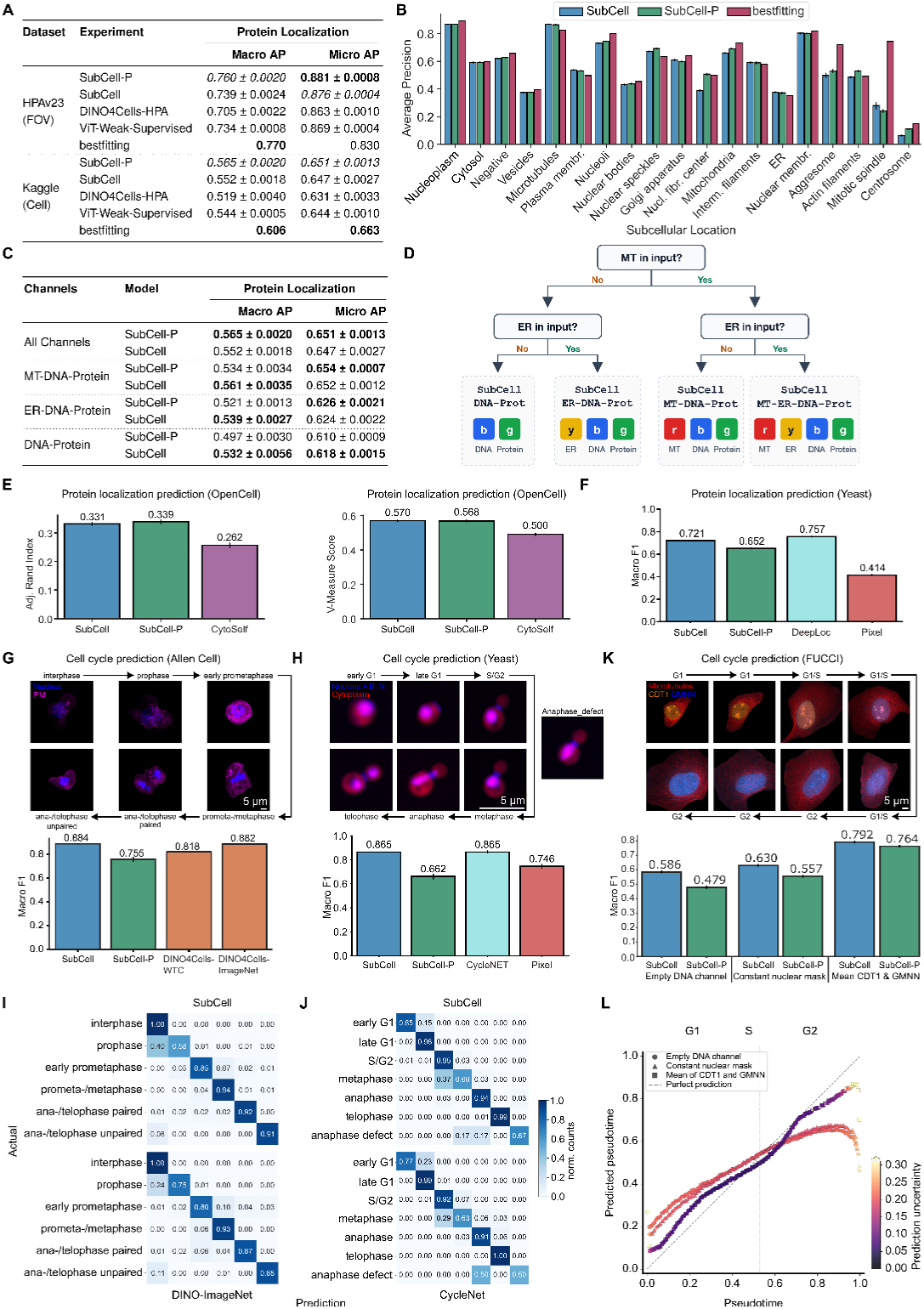
Quantitative evaluation of learned representations for predicting protein localization and cell cycle stages across different datasets. (A) Results of protein localization prediction on the HPA test set (FOV level) and hidden Kaggle test set (single-cell level), showing oveall macro and micro average precision. The best performance for each metric is highlighted in bold, and the second-best is highlighted in italics. (Results are obtained by training the MLP classifiers on the features obtained by the models. The classifiers were trained 10 times with different random seeds, and the mean and standard deviation are reported. Since bestfitting is a supervised model, the results were reported as is. Similar analysis is shown in (C).) (B) Protein localization results by category on the Kaggle test set. Bar plots show the average precision for each category for bestfitting (magenta), SubCell-P (green), and SubCell (blue). (C) Evaluation of our models trained with different channel combinations for protein localization prediction and cell-line classification on the Kaggle test set (the best performance for each metric is highlighted in bold). (D) Decision tree indicating which SubCell variant is chosen based on the channels present in the dataset. The protein channel can be replaced with any new channel in the dataset that’s not present in HPA. (E) Protein localization clustering performance of the models on the OpenCell dataset, evaluated using adjusted Rand index (left) and V-measure score (right). (Clustering was performed 10 times with different seeds for K-Means clustering, and the error bar represents the standard deviation of the metrics.) (F) Protein localization classification performance on the yeast cell dataset, with bar plots showing the macro F1-score. Error bars show the standard deviation across ten models trained with ten different random seeds. (G-H) Cell cycle stage classification in the Allen Cell dataset (G) and yeast dataset (H). Top: Example images for the different cell cycle stages in the data set. Bottom: Macro F1 scores for Cell Cycle Stage Classification. Error bars show standard deviation across ten random seeds. No error bar is shown for the DINO results because the classifier weights provided by the original authors were used. Abbreviation B.N.: Bud Neck. (I-J) Confusion matrices of the two best classification models in the AllenCell dataset (normalized by the ground truth category counts) (I) and the yeast dataset (J), showing the actual vs predicted cell cycle stages. (K) Cell cycle stage classification in the FUCCI U2OS dataset, which also allows pseudo-timing based on the nuclear intensity of the cell-cycle-related proteins CDT1 (orange) and GMNN (blue). Top: Example images ordered by decreasing CDT1 and increasing GMNN intensity, annotated with discrete cell cycle stages (G1, G1/S, G2). Bottom: Macro F1-scores for cell cycle stage classification. Because SubCell requires a nuclear stain input, we evaluated the FUCCI dataset with different configurations for replacing the nuclear channel: (1) an empty nuclear channel, (2) a binary nuclear mask with constant intensity inside the mask and zero intensity outside the mask, and (3) the mean of the CDT1 and GMNN channels. The results were obtained by gathering predictions from classifiers trained on the features using ten-fold cross-validation, with ten classifiers trained for each fold with different seeds. (L) The plot displays the results of the pseudotime deep evidential regression model for the three different configurations, showing true pseudotime on the (x-axis) against predicted pseudotime (y-axis). The color of the markers indicates the evidential uncertainty of the prediction of each model at a particular pseudotime (darker colors show less uncertainty, brighter colors show higher uncertainty). Perfect prediction is indicated by the dashed diagonal line, and pseudotime intervals corresponding to G1, S, and G2 phases are marked by the vertical dashed lines. (Similar analysis as (G), but with regression heads instead of classifiers.)

To assess generalizability, we comprehensively evaluated SubCell on external datasets that differ from HPA in microscopy modalities, image resolution, reference marker combinations, and species, such as the OpenCell dataset [9] and a yeast dataset [28] (Fig. 1D, 1E). To enable such a general application, we trained variants of SubCell and SubCell-P that operate with fewer reference markers than in the HPA. For example, HPA images include three cellular reference markers staining the nucleus (DNA), endoplasmic reticulum (ER), and microtubules (MT), while the OpenCell dataset includes only the nucleus (DNA) (Fig. 1A, E). Performance across different channel combinations was broadly comparable, indicating that SubCell is applicable with fewer reference channels, though performance slightly improved when the ER marker was removed, worsened upon removal of the MT marker, and was worst with only DNA as a reference (Fig. 2C). We provide users with automatic model selection based on the available channel combinations in our repository (Fig. 2D, Example: Supplementary Fig. S2).

When evaluating the OpenCell dataset, SubCell outperformed CytoSelf [22], a self-supervised model trained directly on that dataset (Fig. 2E), on the clustering metrics. On the yeast dataset, SubCell-based classifiers significantly outperformed a simple pixel-based MLP (multi-layer perceptron) baseline, demonstrating that SubCell captures meaningful biological features better than MLPs trained on raw pixel intensities, even for the small yeast images (64×64 pixels) that had little resemblance to the training data. Additionally, SubCell nearly matched the performance of a supervised model trained specifically on the yeast images (DeepLoc) [28] (Fig. 2F). DeepLoc’s advantage likely stems from its exposure to dataset-specific characteristics, including low resolution (64×64 pixels), noise patterns, and species-specific morphology, which SubCell, trained on high-resolution human cell images, has not encountered during training.

Generally, SubCell and SubCell-P performed comparably for protein localization, while SubCell performed better when not all channels were provided (Fig. 2C) or when applied to out-of-distribution datasets (Fig. 2E, F). Together, these results show that SubCell generalizes across diverse microscopy datasets, achieving competitive or superior protein localization performance without any dataset-specific fine-tuning.

Beyond protein localization, we examined whether SubCell captures cell morphology by testing its ability to represent cell cycle stages – a process driven by coordinated morphological and molecular transformations [1, 28–30]. We assessed performance on three unseen datasets with cell cycle annotations: human induced Pluripotent Stem Cells (hiPSC) (AllenCell dataset) [10], yeast cells [28], and FUCCI-U2OS cells [1] (see Methods for dataset details, Fig. 1E). On the AllenCell dataset, SubCell outperformed two DINO-based models trained on either the AllenCell dataset itself [19] or on ImageNet [31] (Fig. 2G). On the yeast dataset, SubCell matched the performance of CycleNet, a supervised model trained specifically for cell cycle classification [28]; SubCell outperformed a classifier trained on the pixel-based baseline model, confirming that it extracts meaningful morphological features even from the small yeast images (Fig. 2H). Notably, the best results for SubCell on the yeast data were obtained using a simple logistic regression classifier (Supplementary Table S1), suggesting that the embedding space learned by the model linearly separates cell cycle classes and is rich in morphological information (see Supplementary Fig. S3). Classification errors in both datasets (AllenCell, yeast) mainly resulted from confusion between specific cycle stages that are visually difficult to distinguish: interphase and prophase in the hiPSC data, or S/G2 and metaphase in the yeast data (Fig. 2I, J).

We next tested whether SubCell embeddings support fine-grained cell-cycle pseudotime analysis using the FUCCI-U2OS dataset, which provides continuous cell-cycle annotations based on the intensity of the fluorescent cell-cycle markers CDT1 and GMNN [1]. Since the FUCCI-U2OS data lack nuclear (DNA) staining, we explored alternative image inputs for the DNA channel. SubCell predictions of pseudotime and cell cycle stages were most accurate when a nuclear pseudo-image was generated from GMNN and CDT1 staining as DNA channel input (mean absolute error of 0.105 on a 0-to-1 cell-cycle scale), rather than leaving the DNA channel empty or filling it with a binary nuclear mask (Fig. 2K, L). This shows that cell cycle prediction accuracy benefits from nuclear texture information.

Taken together, the benchmark results demonstrate that SubCell captures rich protein localization and morphological information across organisms, resolutions, and imaging modalities without fine-tuning. Across all benchmarks, SubCell matched or outperformed SubCell-P (Fig. 2C), with particularly strong gains for morphology, demonstrating that the multi-task objective in SubCell captures both protein-specific and morphological features without a significant trade-off. We therefore focus on SubCell in subsequent analyses (for more test results on SubCell-P, see Methods). In a proof-of-concept experiment, we observed that SubCell’s morphological representation may extend to profiling cells in tissue images (Supplementary Fig. S4).

### Biologically meaningful representations

Having established SubCell’s performance on supervised benchmarking tasks, we next examined whether SubCell’s embeddings extract meaningful phenotypic information that could be used for assessing molecular and biological mechanisms. We first explored whether SubCell embeddings capture cell identity: whether SubCell’s representation of cell identity clusters according to their transcriptomic similarity (as in [19]). Indeed, clustering SubCell embeddings across 36 cell lines closely resembled a clustering based on mRNA sequencing of the same cell lines (Fig. 3A). Furthermore, aggregating and clustering SubCell embeddings by protein localization closely matched hierarchical groupings of 26 organelles (Fig. 3B, Supplementary Table S2), confirming that SubCell’s representations encode subcellular spatial organization. Comparing these similarity measurements to those obtained from state-of-the-art image representation models, SubCell outperformed or matched all models in representing cell line characteristics and self-supervised models in subcellular architecture (Fig. 3C). Since protein function is dependent on subcellular localization [6, 8], we further tested whether SubCell’s image embeddings also capture functional relationships among proteins, a task beyond human visual assessment. SubCell outperformed all models in predicting protein groupings at different hierarchical levels in Reactome [32] and Gene Ontology (GO) [33] databases (Fig. 3D). The modest absolute performance across all models reflects the inherent difficulty of inferring functional relationships from images, as well as the sparsity and cell-type-agnostic nature of manually annotated databases such as Reactome or GO. We conclude that SubCell’s image representations encode meaningful biological features, including cell-line identity, protein localization hierarchies, and functional protein relationships.

**Fig. 3.**
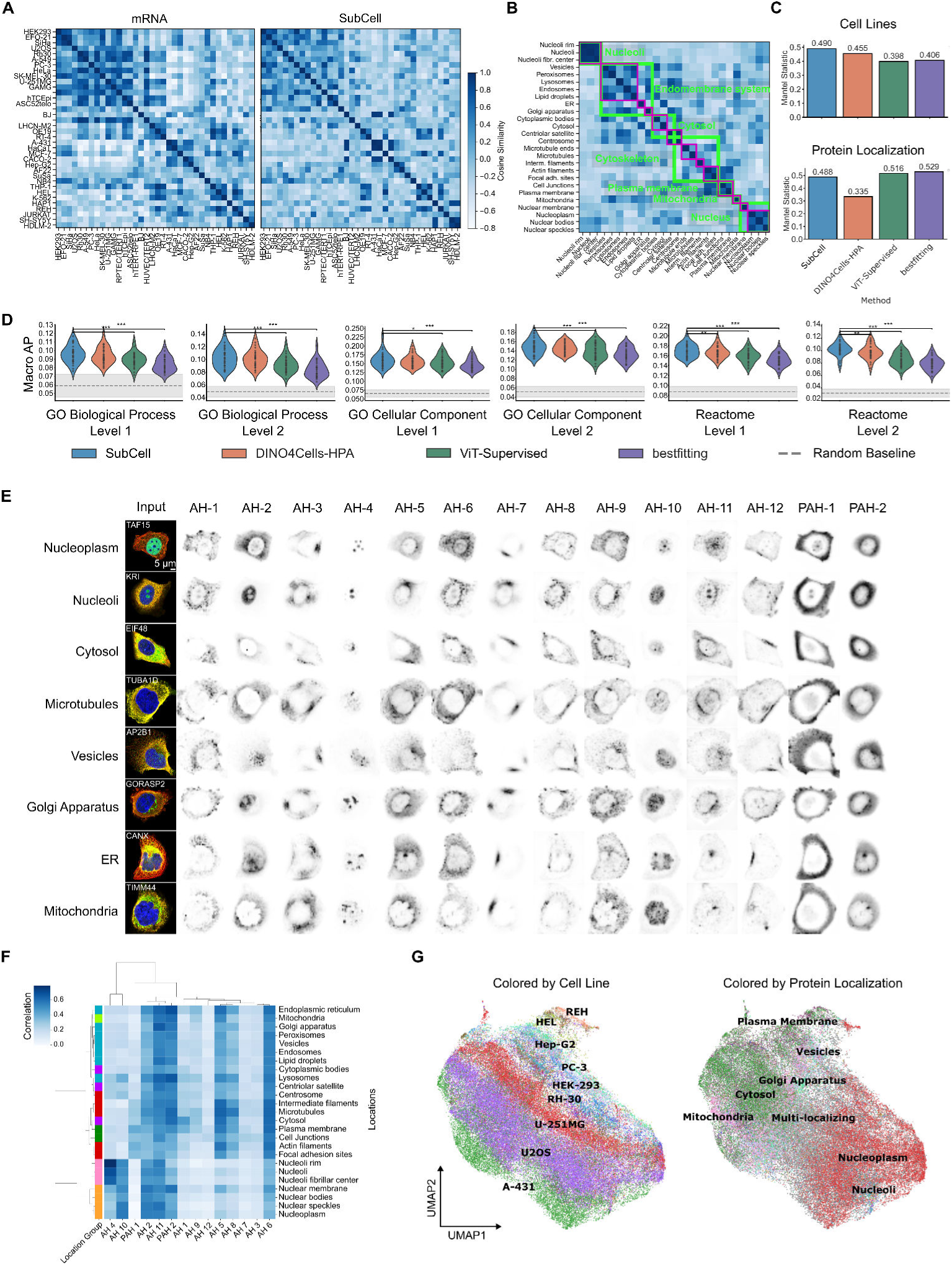
Feature validation of SubCell. (A) Profiling of the cell lines using bulk mRNA levels (left) and aggregated SubCell features (right). The matrices show the cosine similarity between cell lines based on mRNA levels or model features. The order of rows and columns follows the groups determined by the hierarchical clustering of the mRNA similarities. (B) Cosine similarity among the SubCell features aggregated by localization categories. Clusters highlighted in green are the localizations in seven major categories, and the clusters highlighted in magenta are the minor categories. (C) Mantel statistic for the correlation between cell-line-aggregated feature profiles from SubCell embeddings and the associated bulk mRNA sequencing profiles (top), and between subcellular localization patterns from SubCell and the hierarchical profiles annotated by experts (bottom). (D) Violin plots showing the macro-AP score comparison of the classification performance of the models across different known Level 1 and Level 2 designations denote the hierarchical granularity of the dataset annotations, with Level 1 corresponding to higher-level groupings and Level 2 being more granular (see Methods). The results were obtained by performing ten-fold cross-validation three times to establish statistical significance, with p-values computed using the Wilcoxon test to assess whether a method performs better than SubCell. Random baselines were established by training the classifiers on normally distributed random features. (E) Examples of fluorescent images and the attention captured by the attention heads and pooled attention heads in the SubCell model for different protein localizations in HPA (Examples for other datasets are shown in Supplementary Fig. S5). The twelve attention heads of the vision transformers are marked by the prefix AH-, and the pooled attention heads by the prefix PAH-. (F) A cluster map showing the aggregated Spearman’s correlation profiles of the attention heads with the protein channel for different localization categories as defined by expert annotations in C. The order of rows and columns is set by hierarchical clustering of the correlation profiles. The positions of major location groups are shown in different colors on the left side of the plot. (G) We compute correlation profiles with attention maps for each channel of each cell in the HPA, then concatenate them to obtain a single correlation profile per cell. Correlation profiles are aggregated across all cells within each FOV and plotted using UMAP. The colors represent cell lines (left) and annotated protein localization groups (right).

To understand how SubCell encodes these biological features, we examined the attention maps of individual transformer attention heads in the SubCell model (see Methods). Most attention heads were directed to specific subcellular regions, many resembling organelle patterns, such as nucleoli (AH-4), microtubules (AH-8), the endoplasmic reticulum (PAH-2), the plasma membrane (AH-12), or the nucleoplasm (AH-10) (Fig. 3E, Other data sets: Supplementary Fig. S5). The attention to biologically recognizable structures without explicit supervision suggests that SubCell internally decomposes cells into organelle-level features, owing to multi-task training objectives. Notably, when multiple cells are present in an image, SubCell attention predominantly focuses on the central or “complete” cell (as seen in the PAH-1 and PAH-2 attention heads in the OpenCell and JUMP examples, Supplementary Fig. S5), further highlighting the generalizability of SubCell without requiring any preprocessing (i.e., cell segmentation).

To quantify this, we computed correlation profiles between image channels with the attention maps from each head, aggregated these profiles by protein localization category, and performed hierarchical clustering (Fig. 3F; see Methods). The resulting cluster map recapitulated organelle hierarchy similar to Fig. 3B, confirming that the learned attention patterns reflect biologically meaningful subcellular compartmentalization. UMAP visualization of these attention-correlation profiles across the full HPA dataset revealed that the major axes of variation aligned with protein localization (vertical) and cell type (horizontal) (Fig. 3F). We further validated the orthogonality of protein localization and cell type in SubCell’s embedding space by statistically analyzing the overlap between the most discriminative directions for protein localization and cell lines in the correlation profiles (see Methods). The discriminative directions were statistically indistinguishable from two randomly oriented sets of directions (overlap = 0.180, 95% CI [0.167, 0.205], p = 0.480) with a mean angle of 68.5° ± 15.3° (90° for perfect orthogonality), indicating that SubCell encodes the two biological factors mostly independently. Taken together, these analyses demonstrate that SubCell’s attention mechanisms effectively disentangle morphological variation across cell lines from subcellular localization patterns.

### Drug-response profiling

Since SubCell encodes biologically meaningful features of protein localization, cell morphology, and protein associations, we next investigated whether SubCell can detect changes in these features induced by external perturbations, representing a key application in large-scale drug discovery and functional genomics.

First, we investigated SubCell’s ability to quantify drug effects using the CM4AI dataset of triple-negative breast cancer cells treated with the anti-cancer drugs Vorinostat or Paclitaxel [34], which comprises images of 454 proteins acquired in a manner similar to HPA images. SubCell classified drug treatments considerably better than the bestfitting and DINO models (Fig. 4A). The improved classification likely resulted from the distinguishable changes caused by drug treatments in the SubCell embedding space (Fig. 4B). To systematically assess SubCell’s utility for discovering drug-induced changes, we compared classical analysis of protein abundance changes (single-cell image intensity) with SubCell embedding-based changes (Fig. 4C; Supplementary Fig. S6). This revealed that SubCell detected changes undetectable by expression-based analysis alone (Fig. 4C), including changes in protein localization without corresponding changes in abundance (Fig. 4D, RNF20 and NSD1). Complementary analysis of SubCell embedding shifts and intensity changes allowed stratification of drug effects into abundance- and localization-based responses. This investigation shows clear proof-of-principle that SubCell readily generalizes to perturbation testing because its learned embedding space captures drug-induced changes in protein abundance, localization, and cellular morphology, enabling a richer characterization of perturbation effects than intensity-based analysis alone.

**Fig. 4.**
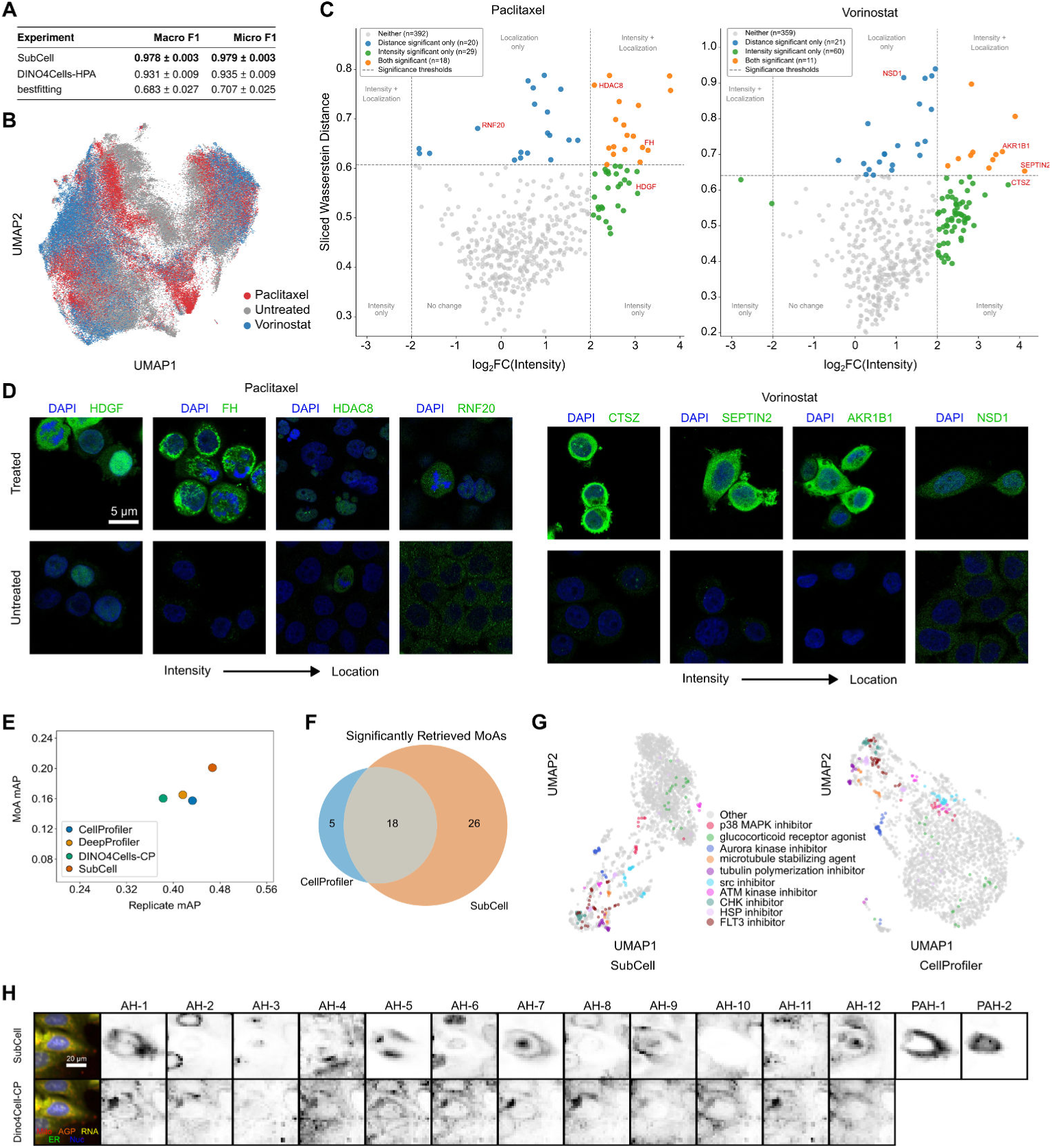
Evaluation of morphology predictions in datasets of drug-perturbed cells. (A) Exploration of drug prediction accuracy based on SubCell embeddings of the CM4AI dataset. Results of ten-fold cross-validation for drug prediction using single-cell features extracted from different models (mean and standard deviation are shown with ±, and the best performance for each metric is highlighted in bold). (B) UMAP of SubCell embeddings of the CM4AI dataset. Each dot represents the embedding of an individual cell. (C) 2D volcano plot comparing intensity differences (*x* -axis) and shifts in SubCell embedding space (*y* -axis) (sliced Wasserstein distance between single-cell embedding distributions) for treated versus untreated cells; individual volcano plots are shown in Supplementary Fig. 6. Each point represents one protein. Proteins are colored by joint significance: grey, neither; blue, embedding only; green, intensity only; orange, both. Intensity significance: | log_2_ FC| *>* 2. Embedding significance: SWD exceeding *d̄* + 1.5*σ*, where *d̄* and *σ* are the mean and standard deviation across all proteins. Statistical significance was assessed by the Mann-Whitney U test with BH and Bonferroni correction for intensity and embedding, respectively (see Supplementary Fig. 6 and Methods). Example proteins explored in (D) are annotated in red. (D) Example proteins revealed to change in intensity only, in both intensity and localization, or in localization only, upon drug perturbation. (E) The mAP non-replicate values for replicate and MoA retrieval for each model on U2OS cells (see Methods). (F) Venn diagram of the number of MoAs retrieved for SubCell and CellProfiler. Significance as determined by permutation testing, with p-value (*p <* 0.05)(see Methods). (G) UMAP visualizations of the well-level profiles for each SubCell and CellProfiler. The ten MoAs that achieved the highest mAP among all models are colored, and the rest shown in grey. (H) Visualization of attention heads and pooled attention modules captured by SubCell and DINO4Cells-CP on JUMP crop of Aminopurvalanol-A perturbed cell. Channels are colored as follows: Mitochondria, red; AGP, orange; RNA, yellow; ER, green; Nucleus, blue.

Next, we examined whether SubCell’s drug-response representations generalize to Cell Painting assays, which have become the dominant morphological profiling assays in pharmaceutical screening [12]. In this assay, cells are stained with six multiplexed dyes labeling general cellular compartments (nucleus, ER, nucleoli, cytoskeleton, Golgi, and mitochondria) rather than individual proteins. Unlike the high-resolution scanning confocal microscopy used in HPA, Cell Painting typically employs lower-resolution fluorescence microscopy. Using a publicly available Cell Painting dataset (JUMP-Pilot, Fig. 1E) [12], we compared SubCell against three state-of-the-art single-cell methods for interpreting Cell Painting data: CellProfiler [3], the gold standard tool for computing hand-crafted pixel-level features; DeepProfiler [35], a weakly-supervised model trained to predict treatment labels; and DINO4Cells-CP [19], a self-supervised model trained on Cell Painting data.

SubCell outperformed all three methods on the standard tasks used to assess Cell Painting images: replicate retrieval (identifying wells treated with the same compound across plates), and mechanism-of-action (MoA) retrieval (identifying distinct compounds affecting the same biological mechanism) (U2OS cells: Fig. 4E; A549 cells: Supplementary Fig. S7A). SubCell also retrieved more statistically significant MoA classes than any other method (U2OS cells: Fig. 4F; A549 cells: Supplementary Fig. S7B). For example, for U2OS cells, SubCell identified 26 MoAs that CellProfiler (the second-best method) missed, while CellProfiler retrieved only 5 MoAs missed by SubCell (Fig. 4F). UMAP projections of JUMP-Pilot embeddings showed that SubCell groups drug responses by MoA label comparably to CellProfiler (Fig. 4G). Quantitative evaluation revealed that SubCell achieved the most cohesive MoA groupings (measured by mAP per category; Supplementary Fig. S8). This strong performance is particularly notable because, unlike DINO4Cells-CP and DeepProfiler, SubCell was never trained on Cell Painting data and was therefore naive to the cellular reference markers used. Qualitatively, we observe that SubCell attention maps show more intricate and diverse subcellular patterns than DINO4Cells-CP on Cell Painting data (Fig. 4H, I Supplementary Fig. S9), suggesting that SubCell’s training on proteomewide pattern diversity enables generalizable recognition of subcellular features. We conclude that SubCell captures morphological signatures of drug perturbations across imaging modalities and resolutions without any domain-specific training, positioning it as a versatile tool for pharmaceutical screening.

### Vision-based multiscale cell map

Having established that SubCell captures biologically meaningful signatures of cellular morphology and protein localization across cell types and perturbations, we next explored whether SubCell can resolve subcellular organization beyond what is currently annotated and reveal new protein associations. HPA annotations for protein localization consist of 35 subcellular compartments, though this is still an incomplete list. Additional known and unknown compartments, condensates, or protein complexes may be detectable in images, beyond what humans can easily annotate. Previously, we resolved such fine structures by integrating image embeddings with protein-protein interaction data in multiscale integrated cell maps (MuSIC) [36–38]. This approach was limited to fewer than 5,000 proteins due to the sparsity of protein–protein interaction data. Given that SubCell representations are not constrained to predefined categories, unlike the supervised models used in MuSIC, and that SubCell outperforms supervised models at capturing protein associations (Fig. 3D), we hypothesized that SubCell could enable the construction of a proteome-wide cell map from image data alone.

We embedded all HPA images for the U2OS cell line (9,543 proteins) with SubCell and applied hierarchical Leiden clustering [39] at increasing resolutions (Fig. 5A) to construct a hierarchical tree of subcellular protein localization (Fig. 5B; Supplementary Table S3). Gene Ontology (GO) enrichment analysis of the resulting clusters identified them as subcellular compartments at coarse resolutions and protein complexes at fine resolutions. For example, the map identified a cytosolic ribosome cluster (6.5), which at finer resolution split into three sub-clusters (7.12, 7.13, 7.14; Fig. 5B). Protein-protein interaction data from STRING [40] confirmed that cluster 7.12 represents the ribosomal protein complex (Fig. 5C), suggesting that at the finest resolution, the map distinguishes co-localizing protein complexes from more loosely associated proteins.

**Fig. 5.**
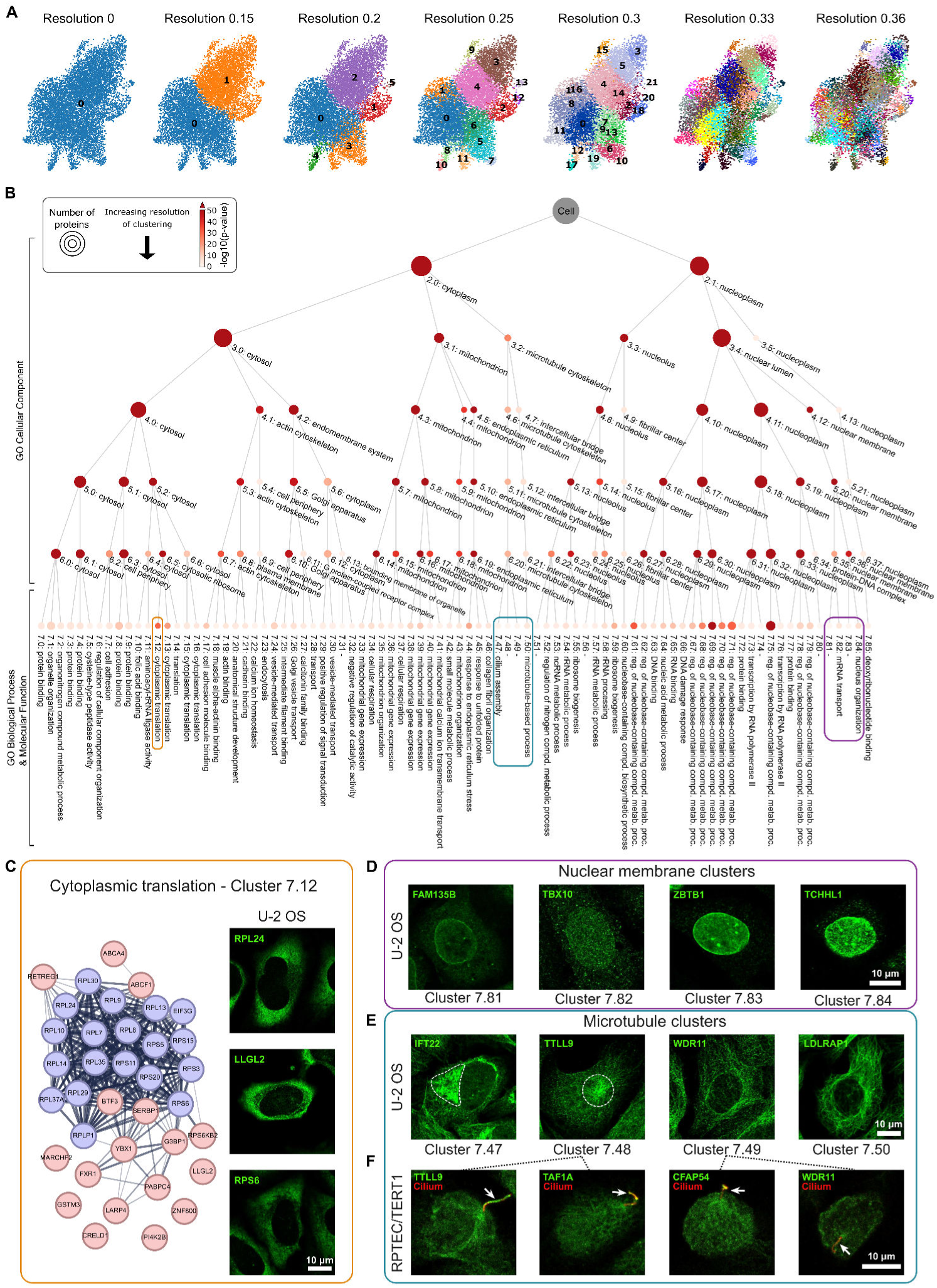
A vision-based multiscale map of cellular protein architecture in U2OS cells. (A) HPA protein embedding UMAPs showing the resulting clusters of Leiden subclustering at increasing resolutions. (B) Multiscale hierarchical map obtained by subclustering of the 9543 protein representations of HPA images. Clusters and subclusters were annotated using GO Cellular Compartment (levels 1-6) or GO Biological Process/Molecular Function (level 7) enrichment analyses; the most significant term is shown. Each node is a cluster from A at the given subclustering / resolution level. The node size indicates the number of proteins in the cluster; the node color shows the − log_10_(p-value) of the functional enrichment. (C) STRING network of the proteins contained in cluster 7.59, annotated as cytoplasmic translation (left, blue nodes are part of the GO term, red nodes not part of the GO term). Exemplary HPA immunofluorescence microscopy images of proteins contained in the cluster showing the distinct staining pattern (right). (D and E) Representative images for the four different nuclear membrane-related clusters (D) and the four different microtubule-related clusters (E) displaying the different staining patterns in the respective clusters. (F) Staining of proteins from the uncharacterized microtubule clusters (E) in ciliated kidney cells (RPTEC/TERT1), confirming that these represent cilium-localizing proteins.

In another example of SubCell detecting protein complexes, the multiscale map also resolved functional groups related to the nuclear envelope. Two major nuclear envelope-associated clusters (6.35 and 6.36) each split into two biologically distinct subclusters (Fig. 5D; Supplementary Table S3). Within cluster 6.36, one subcluster (7.84) contained nuclear envelope structural proteins, while the other (7.83) was enriched for nuclear lamina components. Within cluster 6.35, one subcluster (7.82) comprised proteins involved in mRNA processing, splicing, and export, while the other (7.81) contained proteins with nuclear export and nuclear enzymatic functions. This separation of transport-associated (6.35) from architectural (6.36) proteins suggests that SubCell embeddings enable the distinction between static and dynamic localization patterns.

The map also yielded unexpected biological predictions. Microtubule-localizing proteins (cluster 3.2) separated into two visually distinct classes already at coarse resolution: continuous (cluster 4.7) and spotty microtubule stainings (cluster 4.6) (Fig. 5B, E). The continuous-pattern cluster represented canonical microtubule-associated proteins and did not subdivide further (4.7, 5.12, 6.21, 7.50). By contrast, the spotty-pattern cluster resolved into three subclusters at highest resolution (7.47, 7.48, 7.49), one of which (7.47) was enriched for ”cilia assembly” proteins while the other two clusters (7.48, 7.49) were not GO-enriched but contained ciliary proteins (e.g., TTLL9 and NEK8 [41] in 7.48, WDR11 and CFAP54 [42] in 7.49; Fig. 5B, E; Supplementary Table S3).

Since these clusters converge on ciliary biology, we further explored these proteins to see if the SubCell hierarchical map had detected novel protein localizations or functions. The cilium is a hair-like membrane protrusion that serves as a motility apparatus (motile cilia) or a sensory antenna (primary cilia) [43]. Although cilia have never been observed in U2OS cells, ciliary proteins are expressed in this cell line and are known to be located in non-ciliary compartments [44]. We noticed that each of the three subclusters displayed staining patterns matching specific routes of ciliary protein trafficking (Fig. 5E): Some proteins in cluster 7.47 showed an additional Golgi staining pattern, hinting at the known vesicle-based transport of ciliary membrane proteins from the Golgi to cilia [43]; Some proteins in cluster 7.48 showed an additional staining pattern resembling pericentriolar material or centriolar satellites, known to serve as transit stations for ciliary proteins and to regulate ciliogenesis [45, 46]; cluster 7.49 showed spots of equal intensity along microtubules, suggestive of microtubule-based cargo transport that underlies intra-ciliary transport [43]. Consistent with these interpretations, cluster 7.47 contained proteins that mediate transport to cilia or are present in ciliary membranes (ARL13B [43], IFT22 [47], OCRL [48], PIK3R4 [49]) and cluster 7.48 contained NEK8, involved in ciliary transport and biogenesis, as well as under-characterized ciliary proteins TTLL9 [50] and TEKT2 [51] (https://cilia.pro database: [52]). Cluster 7.49 contained additional, less-studied proteins, such as WDR11 [53], thought to modulate ciliogenesis and ciliary signaling.

We hypothesized that proteins not previously known to localize to cilia but grouped into these ciliary clusters would localize to cilia in a ciliated cell type. To test this, we stained kidney cells, featuring primary cilia, for candidate proteins within a parallel study of the ciliary proteome [44]. This confirmed localization to primary cilia for predicted proteins: TAF1A (no prior link to cilia), PIK3CD (linked to cilia only through disease association [54, 55]), CFAP54 (reported only in motile but not primary cilia [56]), TTLL9 (implicated in motile cilia function but not revealed in cilia [50]), as well as an established cilium-localizing proteins (WDR11 [53]) [52] (Fig. 5F). These results demonstrate that SubCell can detect functional protein similarities from staining patterns in microscopy images alone, beyond what humans can realistically annotate. We conclude that SubCell can suggest protein function in cellular contexts not visible in the training data, such as inferring ciliary protein localization from non-ciliated cells.

### Combining Protein Localization and Sequence Information

Predictions of protein function from SubCell are driven purely from images and therefore may not reflect properties encoded in the amino acid sequence, such as biochemical or structural features. Protein Language Models (pLMs), by contrast, are trained on sequences across the tree of life, and their embeddings have become powerful prediction tools for protein function based on biochemical and structural properties[57–60]. However, pLM embeddings often fail to fully predict subcellular localization [61], which is crucial for understanding context-dependent protein function [6]. Comparing SubCell embeddings of proteins from HPA U2OS cell images with embeddings of the same proteins’ sequences produced by the well-established pLM ESM-2 [58], we confirmed that SubCell and ESM-2 provide complementary information: hierarchical cluster labels derived in one embedding space did not transfer to the other (Fig. 6A, top two rows; Quantification: Supplementary Fig. S10A). We hypothesized that combining the two modalities would create an information-rich protein embedding with both sequence and localization perspectives, so we integrated them using the MuSIC co-embedding approach [37] (selected over MIRAGE [62], see Methods; Supplementary Fig. S10B). The multimodal embedding recovered clusters from both image-only and sequence-only embedding spaces (Fig. 6A, bottom row; Quantification: Supplementary Fig. S10A), confirming that the co-embedding preserved information from both imaging and sequence.

**Fig. 6.**
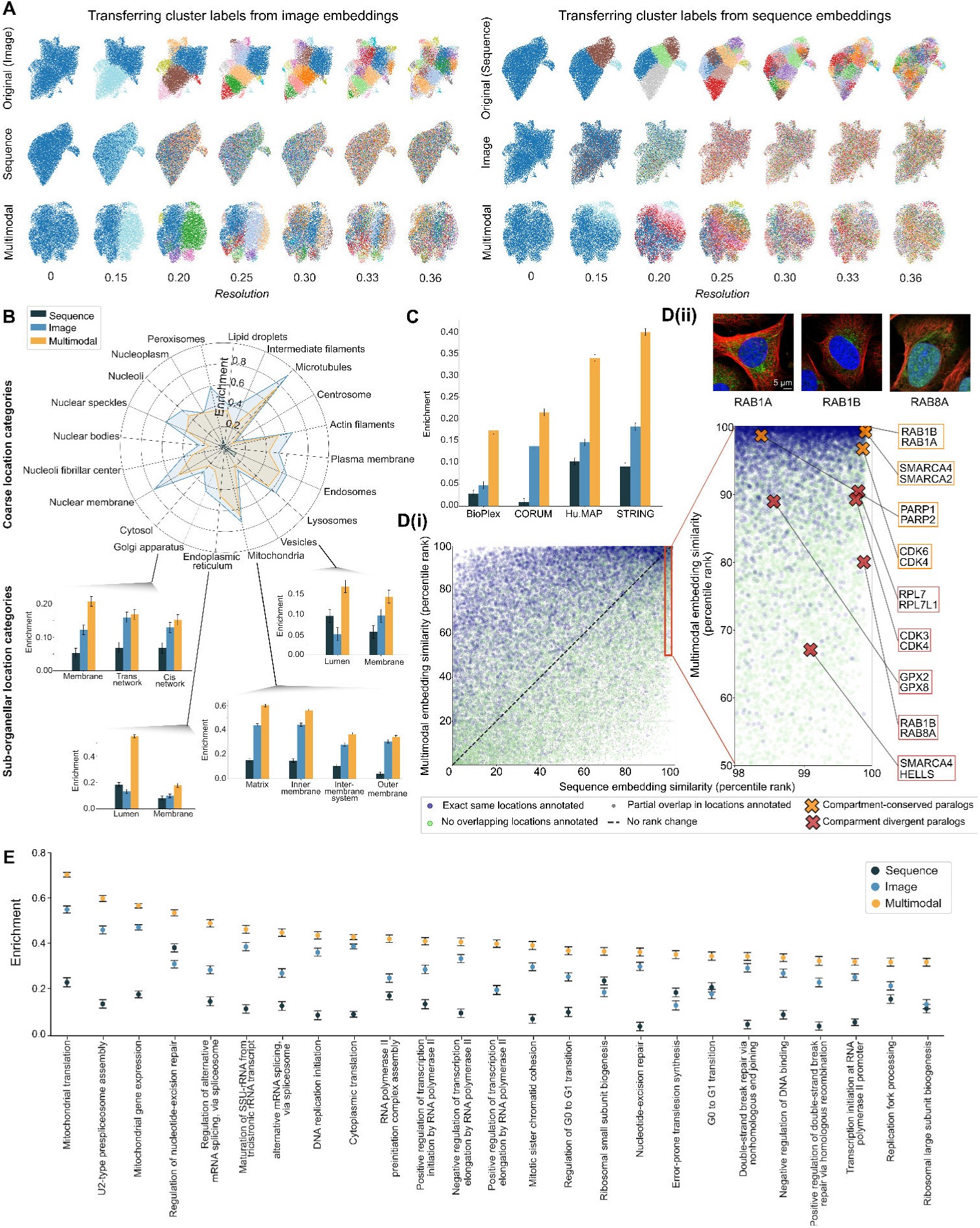
Evaluating the multimodal integration of image and sequence data. (A) Top row: UMAPs of iterative Leiden subclustering of the SubCell embedding space (left) and ESM-2 embedding space (right) on proteins in HPA’s U2OS images; Middle row: The same cluster assignments as in top row, but transferred onto the other embedding space; Bottom row: The same cluster assignments as above but transferred to the multimodal space. (B) Radar plot of enrichment (via approximated Cliff’s Delta throughout this figure) of coarse subcellular localization categories for ESM-2 sequence (navy), SubCell image (light blue), and multimodal (yellow) embeddings. Sub-plots: Enrichment of each embedding space for select sub-organellar localizations (a level below the coarse labels in GO:CC). (C) Similarity enrichment of the sequence embedding (red), image embedding (orange), and multimodal embedding (blue) for interacting pairs as noted in the BioPlex, CORUM, Hu.MAP, and STRING databases. Bar plots show the approximated Cliff’s delta, with error bars indicating the standard deviation. (D) (i) Scatter plot comparing the percentile rank of protein pair cosine similarity in sequence vs. multimodal embedding space. Proteins that have the exact same localization pattern (blue) tend to be closer in multimodal space than sequence space, and those that have non-overlapping localization patterns tend to be farther (green). (ii) Examples of redundant (red) and divergent (orange) paralogs highlighted on the percentile rank comparison plot with the images of RAB1A, RAB1B, and RAB8A displayed as paralog co-localization and divergent localization, respectively. All paralogs are highly similar in sequence space, but divergence is detected in multimodal space. (E) Enrichment measure for gene members in GO biological processes, across sequence, image, and multimodal embedding spaces. Top 25 of 158 enriched biological processes shown. The mean approximated Cliff’s Delta with arrow bars displayed.

Next, we compared the three embeddings based on their ability to resolve known biological relationships of proteins. For localization at the coarse compartment level (HPA annotations), image embeddings performed best, with multimodal embeddings close behind, while sequence embeddings performed poorly, likely because ESM-2 captures homology rather than localization (Fig. 6B, radar plot). At the sub-compartment level (finer GO:CC annotations), sequence features become informative: although sequence alone only sometimes outperforms image, the multimodal embedding consistently outperforms both, sometimes more than additively (e.g., ER lumen; Fig. 6B, panel plots). This indicates that sequence information improves localization resolution specifically at the sub-organellar level, perhaps because distinguishing functionally distinct proteins within the same compartment requires information beyond spatial patterns alone (e.g., due to function-specific motifs found in the sequence). The multimodal embedding also outperformed each unimodal embedding in recovering physical protein interactions from BioPlex [63], CORUM [64], huMAP [65], and STRING [40], again more than additively (Fig. 6C). Across all of these co-localization tasks, though image and sequence performance vary, the integration of the two consistently tracks or exceeds the stronger single modality.

Although pLMs generally perform well on protein function prediction [66], we next asked whether multimodal embeddings resolve functional differences between related proteins that ESM-2 alone cannot distinguish. Paralogs (proteins arising from evolutionary gene duplication) are a natural test case because they often retain high sequence similarity but can diverge in localization and function, limiting what sequence alone can reveal. We found that co-localizing proteins had higher pairwise similarity scores with the multimodal approach compared to sequence-only (Fig. 6D(i)). Given this, we selected examples of location-conserved vs. location-diverged paralogs and showed that they were separated in multimodal but not sequence space (Fig. 6D(ii); Example images for a conserved location (RAB1A vs. RAB1B) and a diverged location (RAB1B vs RAB8A) are shown). Image space alone also failed to capture paralog localization divergence because co-localization of many proteins obscures paralog-specific signals (Supplementary Fig. S10C). These results demonstrate that the multimodal embedding of SubCell and ESM-2 creates a single representation of how gene duplication followed by spatial relocalization drives divergence in protein evolution.

Beyond pairwise relationships, we examined whether the multimodal embedding captures systems-level biological processes. We assessed whether proteins sharing a Gene Ontology Biological Process annotation are more similar to one another than to other proteins in the dataset. Proteins that shared a process annotation were often more enriched in the multimodal space than in either unimodal space (Fig. 6E; remaining in Supplementary Fig. S11), including temporally dynamic processes such as the cell cycle (e.g., G0–G1 transition). These results indicate that the multimodal embedding encodes systems-level cellular operations, suggesting it can serve as a foundation for pathway discovery and functional annotation of uncharacterized proteins.

## Discussion

SubCell is a generalizable vision foundation model trained on subcellular protein localization patterns from the Human Protein Atlas using a multi-task learning framework that combines reconstruction, cell-specific, and protein-specific objectives. SubCell outperforms state-of-the-art models across a diverse range of tasks in single-cell biology, including localization prediction and morphological phenotyping. It readily generalizes to datasets with different cell types, reference markers, and resolutions without fine-tuning. Together, these properties make SubCell a versatile analytical tool for single-cell fluorescence microscopy, with demonstrated applications across drug profiling, mapping cellular architecture, and characterizing protein function.

The complexity of dynamic subcellular protein localization is often oversimplified into hand-crafted sets of categorical compartments. Deep-learned continuous representations of cellular morphology are now well established [15, 19, 35], but continuous representations of protein localization have remained out of reach. SubCell closes this gap by providing continuous, learned representations of protein localization as well as morphology, thereby unlocking new modes of biological analysis.

SubCell representations enable quantification of how a protein’s localization shifts under different cellular conditions, such as chemical or genetic perturbations. We demonstrate this on the CM4AI dataset, where SubCell detected whether a cancer cell was treated with greater accuracy than established methods and identified drug-induced relocalization events that are invisible to common expression-based analyses. Recent studies have demonstrated that mislocalization of pathogenic coding variants is common among human disorders [4], and that technologies for protein relocalization may open up novel therapeutic opportunities [6, 7]. Therefore, SubCell’s ability to detect subtle spatial shifts in localization at scale positions it as a powerful tool for advancing therapeutic research.

SubCell representations enable comparisons of localization patterns across the proteome. Using SubCell embeddings, we constructed a vision-based, proteome-wide multiscale map of subcellular protein architecture. This resource serves as a data-driven alternative or complement to GO for gene, protein, and pathway annotations — one less biased by human-defined localization categories. Using this map, we identified novel ciliary proteins based on cluster similarities and confirmed them by subsequent staining. Looking forward, we envision future SubCell maps that capture the rewiring of cellular organization and pathway interactions in chemically or genetically perturbed cells. Integrating our map with transcriptome-based perturbation models could improve spatial contextualization and facilitate causal reasoning toward a virtual cell model [67].

SubCell representations enable the integration of complex continuous spatial information with non-imaging modalities. Combining SubCell spatial protein embeddings with ESM-2 sequence embeddings led to superior performance in predicting protein sub-organellar localization, protein-protein interaction, functional divergence of paralogs, and biological process tasks. Neither SubCell nor ESM-2 alone matches the performance of the joint embedding, emphasizing the biologically meaningful synergy between localization and sequence information. Our results suggest that spatial context is necessary to move from a molecular to a systems-level understanding of protein function in the cell, and that the next generation of PLMs should be able to reason over sequence, structure, function, and localization.

In addition to these protein-specific applications, SubCell serves as a generalizable feature extractor for single-cell images across microscopy methods, resolutions, cell types, drug perturbations, cells in tissue contexts, and even species. Most notably, it remains robust to datasets with unseen channel-marker combinations. SubCell’s flexibility likely stems from training on the HPA and its proteome-wide representation of localization patterns: protein-specific supervision encouraged the model to encode the diversity of subcellular patterns in a structured latent space rather than relying on channel identity. This organization is reflected both in SubCell’s strong performance across downstream tasks and in its visually interpretable attention maps. Consequently, SubCell can meaningfully represent images exhibiting a wide range of subcellular patterns, regardless of the marker used to visualize them. We demonstrate this on the JUMP-Pilot Cell Painting dataset: by providing Cell Painting stains as input to SubCell’s protein channel, SubCell profiles perturbed morphologies more effectively than supervised models specifically tailored to Cell Painting data, despite SubCell never having seen these stainings during training.

The prevailing trajectory in machine learning emphasizes scaling — larger datasets and increasingly large models. A recent study introduced CHAMMI-75, a large-scale microscopy training resource that integrates diverse image datasets spanning different channel numbers and marker combinations [68]. The same study also presented MorphEm, a model trained on CHAMMI-75 that employs a flexible channel-adaptive architecture to accommodate channel heterogeneity. SubCell takes a different approach: instead of training across many heterogeneous datasets, we train on a single, high-resolution dataset encompassing a broad and balanced diversity of protein patterns. Thereby, we focus model training on biological instead of technical variation. SubCell outperforms MorphEm and other feature extractors on four of six tasks in CHAMMI-75’s own benchmark evaluations; the remaining two tasks fell outside SubCell’s supported input regime, involving brightfield and 14-channel imaging. We conclude that training on high-quality data of protein localization patterns, paired with pattern-specific supervision, provides a powerful alternative to sheer data scale for training robust biological vision foundation models. A natural next step is to integrate channel-adaptive architectures with SubCell’s protein-pattern-driven training, unifying SubCell’s channel-specific variants into a single model to enable more direct comparisons across datasets.

SubCell’s generalizability across imaging modalities, cell types, and reference markers enables broad utility for the research community. The model weights and source code are freely available, and the accompanying repository provides a flexible framework for adapting SubCell to new datasets through automatic channel-variant selection. SubCell requires only a DNA channel as minimal input, and reference markers not included in the HPA can be incorporated through the protein channel. We also provide tutorials, a ready-to-use online application, and an interactive version of the proteome-wide hierarchical cellular map. By making the spatial proteome computationally accessible at single-cell resolution, SubCell enables systematic discovery of protein function from microscopy images and lays a foundation for integrating spatial and molecular information toward a data-driven understanding of the cell.

## Methods

### Datasets

We used the HPAv23 dataset to train SubCell and evaluated it using the other datasets. The dataset descriptions are as follows, and details on their sizes and pixel resolutions are provided in Supplementary Table S4.

### Human Protein Atlas Subcellular Dataset

The HPA subcellular dataset is a collection of immunofluorescence images that encode the expression and spatiotemporal distribution of 13,141 genes across 37 cell lines [8]. The cell lines represent a broad band of tissues, cell types, age, and sex: e.g., U2OS is an osteosarcoma cell line derived from the tibia of a 15-year-old female patient; ASC52telo is a primary mesenchymal stem cell line derived from the adipose tissue of a female donor; RPTEC/TERT1 is a primary kidney proximal tubular cell line derived from the kidney cortex of a male donor; SH-SY5Y is a neuroblastoma cell line derived from a bone metastasis in a 4-year old female patient; AF22 is an induced-pluripotent-stem-cell-derived neuroepithelial stem cell line from a 36-year old female patient; CACO-2 is a colon-derived adenocarcinoma cell line derived from a 72-year old male; U-251MG is a glioblastoma cell line derived from the parietal lob of the brain of a 75-year old male. The images were generated by staining the cells with DAPI, a fluorescent dye that labels DNA/nucleus, and antibodies that label the endoplasmic reticulum (ER), microtubules (MT), and the protein of interest (Protein). The protein localization in the dataset is categorized into 35 organelles and fine subcellular structures. Version v23 of the image dataset was used. Single cells were extracted from the images using the segmentation masks generated by the recommended HPA-Cell-Segmentation method [27]. The segmentation masks were processed to remove nuclei touching the border and those that were too small, and then to merge detections that were too close to each other. Altogether, 1,138,378 cells were extracted from the images and used as the model’s dataset.

The dataset was split into training, validation, and test sets based on the antibodies, with an approximately 7:1:2 ratio. A multilabel stratification strategy [69, 70] was used to ensure a similar multilabel distribution between the sets. Further details on the data distribution across the test, train, and validation sets can be found in the GitHub repository (https://github.com/CellProfiling/subcell-analysis).

HPA antibodies intentionally target as many protein isoforms of a gene as possible, so when we refer to proteins, we more specifically mean the protein products of a given gene. This is why all annotations throughout the paper are on the gene (rather than protein) level. For the same reason, all analyses are performed at the level of gene identifiers, meaning that protein identifiers in task-specific datasets are converted to gene identifiers before SubCell is evaluated on the task.

#### OpenCell Dataset

The OpenCell dataset comprises live-cell confocal images of 1,310 CRISPR-based, endogenously tagged proteins on the HEK293T cell line. The cells were imaged using a nuclear stain (Hoechst 33342) in conjunction with the protein. The single-cell images were generated by segmenting the nuclei and cropping a 256x256-pixel region around them. Overall, 94,426 cells were obtained. The annotations for protein localization are divided into 18 categories, including multi-localizing proteins. The OpenCell categories were converted to the HPA categories by an expert and are reported in Supplementary Table S2.

#### AllenCell Dataset

We used a subset of the Allen Institute Cell Explorer dataset [10] used in the DINO4Cells paper [19], containing 214,037 human induced pluripotent stem cells (hiPSC) from 25 isogenic cell lines, each containing one fluorescently tagged protein via CRISPR/Cas9 gene editing. The cells were imaged in 3D using fluorescently tagged markers for the nucleus and plasma membrane, as well as the protein. The masks for all the markers were provided within the dataset. For our analysis, we used the masked maximum projection of the channels across the z-plane. The cell cycle stage labels for the cells were annotated by experts and were used to validate the models.

#### Yeast Dataset

This dataset contains fluorescence microscopy images of live Saccharomyces cerevisiae cells expressing a GFP-tagged protein of interest, along with two reference markers: RFP to label the cytoplasm and F-RFP to label both the nucleus and the bud neck via the septin protein CDC11 across 4,100 proteins [28]. Images were acquired using a high-throughput spinning-disc confocal microscope. Two datasets of single-cell image crops are provided: one for subcellular localization and another for cell cycle classification, both available at https://github.com/BooneAndrewsLab/CycleNET. Each dataset was manually annotated, comprising 22 localization classes and 9 cell cycle classes, respectively. Each dataset is split into training and testing sets. For our analyses, we excluded the “Over-segmented” and “Aberrant” categories from the cell cycle dataset, as these reflect segmentation artifacts or empty images rather than biologically meaningful states.

#### U2OS FUCCI Dataset

The dataset comprises endogenously tagged cells expressing two fluorescent proteins, CDT1 and GMNN, which are fused to cell-cycle regulators, enabling monitoring of the cell cycle [1]. Along with these proteins, the cells were also tagged with a protein of interest and a microtubule marker. The cell segmentation was done using the HPA-Cell-Segmentation method [27]. Overall, 357,083 cells were obtained, for a total of 1,166 antibodies. A polar-coordinate pseudotime model was employed to generate a continuous representation of the cell-cycle position, spanning 0 to 1, with 0 corresponding to the beginning of G1 Phase and 1 to the end of M phase. Furthermore, based on the cell cycle estimation, the cells were labeled between three discrete stages: G1, G1S, and G2.

#### CM4AI dataset

The image dataset of drug-treated triple-negative breast cancer cells originates from the Cell Maps for Artificial Intelligence (CM4AI) project, a component of the U.S. National Institute of Health’s (NIH) Functional Genomics Data Generation Project within the Bridge2AI program [34]. We used only the subcellular proteomics dataset, i.e., the immunofluorescence imaging dataset, and of this, only the dataset focusing on the triple-negative breast cancer cell line MDA-MB-468 ([71]). The dataset comprises 63x confocal images of cultured MDA-MB-468 cells treated with the drug Paclitaxel (NSC 125935, Selleckchem S1150), the drug Vorinostat (SAHA, Selleckchem S1047), or untreated. The experimental workflow followed the workflow applied in the Human Protein Atlas (HPA) subcellular section: The cells were seeded in fibronectin-coated glass-bottom 96-well plates (18,000 cells per well), treated with the different treatment conditions, incubated for 24 hours, fixed with 4% paraformaldehyde (PFA) and stained with a nucleus marker (DAPI), a microtubule marker (alpha-Tubulin antibody, Abcam, ab7291, RRID: AB 2241126), an ER marker (Calreticulin antibody, Abcam, ab2908, RRID: AB 303403), and an antibody against a protein of interest from the HPA antibody library. Each well was stained with a different antibody against a protein of interest. For each protein of interest, experiments across the different treatments and staining were performed simultaneously to minimize effects unrelated to the perturbation. The dataset contained the images acquired for 454 different proteins.

#### JUMP-Pilot Dataset

A subset of the JUMP-Pilot dataset (cpg0000-jump-pilot) was used [12]. The complete JUMP-Pilot dataset contains 300 million images of U20S and A549 cells under both chemical and genetic perturbations at both short (24hr) and long (48hr) time points. JUMP-Pilot data, segmentation masks, metadata, and CellProfiler profiles are publicly available at https://cellpainting-gallery.s3.amazonaws.com/index.html#cpg0000-jump-pilot/. A subset of plates was selected by filtering for chemical perturbations and the absence of any antibacterial agent. After filtering, 15 plates remained. BR00116995 contained an unusual number of empty wells and was excluded from our analysis. The layout of treatments across wells was the same for each plate, containing 64 negative control wells and 320 wells with chemical perturbation from one of 302 compounds. We used 77 pre-defined MoA classes [72]. When calculating MoA mAP values, only 192 compounds were used because they shared at least one MoA with another compound.

#### Tissue Dataset

The image dataset of a pancreatic cancer (adenocarcinoma) tissue section was generated as follows. The tissue sample preparation protocol was described before in [73]. Tissue MicroArrays (TMA) were constructed with strategies used in the Human Protein Atlas previously described [74, 75] in Formalin-fixed paraffin-embedded (FFPE) format. In summary, hematoxylin-eosin stained tissue sections from FFPE donor blocks were examined to confirm histology and identify representative regions for sampling into the TMA. Normal tissue was characterized as microscopically normal, predominantly sourced from proximity areas to surgically excised tumors. TMA blocks were created, each containing triplicate 1 mm cores from the different types of normal tissue.

We used a described indirect immunofluorescence staining protocol [76] with a few adjustments. In brief, a TMA FFPE block was sectioned at 4 *µm* thickness. The section was dewaxed and underwent heat-induced epitope retrieval (HIER) treatment using pH 9 EDTA buffer (Immuno Retriever 20× with EDTA, Bio SB, BSB 0030) in a pressure cooker (TintoRetriever, Bio SB, BSB 7087) at 114 − 121 °*C* during 20 minutes. After two ddH2O washes, the material was exposed to a photo-bleaching procedure to reduce tissue autofluorescence; following a blocking step using TNB buffer containing 0.1 M Tris-HCl /0.15 M NaCl/0.5% blocking reagent pH 7.5 (Akoya, SKU FP1020) for 30 min. Primary antibodies targeting alpha Tubulin as an MT marker (Abcam, AB7291, RRID:AB 2241126, 1:400) and calreticulin as an ER marker (Abcam, ab2908, RRID:AB 303403, manually conjugated to CF725 (Biotium, # 92583,) and thereafter used at 1:100) were diluted in antibody diluent containing 0.3 % Triton (Sigma Aldrich, T8787) 1x PBS pH 7.4 and added on the TMA section to be incubated overnight at 4 °C. The next day, the material was washed with TBS-T and blocked using TNB for 30 min. A secondary antibody to label the alpha-Tubulin primary antibody (goat anti-mouse IgG1 Cross-Adsorbed Alexa Fluor 555, ThermoFisher, A-21127, RRID:AB 2535769, 1:1000) and Hoechst 33342 to label DNA (Invitrogen, H3570, 1:5000) was diluted in TNB, and added on the section for 90 min incubation at room temperature. After 3 washes in TBS-T, the stained material was mounted with Fluoromount-G (Thermo Fisher Scientific, 00-4958-02).

A multi-tile scan of a region in the TMA was imaged with a Leica Stellaris 8 confocal microscope equipped with an HC PL APO 63×/1.20 water-immersion CS2 objective (Leica Microsystems, Mannheim, Germany). The settings for each image were Pinhole 1 Airy unit, 16-bit acquisition, 12 z-steps, z-step size of 0.7 *µm*, 1.28× zoom, 2048×2048 pixels field of view, resulting in a pixel size of 0.07 *µm*. Each channel was acquired in a separate series to avoid bleed-through, with only the laser required for the specific channel activated. Excitation wavelengths: 405 *nm* (Hoechst, DNA), 553 *nm* (MT), 729 *nm* (ER). Detectors: HyD S1 was used for imaging the Hoechst (DNA) channel (Spectral range setup: 429 − 495 *nm*), Hyd S3 was used for imaging the Alexa Fluor 555 (MT) channel (Spectral range setup: 558 − 620 *nm*), and HyD R for the CF725 (ER) channel (Spectral range setup: 734 − 757 *nm*).

Each tile z-stack was exported as an .ome.tif image from the LAS-X 3D software integration and maximum intensity projected in python. Nuclei were segmented using a customized version of CellPose segmentation (https://github.com/CellProfiling/ell_code_template/tree/master/examples/cellpose_segmentation) to create single-cell crops (https://github.com/CellProfiling/ell_code_template/tree/master/examples/cell_cropper). Single-cell crops were embedded using MultiT (Reference channels: DNA, MT, ER; Protein channel: MT; https://github.com/CellProfiling/SubCellPortable) and a UMAP was generated from the cell embeddings. k-means clustering (*k* = 6) was applied on the UMAP to distinguish cell types (Supplementary Fig. 3C).

For display of the analyzed image region (Supplementary Fig. 3A), maximum-intensity-projection tile images were merged using FIJI’s [77]. Grid/Collection Stitching function and image coordinates retrieved from the microscope’s xml metadata. To display what cell in the tissue was assigned to which cluster, the colored mask image (Supplementary Fig. 3B) was generated by creating single cell masks only for display in the figure (DNA, MT, and ER channel used as input for Cell-Pose segmentation, https://github.com/CellProfiling/ell_code_template/tree/master/examples/cellpose_segmentation) and coloring them by cluster.

### Model Development

#### Vision Transformers

Vision transformers (ViTs) are based on the transformer architecture [26], originally developed for textual data [78], which utilizes a novel self-attention mechanism. A set of word tokens is fed into multiple attention heads, which convert them into keys, queries, and values to get an attention matrix. In ViT architecture, the input tokens are created by dividing the image into fixed-sized patches similar to the words in a sentence. The transformer architecture effectively captures global relationships across the image without relying on convolutional architectures. We selected the ViT-B model, which has approx. 86.4 million parameters and a patch size of 16x16 pixels, for our experiments.

#### Masked Autoencoders

Masked autoencoders (MAEs) [21, 79] introduced an efficient way to train vision transformers at scale in an unsupervised manner. A random subset of the tokens in an image is masked out, and the remaining tokens are fed into the encoder. After encoding, the masked tokens are reintroduced, and all tokens are fed into a small decoder that reconstructs the original image from them. A reconstruction loss is applied to the masked patches to facilitate learning. After the pre-training, the encoder can be used as a feature extractor for the downstream tasks, and the decoder is discarded. A detailed architecture description is shown in Fig. 1. Since only the non-masked tokens are processed through the encoder, MAEs require less computation than supervised training. The tokens to be removed from the image are chosen randomly. However, since we also had the cell masks available, we explored mask-aware token removal, i.e., removing the tokens present in the cells (Supplementary Fig. S12).

#### Contrastive Learning

We employed contrastive loss to train the models with cell-specific and protein-specific objectives. Contrastive learning is a self-supervised learning paradigm that learns image representations by exploiting similarities and dissimilarities between data points [80]. The loss aims to minimize the distance between the positive pairs and maximize the distance between the negative pairs in the representation space. For the cell-specific objective, the positive pairs in a batch are generated by applying different augmentations to the cell image, and the remaining pairs are treated as negative. For the protein-specific objective, all cells treated with the same antibody are considered positives, and the rest are considered negatives.

#### Attention-based Pooling

Attention-based pooling [81] summarizes the contribution of a feature in a large feature set to form a single feature vector, where each feature in the set is weighed based on its importance. Typically used for multi-instance learning, it provides an explainable way to interpret the importance of each feature. Instead of computing the mean of the token embeddings, we used attention-based pooling weights to get a single feature embedding for the image. Each attention head processes the tokens separately and outputs an aggregated feature vector. We empirically set the number of pooled attention heads to 2 in our experiments and concatenated the outputs to double the encoder’s output feature dimension. We observed improved performance when using the pooling module compared to simple mean pooling. Further, the module also provides an interpretability measure in the form of a token importance map.

#### Multi-task Learning

A multi-task framework was used to learn a diverse set of features for the ViT encoder. To enable effective contrastive learning with the MAE framework, only one of the views is fed to the decoder, similar to [82, 83], and the augmentation strategy was modified. The images were augmented with geometric transformations applied to the view fed into the decoder, as MAEs reconstruct pixel intensities. Meanwhile, the other view was augmented with both geometric and color transformations. The outputs of both views are passed through the attention pooling module, and the contrastive loss is computed based on the resulting feature representations. For the SubCell-P model, both views were augmented with geometric and color transformations.

The geometric augmentations used in our experiments were random vertical and horizontal flips; random affine transformations with a 90-degree rotation, a translation of 0.2, and scale factors of (0.8, 1.2); and random perspective transformations with a distortion scale of 0.25. For the color transformations, we used per-channel transforms as used in Doron et al. [19], with random brightness and contrast scaling of 0.5, a Gaussian blur with a kernel size of 7 and sigma range (0.1, 2.0), and adjusting sharpness with a factor of 2, and random erasing the parts of the channels. We also randomly erased channels with probability 0.25 when experiments involved more than three channels. We used intensity rescaling of the protein channel with a probability of 0.25. We randomly masked the cell crops in a batch with a probability of 0.5 to guide the model to ignore the surrounding cell regions within the crop. This reduces the need for cell segmentation in the analysis, allowing the model to be applied directly to the raw image in a grid-like manner for feature extraction. Furthermore, the augmentation focuses attention on regions within the cell, rather than on the background.

#### Model Training and Inference

All models were trained for 300 epochs using the AdamW optimizer [84] with a base learning rate of 1e-4 and weight decay of 0.05. We used a linear warm-up for the first 5 epochs and decayed the learning rate using a linear decay schedule without restarts. The training batch was modified to ensure an equal distribution of cells from different antibodies and was empirically set to 8 cells per antibody. For training, a 896x896- pixel region around the cell was extracted and resized to 448x448 pixels for feeding into the models.

We conducted an ablation study to determine the optimal cell crop size for inference (shown in Supplementary Fig. S13). We found that the models performed best when using 640x640-pixel crops centered on the cell. The inference was performed on cell crops, using cell masks to ensure a fair comparison with other methods and to ignore the effects of surrounding, incomplete cells (as shown in Supplementary Fig. S12; cells on the borders of the cells are ignored in PAH- attention heads in OpenCell and JUMP examples).

#### Development of the multi-task components of the model

We initiated experiments with masked autoencoders (MAEs) using different mask ratios and object masking ratios (Supplementary Table S5). As shown in the results (Supplementary Table S5), increasing the cell masking ratio while keeping the overall masking ratio constant improved model performance in both tasks. However, we observed that while a higher masking ratio enhances performance in the cell-line classification task, it results in poorer performance in protein localization prediction tasks. Overall, we found that the MAE models lag behind the DINO model in protein localization but are closer to it in the cell-line classification task. We hypothesize that, because MAEs focus on cell morphology in images and protein localization in HPA is independent of cellular morphology, when cells are unperturbed, the MAEs perform poorly on this task.

Subsequently, we trained the models with cell-specific contrastive loss and its combination with the MAEs (Supplementary Table S6). The models trained with MAE and contrastive loss (MAE-CellS) outperformed those trained solely with contrastive loss (CellS) for cell-line classification; however, they lagged in protein localization prediction tasks. The models outperformed those trained solely with MAE on both tasks. We observed that increasing the masking ratio degrades the models’ localization performance, and that the cell masking strategy further degrades it. This might be attributed to differences in the images the encoder sees: in one view, certain parts of the cell are masked (e.g., protein regions), while in others they are not. Still, the best model could not match DINO’s performance.

We performed experiments using the protein-specific contrastive loss framework (Supplementary Table S7). We evaluated the addition of protein-specific loss to the base model (i.e., ViT) and to different top-performing cell-specific and MAE configurations. We found that the model trained with just the protein-specific loss (ProtS) performed best for the protein localization task (micro AP of 0.878), outperforming the DINO (0.863), weakly-supervised (0.869), and bestfitting (0.83) models on the HPA test set, while the model with the MAE, cell-specific, and protein-specific loss (MAE-CellS-ProtS) performed second best (0.872). The detailed architecture of the multitask model is shown in Supplementary Fig. S1. The ViT model trained with cell-specific and protein-specific loss performed best on the Kaggle test set for protein localization but lagged behind in cell-line classification.

Finally, we conducted experiments to evaluate the utility of the pooling module on the two best-performing models from the previous experiments (Supplementary Table S8). The pooling module improved the performance of both models for both tasks in the HPA dataset. Significant improvements were observed in both models across cell classification and protein localization tasks. The multitask pooling model (MAE-CellS-ProtS-Pool), referred to as SubCell, and the protein-specific pooling model (ProtS-Pool), referred to as SubCell-P, were selected as the final models for further experiments.

#### Classification based on SubCell features

For all supervised learning tasks, i.e., localization prediction and cell cycle stage prediction, we trained a multi-layer perceptron (MLP) classifier on features extracted from the models and generated predictions. We used the same three-layer classifier architecture as Doron et al. [19] and trained the classifiers with focal loss to address class imbalance in the dataset. The classifiers were trained on the features extracted from the same images used to train and validate the encoders. We trained the models for 200 epochs using the AdamW optimizer with a learning rate of 1e-3, and reduced the learning rate by 0.5 if the validation mAP didn’t improve for more than four epochs. The training is stopped if no improvement is observed for 20 consecutive epochs. We trained ten classifiers with different seeds for each task and reported the mean and standard deviation of the metrics.

### Evaluation on HPA datasets

We trained two sets of classifiers for the localization classification task. The first set comprised the 19 categories specified in the Kaggle challenge, while the second set encompassed a broader range of 31 categories. This approach enabled us to perform a fair comparison with the bestfitting and DINO4Cells-HPA models, further demonstrating our models’ capability to encode more comprehensive localization information compared to previous methods. The results of the classifier trained with 31 classes are shown in Supplementary Table S9. We evaluated the performance of the models at the field-of-view (FOV) level on the HPA test set and at the single-cell level on the hidden test set of the Kaggle single-cell classification challenge. We averaged the predictions across cells within an FOV to report FOV-level classification results. We reported micro- and macro-average precision (AP) as classification metrics and used multilabel ranking average precision (MLRAP) and coverage error to evaluate multilabel performance on the test sets. MLRAP evaluates how well a model ranks predicted labels by measuring the proportion of relevant labels ranked higher than irrelevant ones, while coverage error computes the average number of labels to be included in the prediction such that all true labels are predicted.

### Evaluation on the OpenCell dataset

The cell features were extracted using the DNA-Protein variant of SubCell. We evaluated the features on the original-resolution images and the resized images, which were matched to the pixel size of HPA images cropped to 640x640 pixels, and found that the features extracted from the resized images performed better. We filter out multi-localizing proteins and aggregate single-cell features to create FOV-level profiles within the dataset. We employed the K-Means clustering algorithm for unsupervised clustering of the features and evaluated the agreement between the resulting clusters and the OpenCell annotation labels to compare the feature representations across the models. We reduced the number of features by keeping the top 100 principal components to minimize the noise in clustering. The number of clusters was set to the number of localization categories in the data. The clustering labels were then compared with the ground truth labels using the adjusted Rand Index [85], which assesses the similarity in cluster assignments, and V-Measure [86], which assesses the homogeneity and completeness of the assignments. We evaluated performance using clustering rather than classification for two main reasons: the number of available samples (approximately 6,300 FoVs) was insufficient to train classifiers reliably, and we aimed to maintain consistency with the analysis performed in CytoSelf [22].

### Evaluation on the AllenCell dataset

The cell features were extracted using the DNA-Protein and MT-DNA-Protein variants of SubCell and SubCell-P. Different channel combinations and image normalization techniques were evaluated (Supplementary Table S10). The images were resized to match HPA pixel size and cropped to 640x640 pixels for inference. We used the same training and test sets as those in the DINO4Cells paper [19]. Additionally, a validation set was extracted from the training set to choose the optimal model weights for the MLP classifiers during training. Ten different classifiers were trained with different random initializations of the model weights, and the mean and standard deviations were reported.

### Evaluation on the FUCCI dataset

The cell features were extracted using the MT-DNA-Protein variant of SubCell and SubCell-P. The images were resized to match HPA pixel size and cropped to 640x640 pixels for inference. Since the DNA channel is not present in the images, three different combinations of the CDT1 and GMNN channels were used to substitute for the channel in the experiments, i.e., passing an empty image, passing a nuclear mask with a constant value, and passing the nuclear channel as the average of the cell cycle markers (CDT1 and GMNN). The nuclear mask was obtained by simply applying Otsu’s thresholding to the mean image of the CDT1 and GMNN channels. The dataset was split into 10 folds stratified by the antibody ID, and cross-validation performance was recorded. For each fold, models were trained 10 times with different random seeds to account for prediction variability. For the regression task, i.e., predicting pseudotime from cell features, we used deep evidential regression training [87], which learns to account for both aleatoric and epistemic uncertainties in predictions.

### Evaluation on the yeast dataset

The cell features were extracted using the MT-DNA-Protein variant of SubCell and SubCell-P. Using the provided training and test sets, we trained each model with 10 different random seeds and averaged the resulting performance metrics and confusion matrices across seeds. We retrained the original CycleNET and DeepLoc [28] models on the cell cycle classification and subcellular localization tasks, respectively. We also trained a multilayer perceptron (MLP) classifier trained directly on the image pixels. We chose to include the additional pixel-based baseline because of the small size of the yeast images (64x64, 4096 pixels) and to show that even though SubCell models, which produce 1536-dimensional embeddings, can capture more than simple pixel intensities for such small images. For SubCell, we first preprocessed the image data by applying global min-max normalization and resizing the image crops to 1×, 2×, 3×, or 4× their original dimensions. We then generated feature embeddings using the two best-performing models. To achieve this, we mapped the imaging channels as follows: the nucleus and bud neck channels were assigned to the SubCell nucleus channel, the cytoplasm channel to the SubCell microtubule channel, and the GFP-tagged protein channel to the SubCell protein channel. We trained both two-layer MLPs and logistic regression classifiers using these SubCell embeddings as input features. For comparison, we also trained classifiers directly on globally min-max normalized pixel values. MLPs were trained for up to 100 epochs, with the best-performing model selected based on test set accuracy. All reported results are computed on the provided test set.

### Analysis of the factors of variation and subsequent integration

As shown in Doron et al. [19], self-supervised features often encode diverse factors of variation that are learned during training. Since the HPA dataset was acquired over more than a decade using different microscopes and experimental setups, we found that SubCell features also encode several technical variations, namely, variations across cell lines, microscopes, and plates (See Supplementary Fig. S14). We utilized Harmony [88], an algorithm designed for gene expression data, to integrate across these factors of variation and remove confounding effects. Specifically, we integrate the features across cell lines, plate IDs, and the microscopes.

We employed a mutual information-based metric [89], following the same procedure as Doron et al. [19], to investigate the hierarchy of features with respect to the factors of variation: protein localization, cell lines, and well IDs. We first calculate a discrete joint distribution of a factor of variation. The number of bins for the distribution was set to 50. Next, the mutual information between the factors of variation and individual features was calculated, estimating how informative the feature is to each factor of variation. Finally, the features were ranked by the factor of variation for which they were most informative, and the percentage of features most informative for each factor of variation was calculated and shown in Supplementary Fig. S14C. SubCell-P features, the significantly larger features were informative for well IDs than SubCell (original and microscopy-harmonized features), highlighting the bias present in the model.

For subsequent biological analysis, we evaluated different harmonized features and presented the best results.

### Analysis of biologically meaningful representations

#### Aggregating profiles over cell populations

We followed the same methodology as Doron et al. [19] to construct similarity matrices for cell lines and protein localizations. We averaged all populations across the categories and retained the top 10 principal components. We removed the cell lines and localization categories that were not present in the bulk RNA-seq data and the protein hierarchy, respectively. We then constructed the hierarchical representation of the populations using cosine similarities between population representations. The ground-truth genetic relationships were established by bulk mRNA sequencing, and the localization hierarchy was defined by expert annotation, following an approach similar to that of Doron et al. [19]. Comparisons of the models’ representations with the ground truth were assessed using the Mantel statistic [90].

#### Evaluating protein relations across known biology

The single-cell features were combined across proteins to create the protein representation. We utilized the Reactome [32], Gene Ontology (GO) [33] cellular compartment (CC), and biological process (BP) databases for our analysis. Protein relationships were used to establish the hierarchy among groups in the database. The top two levels of this hierarchy were used in our evaluation. Protein groups with fewer than ten members were excluded, and proteins present in more than five groups were also removed. For GO:BP level 1, 8208 proteins across 63 groups were analyzed, while at level 2, 5895 proteins across 108 groups were included. For GO:CC level 1, 8784 proteins across 62 groups were used, and at level 2, 8734 proteins across 86 groups were used. For Reactome, level 1 included 6692 proteins in 28 groups, and level 2 included 6382 proteins in 131 groups. The datasets were divided into different training and validation sets for 10-fold cross-validation. We perform a hyperparameter search to find the optimal parameters resulting in the best performance for each configuration and model.

The hyperparameters that were optimized were the number of layers: {1, 2, 3}, hidden MLP dimensions: {128, 256, 512}, and dropout: {0.0, 0.1, 0.2, 0.3, 0.4, 0.5, 0.6, 0.7, 0.8}.

The experiments were repeated three times with different training and validation sets to get N=30 for statistical significance.

#### Evaluating attention maps

We assessed the underlying transformer’s attention pooling module to examine which parts of an image SubCell learned that capture biologically meaningful information. ViTs convert an image into a sequence of patches and leverage an attention mechanism between them to learn complex features. The attention generated for each patch provides valuable insights into the model’s behavior and decision-making process. We quantified the regions our models attend to in a cell by calculating the correlation between the attention maps and the input channels. For the attention analysis, we used the output of the last-layer attention head of the ViT model with respect to the [CLS] token and the pooled attention heads. For correlation evaluation between input channels and attention maps, we normalize each input channel and each attention map to the range [0, 1]. The correlation was calculated for each input channel and the attention map, resulting in a 14 × 4 = 56-dimensional feature vector for each cell. For the hierarchy analysis, only the protein channel was used. To perform the UMAP visualization of the correlation profiles, we aggregated them across FOVs and harmonized them across microscopes to remove that source of variation.

To further quantify whether the models encode cell-line identity and protein localization through shared or independent directions in attention space, we computed the geometric overlap between the embedding directions most informative for distinguishing cell lines and those most informative for distinguishing localization patterns. We identified the ten directions in the embedding space that best separate by cell line using regularised Linear Discriminant Analysis (LDA) with Ledoit-Wolf shrinkage estimation of the within-group covariance matrix [91], which stabilizes the solution when the number of groups is large relative to the number of features (37 cell lines vs 56 feature dimensions). We repeated this procedure independently for each of the 34 localization categories using binary LDA, which identifies the single direction along which the positive and negative for a given localization label are maximally separated relative to their within-group spread using Fisher linear discriminant [92]. The 34 resulting discriminant directions were combined into a single localization-discriminative subspace via singular value decomposition [93], retaining the top ten components. The overlap between the cell-line subspace W_cell_ and the localization subspace W_loc_ was then quantified using the subspace overlap coefficient, defined as the mean squared cosine of the principal angles *θ*_1_ ≤ *θ*_2_ ≤ · · · ≤ *θ_k_* between the two subspaces [94] where overlap = 0 indicates completely independent directions and overlap = 1 indicates identical directions. To assess whether the observed overlap between the cell-line and localization subspaces was greater than expected by chance alone, we generated a null distribution by computing the overlap between W_cell_ and 1,000 randomly oriented subspaces of the same dimensionality, sampled uniformly using the Haar measure on the Grassmannian manifold Gr(*k, D*) [95]; the *p*-value was then defined as the proportion of these random subspaces whose overlap with W_cell_ equalled or exceeded the observed overlap, such that a small *p*-value indicates that the true localization subspace is more aligned with the cell-line subspace than would be expected if the two sets of directions were unrelated. The expected overlap between two such unrelated random subspaces of rank *k* in a *D*-dimensional space is given analytically as [94]:

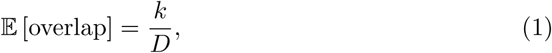

which equals 10*/*56 = 0.179 for SubCell and SubCell-P, and 10*/*48 = 0.208 for DINO. A normalized excess overlap was computed as:

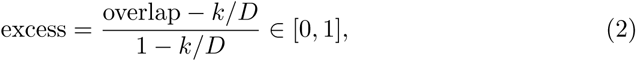

providing an effect size directly comparable across models with different embedding dimensions, where 0 indicates overlap at the level of unrelated random subspaces, and 1 indicates maximum possible entanglement. Uncertainty in all estimates was quantified by bootstrap 95% confidence intervals (500 resamples) [96].

Applying this analysis to 79,402 FOV correlation profiles from 37 cell lines and 34 localization categories, the three models showed markedly different degrees of overlap between their cell-line and localization directions. SubCell showed no detectable entanglement: its overlap coefficient of 0.180 (95% CI: [0.167, 0.205]; *p* = 0.480) was statistically indistinguishable from the random expectation of 0.179, with a normalised excess of 0.001 and a mean angle of 68.5^°^±15.3^°^ between the two sets of directions, with only one of ten direction pairs showing collinearity below 45^°^ (40.3^°^). DINO showed modest but statistically significant entanglement, with an overlap of 0.270 (95% CI: [0.244, 0.297]; *p* = 0.008), 7.8% above the random expectation of 0.208, and a mean angle of 61.9^°^ ± 18.2^°^, with two direction pairs falling below 45^°^ (43.7^°^ and 27.2^°^), indicating that a small subset of directions jointly encodes both cell-line and localization information. SubCell-P was the most severely entangled, with an overlap of 0.538 (95% CI: [0.512, 0.561]; *p* = 0.001), three times the random expectation of 0.179 and a normalised excess of 0.438, and a mean angle of only 42.8^°^ ± 28.1^°^, with five of ten direction pairs below 45^°^ including four below 20^°^, demonstrating that the majority of the most discriminative directions in this embedding are shared between cell-line and localization. Together, these results establish a clear ranking of representational independence (SubCell *>* DINO *>* SubCell-P), with SubCell being the only model whose cell-line and localization information occupies statistically independent directions in feature space.

We performed a similar analysis for SubCell-P (Supplementary Fig. S15). We found that SubCell-P lags behind SubCell in capturing population-level phenotypic information for both protein localization hierarchy and cell identity (Supplementary Fig. S15C). Conversely, SubCell-P significantly outperforms SubCell in predicting known biological relations within the GO:CC and Reactome databases (Supplementary Fig. S15D). Furthermore, unlike SubCell, the attention maps of SubCell-P fail to localize to specific subcellular compartments, with several attention heads yielding entirely empty representations (e.g., AH-4, AH-11; Supplementary Fig. S15E). However, as previously noted, the enhanced protein-relation information in SubCell-P remains entangled with confounding factors (similarly noted in Supplementary Fig. S14D). Consequently, it is difficult to use SubCell-P as a generalizable model across diverse downstream tasks or to resolve multi-scale subcellular organization in an unbiased manner.

### Drug Prediction on the CM4AI dataset

The dataset was divided into three treatment categories: Paclitaxel, untreated, and Vorinostat. We perform a ten-fold cross-validation for the drug prediction task on the CM4AI dataset by dividing it into training and validation sets, stratified by protein, with an 80/20 split. The classifiers were trained on cell features extracted by the models. For each fold, ten classifiers were trained using the same hyperparameter search as the previous section. Finally, results from all folds were aggregated, and final classification metrics were computed.

#### Drug-Induced Perturbation Signatures

To assess whether SubCell embeddings capture drug-induced phenotypic changes beyond what is detectable by image intensity alone, we compared perturbation signatures derived from two complementary readouts: single-cell image intensity and SubCell embedding distributions for Paclitaxel- and Vorinostat-treated cells relative to untreated controls. Proteins with known imaging artifacts were excluded, and only proteins with at least 20 cells in both conditions were retained.

##### Intensity analysis

For each protein, the log_2_-transformed 90th-percentile single-cell image intensity (log_2_(cell q90+1)) was compared between treated and untreated cells using a two-sided Mann–Whitney U test as implemented in SciPy [97] (v1.14.0). *P* -values were corrected across proteins using the Benjamini–Hochberg (BH) procedure [98] as implemented in statsmodels [99] (v0.14.2). Intensity significance was defined as | log_2_ FC| *>* 2 and BH-adjusted *p <* 0.01.

##### Embedding analysis

Distributional shifts in SubCell embedding space were quantified using the Sliced Wasserstein Distance (SWD) [100], estimated from 1,000 random one-dimensional projections (random seed 42) using SciPy [97]. For each projection direction *θ*, sampled from a standard normal distribution and *ℓ*_2_-normalized, the one-dimensional Wasserstein distance between projected treated and untreated distributions was computed; SWD was taken as the mean across projections. Statistical significance was assessed by applying a two-sided Mann-Whitney U test independently to each embedding dimension; per-dimension *p*-values were combined using Simes’ method [101], which controls the familywise error rate under positive regression dependence, and the resulting protein-level *p*-values were corrected across proteins using the Bonferroni procedure (both corrections via statsmodels [99] v0.14.2). Embedding significance was defined as SWD *> d̄*+ 1.5*σ*, where *d̄* and *σ* are the mean and standard deviation of SWD across all proteins, and Bonferroni-adjusted *p <* 0.001.

##### Joint classification

Proteins were classified by joint significance across both analyses: significant by intensity only, embedding only, both, or neither. Results are visualized as a 2D scatter plot (log_2_ FC vs. SWD; Fig. 4C) and as marginal volcano plots (Supplementary Fig. S6).

### JUMP Evaluation

For DINO4Cells and SubCell, JUMP images were illumination-corrected, per-channel min-max normalized, and cropped (https://github.com/zwefers/subcell_jump_analysis). JUMP crops for DeepProfiler were created via the DeepProfiler package “prepare” and “export-sc” commands. DeepProfiler embeddings were extracted from the penultimate feature layer of the model, as recommended in the Deep-Profiler documentation (https://cytomining.github.io/DeepProfiler-handbook/docs/00-welcome.html). For performing inference with SubCell, single-cell crops were upscaled to match the physical pixel size of HPA images, and cell segmentation was used to mask out peripheral cells. We used the ER-DNA-Protein version of SubCell because it includes two common Cell Painting channels (ER, DNA). We embedded single-cell crops by processing three-channel subsets through SubCell’s 3-channel model and concatenating the resulting embeddings: (ER, DNA, Mito), (ER, DNA, AGP), and (ER, DNA, RNA) (as shown in Supplementary Fig. 2B).

Raw single-cell profiles underwent several standard post-processing steps before being used in downstream analysis. First, single-cell profiles are combined with FOV profiles by either mean or median aggregation. Second, FOVs are mean-aggregated to well-level profiles. Feature selection is optionally applied to well-level profiles using variance and feature-correlation thresholds. Next, profiles underwent two rounds of normalization. 1) Z-score or MAD-robustize standardization was used to center the well-level profiles of each plate. 2) Sphering with respect to negative control wells across all plates was used to account for plate-level batch effects. Both ZCA and PCA sphering transforms were tested. All models were tested with all combinations of post-processing steps. This included switching the order of normalization steps and omitting sphering altogether. Feature selection, standardization, and sphering were performed with the pycytominer [102] package. The optimal post-processing pipeline was selected independently for each model by computing a Pareto frontier over replicate mAP and MoA mAP; no set of configurations can improve one retrieval metric without making the other worse (Supplementary Fig. S16). We chose a configuration that lay on the frontier for both cell types. For all methods, except Cell Profiler, there was only a configuration on the frontier for both cell types. For CellProfiler, we choose the configuration that achieves significantly better replicate mAP but slightly worse MoA mAP. The configurations chosen for each model are reported in Supplementary Table S11.

All evaluations were conducted separately for A549 and U2OS cells, as cell-type-specific morphology confounds perturbation-specific morphology, making cross-cell-type comparisons uninformative.

Well-level profiles were evaluated using mean average precision (mAP) computed with the copairs package (https://github.com/cytomining/copairs) using cosine similarity as a distance metric [103]. We report the mAP “non-rep” variant for both replicate retrieval and MoA identification tasks. In this variant, for a given query well, positive pairs are wells on different plates that share the same label, while negative pairs are wells on the same plate with a different label. For replicate retrieval, the label is the compound identity; for MoA retrieval, the label is the mechanism of action. This construction tests whether a biologically meaningful signal is retrieved more strongly than plate-level batch effects. Copairs supports multilabel matching for MoA labels, so we consider two wells a match if they share at least one MoA in common. Because the mAP value is heavily affected by the number of positive and negative pairs, the copairs package computes corrected p-values, which we use to determine which MoAs the models retrieve at a significant rate.

UMAP embeddings were computed on the post-processed well-level profiles from each model’s selected configuration using cosine distance, with *n neighbors* = 100 and *min dist* = 0.25. Qualitatively, compared to the other methods, SubCell shows better separation of treatment and control profiles and a less pronounced well effect, but a more pronounced batch effect (Supplementary Fig. S17-20). This confirms that SubCell captures biological signal over technical well-to-well variation, but that sphering may be an insufficient batch correction method for SubCell embeddings.

We extracted self-attention maps from ViT attention heads and the pooled attention heads of both SubCell and DINO4Cells-CP. JUMP crops were upscaled 3.74 to match the HPA physical pixel size for SubCell inference. Although DINO4Cells was trained on standard JUMP crops, at that resolution, its 16×16 patch size yields only an 8×8 attention grid, which was too coarse to resolve subcellular patterns. We therefore up-scaled images for attention visualization to enable direct visual comparison with SubCell (Supplementary Fig. S9).

### Vision-based multiscale cell map construction

The HPA dataset was embedded using the SubCell model. Further processing and analysis of the embeddings was conducted in Python 3.10.14 using ScanPy 1.10.2. To remove unwanted sources of variance from the HPA protein single-cell embeddings, the embeddings were normalized for the covariates plate ID and microscope using the PyTorch implementation of the Harmony algorithm [88].

For this analysis, we focused on the cell line with the highest number of stained proteins, U2OS. The single-cell image embeddings were aggregated into one average embedding vector for each protein. For this, all antibodies with a reliability rating of “uncertain” were removed, as well as those that stained more than one protein. Additionally, if there were stains from multiple antibodies for a protein, the most reliable antibody was selected based on the HPA antibody reliability score when the antibodies had different ratings; otherwise, one antibody was chosen at random.

The resulting protein embedding, consisting of 9386 proteins, was (sub)clustered at multiple resolutions (0, 0.15, 0.2, 0.25, 0.3, 0.33, and 0.36) using the Leiden algorithm [39] with decreasing numbers of neighbors in the corresponding neighborhood graph (125, 100, 90, 55, 40, 25, 10) to focus on the global structure at lower resolutions and local structure at higher resolutions. For this, all proteins were clustered at a resolution of 0; the resulting clusters were then subclustered at 0.15, and so on. From this, a hierarchical graph was constructed, with a node for each cluster at each hierarchy level, and edges were drawn to indicate which lower-resolution clusters yielded which higher-resolution clusters.

Additionally, a functional enrichment analysis was performed for each cluster using the G:Profiler Python package v1.0.0 [104]. The Gene Ontology (GO) Cellular Component ontology was used for resolutions 0, 0.15, 0.2, 0.25, 0.3, and 0.33, and GO Biological Processes and GO Molecular Functions ontologies were used for the highest resolution. The human genome was set as background, the g:SCS algorithm was used to compute multiple testing corrections for p-values, and a significance threshold of *p <* 0.05 was selected. Each node in the hierarchy graph was labeled with the most significantly enriched term.

### Multimodal Integration with sequence data

We used the unsupervised MuSIC [37] co-embedding scheme to integrate protein image and protein sequence embeddings, yielding a final 128-dimensional embedding. Another unsupervised integration method, MIRAGE [62], was evaluated; however, the enrichment results in the databases did not surpass those of MuSIC (Supplementary Fig. S10B). We maintained the same hyperparameters and training settings as those used in the original work. We applied the same cluster-transfer method from the SubCell to the multimodal space and from ESM-2 to the multimodal space to assess whether the multimodal space maintained clustering with respect to both.

All performances are evaluated using an enrichment metric rather than the learned classifiers’ metrics because many of the biological labels evaluated in this section are sparse. The problem of sparse function labels extends beyond those evaluated here, underscoring the general importance of a functionally enriched embedding space and, consequently, of using enrichment as the evaluation approach. We measure enrichment using the approximated Cliff’s Delta [105].

#### Establishing differences between SubCell and ESM-2 embedding spaces

To investigate the orthogonality between image (SubCell) and sequence (ESM-2) embeddings, we first created a set of image embeddings using the SubCell with microscope and plate ID harmonization and performed Leiden clustering in the same way as described in the previous section, using the same gene set (those from HPA U2OS data). Then, we used the ESM-2-650M model [58] to embed proteins by sequence and clustered them in the same way. We used the canonical UniProt sequence for each. We then assigned the cluster labels from the SubCell embeddings to the proteins’ ESM-2 embeddings. We created 2D UMAPs as a rough visualization of how well the clusters transferred. No apparent clusters were observed in the ESM-2 embeddings, especially at higher resolutions. The same process was repeated, this time clustering the ESM-2 embeddings of the proteins and then transferring the cluster labels to the SubCell embeddings. This yielded the same outcome. To quantify these differences, we calculated the average percentile-rank similarities for sequence cluster labels in each of the three embedding spaces (image, sequence, and multimodal), as well as for image cluster labels, across clustering resolutions. We also included random baselines for both.

#### Assessing multimodal enrichment of coarse and sub-organelle localization

We used the HPA annotations of the images used for the SubCell embeddings as coarse localization labels, mapped to the low-level annotations in Supplementary Table S2. We performed Cliff’s Delta analysis on embeddings that are mean-centered within each individual space, sampling 500 comparisons 500 times.

To get sub-organellar localization, we used GO-CC to find proteins associated with each manually selected category. Because GO is an aggregate annotation source, there are proteins in each GO sub-organellar category that did not overlap with the HPA-derived parent organelle labels that are specific to the images we embedded for this experiment. For example, some proteins in the GO-CC category for the mitochondrial matrix were not labeled as mitochondrial proteins in the HPA images used here. To reduce variance, we subsetted the GO-CC proteins for each suborganelle to those that also showed up in the HPA-derived labels for the associated parent organelle. Then, we performed Cliff’s Delta analysis, sampling 500 comparisons 500 times. ESM-2’s worse performance in coarse annotations and better performance in sub-compartment annotations can be explained by the fact that sub-compartments may be enriched for specialized proteins with tuned characteristics (e.g., hydrophobic residues on membrane proteins), thereby providing a more productive signal at that level.

#### Assessing enrichment of interacting pairs in the multimodal space

To assess whether the multimodal embedding encodes relevant biological information, we constructed a similarity enrichment evaluation task, employing a strategy similar to that of Schaffer et al. We used the CORUM [64], hu.MAP [65], BioPlex [63], and STRING [40] databases for the analysis. For the CORUM dataset, we used 2,708 proteins with 88,288 interactions, for the hu.MAP dataset, we used 6,334 proteins with 33,245 interactions. For the BioPlex dataset, we used 8,593 proteins with 54,748 interactions, and for the STRING database, we used 8,956 proteins with 107,597 interactions. We first compute similarities between protein embeddings and partition the databases based on interactions. We then calculate the Cliff’s Delta for the similarity between the interacting and non-interacting sets of proteins. We repeatedly sample 10,000 interacting and non-interacting protein pairs, each set sampled 1,000 times, to establish statistical significance.

We performed this analysis comparatively across different image models (DINO4Cells-HPA, bestfitting, and SubCell) in Supplementary Fig. S10B. In that analysis, we included an additional multimodal integration method (MIRAGE [106]) to compare different co-embedding methods.

#### Detecting functionally divergent paralogs

To compare the organization of the sequence vs. multimodal embedding spaces, we computed all pairwise cosine similarities between proteins in each space, then ranked them by percentile. This creates a normalized metric with which we can compare the different embedding methods. For each pair, we plotted the percentile rank in sequence space vs. multimodal space. We used the HPA-derived annotations from the images to label protein pairs in which each member has the exact same set of locations, and those with no overlap in annotated locations. All other proteins in the plot have partial overlap of their localization annotations.

We searched Uniprot for examples of paralogs with both conserved and divergent locations (as differences in localization are a signal of functional divergence) and compared their ranked percentile similarity in both sequence and image space. The same analysis was done to compare image vs. multimodal space as well(Supplementary Fig. S10C). Sample images are from the HPA U2OS collection.

#### Multimodal enrichment for biological processes

To select biological processes for evaluation, we used the entire gene set for which we had embeddings (i.e., the gene set for which we had images to create a SubCell embedding) and performed Gene Ontology Biological Process enrichment using g-Profiler. The results included Biological Processes (BP) that were statistically enriched in our gene set (*p <* 0.001 in Profiler enrichment analysis). Since Gene Ontology is a tree-based annotation system, we removed any BPs that were parents of other BPs from our results. We also removed any BPs that were clearly cell-type specific (e.g., Myoblast differentiation). Finally, we removed BPs that were dependent on unique cell states, like those associated with viral infection or cell differentiation. This resulted in a final set of 158 BPs. We then computed Cliff’s Delta for these using 5,000 pairs over 500 iterations and ranked them by multimodal embedding enrichment. The top 25 best-performing BPs in the multimodal space are shown in Fig. 6E and the rest in Supplementary Fig. S11.

## 2 Data availability

The model and data are available at: https://virtualcellmodels.cziscience.com/. The HPA dataset is available in the Human Protein Atlas database (www.proteinatlas.org). The cropped image dataset used for training the models is available in the S3 bucket: s3://czi-subcell-public/

## 3 Code availability

The code necessary for training all the models is available in the following repositories: https://github.com/CellProfiling/subcell-embed and https://github.com/czi-aisub-cell-embed.

The code necessary for inference is available in this repository: https://github.com/czi-ai/SubCellPortable.

The code necessary for reproducing the results in the analysis and generating the figures is available in this repository: https://github.com/CellProfiling/subcell-analysis.

The proteome-wide map of cellular substructures is available as an interactive version in the Human Protein Atlas (to be released upon full publication).

## 4 Acknowledgements

Training and evaluation of the models was enabled by computing resources provided by the Chan Zuckerberg Initiative and the National Supercomputer Centre at Linköping University. EL was supported by the Chan Zuckerberg Initiative, the Wallenberg Foundation (2021.0346), Schmidt Futures, Göran Gustafsson Foundation, the Bridge2AI Program (NIH Common Fund; OT2 OD032742), the Cancer Cell Map Initiative (NCI Center for Cancer Systems Biology; U54 CA274502), Stanford Institute for Human Centered AI, and Param Hansa Philanthropies. JNH was supported by a Postdoctoral Fellowship from the Wenner-Gren Foundations and by an EMBO Postdoctoral Fellowship (ALTF 556-2022). We thank Anna Martinez Casals, Stanford University, for assistance with the tissue work and Cecilia Lindskog, HPA tissue section, for providing the tissue material.

## 5 Author contributions

AG and EL conceptualized the project and designed the study. AG designed and trained the models. AG, ZW, KK, MS, and JNH evaluated model performance across tasks. UA, JNH, and WL generated and curated data for evaluations. FB prepared data for model training. AG, ZW, KK, JNH, MKM, AC, WL, and EL analyzed and interpreted the results. FB, DL, and TK contributed to building model-sharing frameworks. AG, ZW, KK, MKM, and JNH prepared figures. AG, ZW, KK, MKM, JNH, WL, and EL wrote the paper. TK contributed to methodology discussions. EL supervised the project. All authors edited and commented on the manuscript.

