## Supplemental information for "SubCell: Proteome-aware vision foundation models for microscopy capture single-cell biology"

#### 1 Supplementary Tables

| Base Model | Prediction Head | Resize (pixels) | Cell-cycle | | | Localization | | |
|---|---|---|---|---|---|---|---|---|
| | | | Acc | Macro F1 | Macro AP | Acc | Macro F1 | Macro AP |
| SubCell | LogReg | - | 0.840 | 0.669 | 0.691 | 0.599 | 0.564 | 0.581 |
| | | 128 | 0.889 | 0.734 | 0.786 | 0.659 | 0.627 | 0.665 |
| | | 256 | 0.931 | 0.835 | 0.890 | 0.730 | 0.698 | 0.768 |
| | | 512 | 0.946 | **0.865** | 0.903 | 0.747 | 0.721 | 0.789 |
| | MLP | - | 0.833 | 0.584 | 0.616 | 0.547 | 0.510 | 0.532 |
| | | 128 | 0.887 | 0.719 | 0.759 | 0.677 | 0.645 | 0.697 |
| | | 256 | 0.927 | 0.809 | 0.850 | 0.725 | 0.698 | 0.750 |
| | | 512 | 0.929 | 0.825 | 0.860 | 0.729 | 0.700 | 0.760 |
| SubCell-P | LogReg | - | 0.840 | 0.596 | 0.700 | 0.588 | 0.548 | 0.578 |
| | | 128 | 0.853 | 0.626 | 0.730 | 0.678 | 0.639 | 0.693 |
| | | 256 | 0.852 | 0.595 | 0.691 | 0.685 | 0.649 | 0.698 |
| | | 512 | 0.808 | 0.488 | 0.620 | 0.647 | 0.610 | 0.637 |
| | MLP | - | 0.844 | 0.667 | 0.703 | 0.571 | 0.527 | 0.562 |
| | | 128 | 0.858 | 0.656 | 0.699 | 0.669 | 0.631 | 0.698 |
| | | 256 | 0.872 | 0.662 | 0.709 | 0.689 | 0.652 | 0.703 |
| | | 512 | 0.855 | 0.602 | 0.666 | 0.649 | 0.608 | 0.653 |
| Pixel | LogReg | - | 0.829 | 0.628 | 0.660 | 0.271 | 0.228 | 0.225 |
| | MLP | - | 0.874 | 0.746 | 0.784 | 0.466 | 0.414 | 0.432 |
| CycleNet | - | - | **0.950** | **0.865** | **0.918** | - | - | - |
| DeepLoc | - | - | - | - | - | **0.788** | **0.757** | **0.825** |

**Table S1**: Results for all models trained on the yeast datasets for cell-cycle stage prediction and protein localization predictions. Metrics shown are averaged over ten random seeds.

| Original Annotation | Low-level Annotations | High-level Annotations | OpenCell Annotations |
|---|---|---|---|
| Actin filaments | Actin filaments | Cytoskeleton | cytoskeleton |
| Aggresome | | | |
| Cell Junctions | Plasma membrane | Plasma membrane | cell_contact |
| Centriolar satellite | Centrosome | Cytosol | centrosome |
| Centrosome | Centrosome | Cytoskeleton | centrosome |
| Cleavage furrow | | | |
| Cytokinetic bridge | | | |
| Cytoplasmic bodies | Cytosol | Cytosol | big_aggregates |
| Cytosol | Cytosol | Cytosol | cytoplasmic |
| Endoplasmic reticulum | Endoplasmic reticulum | Endomembrane system | er |
| Endosomes | Vesicles | Endomembrane system | vesicles |
| Focal adhesion sites | Actin filaments | Cytoskeleton | focal_adhesions |
| Golgi apparatus | Golgi apparatus | Endomembrane system | golgi |
| Intermediate filaments | Intermediate filaments | Cytoskeleton | cytoskeleton |
| Lipid droplets | Vesicles | Endomembrane system | vesicles |
| Lysosomes | Vesicles | Endomembrane system | vesicles |
| Microtubule ends | Microtubules | Cytoskeleton | cytoskeleton |
| Microtubules | Microtubules | Cytoskeleton | cytoskeleton |
| Midbody | | | |
| Midbody ring | | | |
| Mitochondria | Mitochondria | Mitochondria | mitochondria |
| Mitotic chromosome | | | |
| Mitotic spindle | | | |
| Nuclear bodies | Nucleus | Nucleus | nuclear_punctae |
| Nuclear membrane | Nuclear membrane | Nucleus | nuclear_membrane |
| Nuclear speckles | Nucleus | Nucleus | chromatin |
| Nucleoli | Nucleoli | Nucleoli | nucleolus_gc |
| Nucleoli fibrillar center | Nucleoli | Nucleoli | nucleolus_fc_dfc |
| Nucleoli rim | Nucleoli | Nucleoli | |
| Nucleoplasm | Nucleus | Nucleus | nucleoplasm |
| Peroxisomes | Vesicles | Endomembrane system | vesicles |
| Plasma membrane | Plasma membrane | Plasma membrane | membrane |
| Rods & Rings | | | |
| Vesicles | Vesicles | Endomembrane system | vesicles |

**Table S2**: Original localization categories in the HPA dataset with their low-level, high-level, and OpenCell grouping annotated by experts.

| |
|---|
| **Note:** The data is provided as a separate Excel (.xlsx) file in the supplementary materials due to its scale. |

**Table S3**: Excel file containing proteins contained in each cluster and the functional enrichment results of the vision-based multiscale cell map in Figure 5B.

| Dataset | Subset | Proteins | FoVs | Cells | Pixel Size ($\mu$m) |
|---|---|---|---|---|---|
| HPAv23 | Train | 10,433 | 58,480 | 812,688 | 0.0801 |
| | Validation | 1,487 | 8,428 | 103,980 | 0.0801 |
| | Test | 3,460 | 18,041 | 221,358 | 0.0801 |
| Kaggle | Test | 851 | 1,112 | 21,880 | 0.0801 |
| OpenCell | Test | 1,310 | 6,301 | 94,426 | 0.2063 |
| AllenCell | Train | 25 | 18,079 | 136,983 | 0.1083 |
| | Validation | 25 | 15,260 | 34,246 | 0.1083 |
| | Test | 25 | 16,217 | 42,808 | 0.1083 |
| Yeast(Cell Cycle) | Train | NA | NA | 6,633 | 0.1077 |
| | Test | | | 764 | 0.1077 |
| Yeast(Protein Localization) | Train | NA | NA | 16,305 | 0.1077 |
| | Test | | | 1,824 | 0.1077 |
| U2OS FUCCI | Test | 1,166 | 2,798 | 357,083 | 0.1625 |
| Bridge2AI | Test | 474 | 1,404 | 131,206 | 0.0700 |
| JUMP | Test | NA | 3,456 | 1,734,123 | 0.3000 |

**Table S4**: Distribution of datasets into training, validation, and test sets used in this work.

# A.

| Dataset | Experiment | Macro AP | Micro AP | Label Rank. AP | Coverage Error |
|---|---|---|---|---|---|
| HPAv23 | MAE-MR0.25-CMR-0.0 | 0.518 ± 0.0039 | 0.775 ± 0.0014 | 0.827 ± 0.001 | 2.57 ± 0.0084 |
| | MAE-MR0.25-CMR-0.25 | 0.514 ± 0.0037 | 0.773 ± 0.0013 | 0.825 ± 0.0009 | 2.58 ± 0.0091 |
| | MAE-MR0.25-CMR-0.5 | *0.523 ± 0.0035* | *0.778 ± 0.0013* | *0.829 ± 0.0013* | *2.551 ± 0.0124* |
| | MAE-MR0.33-CMR-0.0 | 0.515 ± 0.0043 | 0.772 ± 0.0018 | 0.824 ± 0.0013 | 2.591 ± 0.0121 |
| | MAE-MR0.33-CMR-0.25 | 0.519 ± 0.0027 | 0.775 ± 0.001 | 0.827 ± 0.001 | 2.568 ± 0.0096 |
| | MAE-MR0.33-CMR-0.5 | **0.528 ± 0.0045** | **0.78 ± 0.0017** | **0.831 ± 0.0013** | **2.535 ± 0.0114** |
| | MAE-MR0.5-CMR-0.0 | 0.493 ± 0.0039 | 0.76 ± 0.0017 | 0.816 ± 0.0015 | 2.663 ± 0.011 |
| | MAE-MR0.5-CMR-0.25 | 0.493 ± 0.0036 | 0.765 ± 0.0011 | 0.818 ± 0.0011 | 2.635 ± 0.0082 |
| | MAE-MR0.5-CMR-0.5 | 0.512 ± 0.0036 | 0.773 ± 0.0013 | 0.826 ± 0.0009 | 2.573 ± 0.0092 |
| | DINO4Cells-HPA | **0.705 ± 0.0022** | **0.863 ± 0.001** | **0.895 ± 0.0007** | **2.013 ± 0.0059** |
| Kaggle | MAE-MR0.25-CMR-0.0 | 0.355 ± 0.0036 | 0.545 ± 0.0013 | 0.652 ± 0.0011 | 4.895 ± 0.0161 |
| | MAE-MR0.25-CMR-0.25 | 0.348 ± 0.0031 | 0.544 ± 0.0017 | 0.65 ± 0.0016 | 4.947 ± 0.0227 |
| | MAE-MR0.25-CMR-0.5 | *0.361 ± 0.0019* | **0.552 ± 0.0011** | **0.657 ± 0.0016** | **4.838 ± 0.0237** |
| | MAE-MR0.33-CMR-0.0 | 0.354 ± 0.0033 | 0.544 ± 0.0019 | 0.651 ± 0.0016 | 4.919 ± 0.024 |
| | MAE-MR0.33-CMR-0.25 | 0.355 ± 0.0032 | 0.547 ± 0.0014 | 0.652 ± 0.0017 | *4.872 ± 0.0263* |
| | MAE-MR0.33-CMR-0.5 | **0.363 ± 0.0035** | *0.548 ± 0.0018* | *0.654 ± 0.0015* | 4.873 ± 0.0244 |
| | MAE-MR0.5-CMR-0.0 | 0.329 ± 0.0035 | 0.53 ± 0.0023 | 0.639 ± 0.002 | 5.012 ± 0.0263 |
| | MAE-MR0.5-CMR-0.25 | 0.335 ± 0.0027 | 0.529 ± 0.0018 | 0.638 ± 0.0017 | 5.03 ± 0.0284 |
| | MAE-MR0.5-CMR-0.5 | 0.349 ± 0.003 | 0.54 ± 0.0022 | 0.645 ± 0.0022 | 4.941 ± 0.0276 |
| | DINO4Cells-HPA | **0.519 ± 0.004** | **0.631 ± 0.0033** | **0.718 ± 0.0026** | **4.047 ± 0.0381** |

# B.

| Dataset | Experiment | Macro AP | Micro AP |
|---|---|---|---|
| HPAv23 | MAE-MR0.25-CMR-0.0 | 0.923 ± 0.0073 | 0.977 ± 0.0014 |
| | MAE-MR0.25-CMR-0.25 | 0.93 ± 0.0031 | 0.978 ± 0.0007 |
| | MAE-MR0.25-CMR-0.5 | 0.935 ± 0.0042 | 0.981 ± 0.0008 |
| | MAE-MR0.33-CMR-0.0 | 0.924 ± 0.003 | 0.979 ± 0.0006 |
| | MAE-MR0.33-CMR-0.25 | 0.928 ± 0.0042 | 0.979 ± 0.0009 |
| | MAE-MR0.33-CMR-0.5 | *0.937 ± 0.0047* | *0.981 ± 0.0007* |
| | MAE-MR0.5-CMR-0.0 | **0.94 ± 0.0041** | **0.982 ± 0.0008** |
| | MAE-MR0.5-CMR-0.25 | 0.894 ± 0.0068 | 0.967 ± 0.0015 |
| | MAE-MR0.5-CMR-0.5 | 0.926 ± 0.0034 | 0.978 ± 0.0006 |
| | DINO4Cells-HPA | **0.957 ± 0.0014** | **0.99 ± 0.0002** |
| Kaggle | MAE-MR0.25-CMR-0.0 | 0.888 ± 0.0086 | 0.928 ± 0.0038 |
| | MAE-MR0.25-CMR-0.25 | 0.905 ± 0.0037 | 0.937 ± 0.002 |
| | MAE-MR0.25-CMR-0.5 | 0.907 ± 0.0045 | 0.941 ± 0.003 |
| | MAE-MR0.33-CMR-0.0 | 0.895 ± 0.0035 | 0.932 ± 0.002 |
| | MAE-MR0.33-CMR-0.25 | 0.898 ± 0.0055 | 0.934 ± 0.0035 |
| | MAE-MR0.33-CMR-0.5 | *0.908 ± 0.0041* | *0.941 ± 0.003* |
| | MAE-MR0.5-CMR-0.0 | **0.912 ± 0.0043** | **0.943 ± 0.0026** |
| | MAE-MR0.5-CMR-0.25 | 0.842 ± 0.0083 | 0.896 ± 0.0051 |
| | MAE-MR0.5-CMR-0.5 | 0.892 ± 0.0027 | 0.932 ± 0.0022 |
| | DINO4Cells-HPA | **0.95 ± 0.0014** | **0.964 ± 0.0014** |

**Table S5**: Results for the MAE experiments for A.) localization prediction and B.) cell-line classification tasks. (MR refers to the overall masking ratio for the input image, and CMR refers to the masking ratio of the cells.)

## A.

| Dataset | Experiment | Macro AP | Micro AP | Label Rank. AP | Coverage Error |
|---|---|---|---|---|---|
| HPAv23 | MAE-MR0.25-CMR-0.0-CellS | *0.656 ± 0.0021* | *0.845 ± 0.0006* | *0.881 ± 0.0004* | *2.112 ± 0.0032* |
| | MAE-MR0.25-CMR-0.5-CellS | 0.648 ± 0.0025 | 0.842 ± 0.0004 | 0.879 ± 0.0005 | 2.133 ± 0.0027 |
| | MAE-MR0.33-CMR-0.0-CellS | 0.648 ± 0.0036 | 0.842 ± 0.0014 | 0.879 ± 0.0008 | 2.129 ± 0.0058 |
| | MAE-MR0.33-CMR-0.5-CellS | 0.641 ± 0.0012 | 0.84 ± 0.0004 | 0.878 ± 0.0006 | 2.136 ± 0.0041 |
| | CellS | **0.666 ± 0.003** | **0.848 ± 0.0006** | **0.884 ± 0.0003** | **2.1 ± 0.0036** |
| | DINO4Cells-HPA | **0.705 ± 0.0022** | **0.863 ± 0.001** | **0.895 ± 0.0007** | **2.013 ± 0.0059** |
| Kaggle | MAE-MR0.25-CMR-0.0-CellS | *0.502 ± 0.0032* | *0.623 ± 0.0014* | *0.715 ± 0.0012* | *4.023 ± 0.0171* |
| | MAE-MR0.25-CMR-0.5-CellS | 0.498 ± 0.0015 | 0.621 ± 0.0012 | *0.715 ± 0.0014* | 4.016 ± 0.0163 |
| | MAE-MR0.33-CMR-0.0-CellS | 0.498 ± 0.0029 | 0.621 ± 0.0013 | *0.715 ± 0.0009* | 4.031 ± 0.0214 |
| | MAE-MR0.33-CMR-0.5-CellS | 0.488 ± 0.004 | 0.616 ± 0.0015 | 0.713 ± 0.0013 | 4.06 ± 0.0113 |
| | CellS | **0.509 ± 0.0032** | **0.625 ± 0.0012** | **0.717 ± 0.0013** | **3.975 ± 0.0166** |
| | DINO4Cells-HPA | **0.519 ± 0.004** | **0.631 ± 0.0033** | **0.718 ± 0.0026** | **4.047 ± 0.0381** |

## B.

| Dataset | Experiment | Macro AP | Micro AP |
|---|---|---|---|
| HPAv23 | MAE-MR0.25-CMR-0.0-CellS | 0.939 ± 0.0014 | 0.985 ± 0.0002 |
| | MAE-MR0.25-CMR-0.5-CellS | **0.941 ± 0.0017** | **0.987 ± 0.0003** |
| | MAE-MR0.33-CMR-0.0-CellS | 0.94 ± 0.0022 | *0.986 ± 0.0003* |
| | MAE-MR0.33-CMR-0.5-CellS | **0.941 ± 0.0014** | *0.986 ± 0.0003* |
| | CellS | 0.916 ± 0.0036 | 0.98 ± 0.0005 |
| | DINO4Cells-HPA | **0.957 ± 0.0014** | **0.99 ± 0.0002** |
| Kaggle | MAE-MR0.25-CMR-0.0-CellS | 0.919 ± 0.0016 | 0.944 ± 0.0007 |
| | MAE-MR0.25-CMR-0.5-CellS | **0.929 ± 0.0015** | **0.948 ± 0.0005** |
| | MAE-MR0.33-CMR-0.0-CellS | 0.923 ± 0.0017 | *0.946 ± 0.0008* |
| | MAE-MR0.33-CMR-0.5-CellS | *0.924 ± 0.0014* | *0.946 ± 0.0009* |
| | CellS | 0.894 ± 0.005 | 0.923 ± 0.0022 |
| | DINO4Cells-HPA | **0.95 ± 0.0014** | **0.964 ± 0.0014** |

**Table S6**: Results for the cell-specific (CellS) experiments for (A) localization prediction and (B) cell-line classification tasks. MAE- refers to the ViT model trained in masked autoencoder setting along with the cell-specific loss.

## A.

| Dataset | Experiment | Macro AP | Micro AP | Label Rank. AP | Coverage Error |
|---|---|---|---|---|---|
| HPAv23 | ProtS | **0.742 ± 0.0047** | **0.878 ± 0.0013** | **0.903 ± 0.0012** | **1.934 ± 0.008** |
| | CellS-ProtS | *0.728 ± 0.0031* | *0.872 ± 0.0004* | 0.899 ± 0.0004 | 1.969 ± 0.0027 |
| | MAE-MR0.25-CMR-0.0-ProtS | 0.709 ± 0.0046 | 0.858 ± 0.0012 | 0.893 ± 0.0004 | 2.03 ± 0.0049 |
| | MAE-MR0.25-CMR-0.5-ProtS | 0.707 ± 0.002 | 0.856 ± 0.0008 | 0.891 ± 0.0006 | 2.034 ± 0.0044 |
| | MAE-MR0.25-CMR-0.0-CellS-ProtS | *0.729 ± 0.0037* | *0.872 ± 0.0011* | *0.901 ± 0.0012* | *1.959 ± 0.0055* |
| | MAE-MR0.25-CMR-0.5-CellS-ProtS | 0.719 ± 0.0036 | 0.869 ± 0.0006 | 0.899 ± 0.0007 | 1.974 ± 0.0049 |
| | DINO4Cells-HPA | 0.705 ± 0.0022 | 0.863 ± 0.001 | 0.895 ± 0.0007 | 2.013 ± 0.0059 |
| | ViT-Weak-Supervised | 0.734 ± 0.0008 | 0.869 ± 0.0004 | 0.898 ± 0.0004 | 1.985 ± 0.0029 |
| | bestfitting | **0.77** | **0.83** | **0.877** | **2.029** |
| Kaggle | ProtS | **0.559 ± 0.0058** | **0.653 ± 0.0017** | *0.733 ± 0.0026* | *3.823 ± 0.0464* |
| | CellS-ProtS | *0.552 ± 0.0035* | *0.652 ± 0.0015* | **0.734 ± 0.0012** | **3.813 ± 0.0186** |
| | MAE-MR0.25-CMR-0.0-ProtS | 0.506 ± 0.0054 | 0.614 ± 0.0067 | 0.708 ± 0.0044 | 4.031 ± 0.0478 |
| | MAE-MR0.25-CMR-0.5-ProtS | 0.505 ± 0.0043 | 0.61 ± 0.0034 | 0.705 ± 0.0028 | 4.035 ± 0.0404 |
| | MAE-MR0.25-CMR-0.0-CellS-ProtS | 0.542 ± 0.0036 | 0.64 ± 0.0032 | 0.725 ± 0.0024 | 3.903 ± 0.0352 |
| | MAE-MR0.25-CMR-0.5-CellS-ProtS | 0.53 ± 0.0032 | 0.631 ± 0.0026 | 0.718 ± 0.0019 | 4.012 ± 0.0264 |
| | DINO4Cells-HPA | 0.519 ± 0.004 | 0.631 ± 0.0033 | 0.718 ± 0.0026 | 4.047 ± 0.0381 |
| | ViT-Weak-Supervised | 0.544 ± 0.0005 | 0.644 ± 0.001 | 0.727 ± 0.001 | 3.914 ± 0.0172 |
| | bestfitting | **0.606** | **0.663** | **0.768** | **3.049** |

## B.

| Dataset | Experiment | Macro AP | Micro AP |
|---|---|---|---|
| HPAv23 | ProtS | 0.959 ± 0.0025 | 0.99 ± 0.0003 |
| | CellS-ProtS | 0.958 ± 0.0019 | 0.99 ± 0.0002 |
| | MAE-MR0.25-CMR-0.0-ProtS | 0.956 ± 0.0008 | 0.986 ± 0.0001 |
| | MAE-MR0.25-CMR-0.5-ProtS | 0.958 ± 0.0007 | 0.988 ± 0.0001 |
| | MAE-MR0.25-CMR-0.0-CellS-ProtS | **0.974 ± 0.0009** | **0.994 ± 0.0001** |
| | MAE-MR0.25-CMR-0.5-CellS-ProtS | *0.972 ± 0.0006* | *0.993 ± 0.0* |
| | DINO4Cells-HPA | **0.957 ± 0.0014** | **0.99 ± 0.0002** |
| | ViT-Weak-Supervised | 0.552 ± 0.0103 | 0.813 ± 0.0034 |
| | bestfitting | 0.342 ± 0.0038 | 0.682 ± 0.0013 |
| Kaggle | ProtS | 0.948 ± 0.0024 | 0.968 ± 0.0009 |
| | CellS-ProtS | 0.949 ± 0.0019 | 0.964 ± 0.001 |
| | MAE-MR0.25-CMR-0.0-ProtS | 0.93 ± 0.001 | 0.953 ± 0.0009 |
| | MAE-MR0.25-CMR-0.5-ProtS | 0.94 ± 0.001 | 0.957 ± 0.0006 |
| | MAE-MR0.25-CMR-0.0-CellS-ProtS | **0.961 ± 0.0009** | **0.976 ± 0.0005** |
| | MAE-MR0.25-CMR-0.5-CellS-ProtS | *0.957 ± 0.0011* | 0.969 ± 0.0004 |
| | DINO4Cells-HPA | **0.95 ± 0.0014** | **0.964 ± 0.0014** |
| | ViT-Weak-Supervised | 0.531 ± 0.0058 | 0.613 ± 0.0046 |
| | bestfitting | 0.316 ± 0.0037 | 0.42 ± 0.0029 |

**Table S7**: Results for the protein-specific (ProtS) experiments for (A) localization classification and (B) cell-line classification tasks. MAE- refers to the ViT model trained in masked autoencoder setting along with the protein-specific loss.

# A.

| Dataset | Experiment | Macro AP | Micro AP | Label Rank. AP | Coverage Error |
|---|---|---|---|---|---|
| HPAv23 | ProtS | *0.742 ± 0.0047* | *0.878 ± 0.0013* | 0.903 ± 0.0012 | 1.934 ± 0.008 |
| | ProtS-Pool | **0.76 ± 0.002** | **0.881 ± 0.0008** | **0.908 ± 0.0007** | **1.91 ± 0.0039** |
| | MAE-MR0.25-CMR-0.0-CellS-ProtS | 0.729 ± 0.0037 | 0.872 ± 0.0011 | 0.901 ± 0.0012 | 1.959 ± 0.0055 |
| | MAE-MR0.25-CMR-0.0-CellS-ProtS-Pool | 0.739 ± 0.0024 | 0.876 ± 0.0004 | *0.904 ± 0.0005* | *1.941 ± 0.0038* |
| | DINO4Cells-HPA | 0.705 ± 0.0022 | 0.863 ± 0.001 | 0.895 ± 0.0007 | 2.013 ± 0.0059 |
| | ViT-Weak-Supervised | 0.734 ± 0.0008 | **0.869 ± 0.0004** | **0.898 ± 0.0004** | **1.985 ± 0.0029** |
| | bestfitting | **0.77** | 0.83 | 0.877 | 2.029 |
| Kaggle | ProtS | 0.559 ± 0.0058 | **0.653 ± 0.0017** | **0.733 ± 0.0026** | **3.823 ± 0.0464** |
| | ProtS-Pool | **0.565 ± 0.002** | *0.651 ± 0.0013* | *0.731 ± 0.0012* | *3.824 ± 0.0386* |
| | MAE-MR0.25-CMR-0.0-CellS-ProtS | 0.542 ± 0.0036 | 0.64 ± 0.0032 | 0.725 ± 0.0024 | 3.903 ± 0.0352 |
| | MAE-MR0.25-CMR-0.0-CellS-ProtS-Pool | 0.552 ± 0.0018 | 0.647 ± 0.0027 | 0.73 ± 0.0024 | *3.87 ± 0.0379* |
| | DINO4Cells-HPA | 0.519 ± 0.004 | 0.631 ± 0.0033 | 0.718 ± 0.0026 | 4.047 ± 0.0381 |
| | ViT-Weak-Supervised | 0.544 ± 0.0005 | 0.644 ± 0.001 | 0.727 ± 0.001 | 3.914 ± 0.0172 |
| | bestfitting | **0.606** | **0.663** | **0.768** | **3.049** |

# B.

| Dataset | Experiment | Macro AP | Micro AP |
|---|---|---|---|
| HPAv23 | ProtS | 0.959 ± 0.0025 | 0.99 ± 0.0003 |
| | ProtS-Pool | **0.976 ± 0.0011** | 0.993 ± 0.0001 |
| | MAE-MR0.25-CMR-0.0-CellS-ProtS | 0.974 ± 0.0009 | *0.994 ± 0.0001* |
| | MAE-MR0.25-CMR-0.0-CellS-ProtS-Pool | **0.976 ± 0.0009** | **0.995 ± 0.0001** |
| | DINO4Cells-HPA | **0.957 ± 0.0014** | **0.99 ± 0.0002** |
| | ViT-Weak-Supervised | 0.552 ± 0.0103 | 0.813 ± 0.0034 |
| | bestfitting | 0.342 ± 0.0038 | 0.682 ± 0.0013 |
| Kaggle | ProtS | 0.948 ± 0.0024 | 0.968 ± 0.0009 |
| | ProtS-Pool | **0.968 ± 0.0009** | **0.98 ± 0.0005** |
| | MAE-MR0.25-CMR-0.0-CellS-ProtS | 0.961 ± 0.0009 | 0.976 ± 0.0005 |
| | MAE-MR0.25-CMR-0.0-CellS-ProtS-Pool | *0.965 ± 0.0013* | *0.977 ± 0.0006* |
| | DINO4Cells-HPA | **0.95 ± 0.0014** | **0.964 ± 0.0014** |
| | ViT-Weak-Supervised | 0.531 ± 0.0058 | 0.613 ± 0.0046 |
| | bestfitting | 0.316 ± 0.0037 | 0.42 ± 0.0029 |

**Table S8**: Results for evaluating the impact of the attention pooling module on the best models for (A) localization prediction and (B) cell-line classification tasks. MAE- refers to the ViT model trained in masked autoencoder setting along with the protein-specific loss.

| Dataset | Experiment | Macro AP | Micro AP | Label Rank. AP | Coverage Error |
|---|---|---|---|---|---|
| HPAv23 | SubCell-P | **0.577 ± 0.0018** | **0.861 ± 0.0006** | **0.893 ± 0.0008** | **2.18 ± 0.0058** |
| | SubCell | 0.553 ± 0.0022 | 0.856 ± 0.0005 | *0.889 ± 0.0006* | 2.225 ± 0.0052 |
| | DINO4Cells-HPA | 0.509 ± 0.0033 | 0.839 ± 0.0009 | 0.878 ± 0.0007 | 2.36 ± 0.0078 |
| | ViT-Weak-Supervised | 0.534 ± 0.0013 | 0.847 ± 0.0004 | 0.882 ± 0.0004 | 2.316 ± 0.0028 |
| | bestfitting | **0.569 ± 0.0022** | **0.856 ± 0.0007** | **0.888 ± 0.0004** | **2.228 ± 0.0029** |
| Kaggle | SubCell-P | **0.427 ± 0.003** | *0.634 ± 0.0014* | *0.716 ± 0.0015* | *4.466 ± 0.0522* |
| | SubCell | 0.41 ± 0.0021 | 0.631 ± 0.0014 | 0.715 ± 0.0013 | 4.527 ± 0.0332 |
| | DINO4Cells-HPA | 0.376 ± 0.0026 | 0.611 ± 0.0047 | 0.7 ± 0.0035 | 4.863 ± 0.0761 |
| | ViT-Weak-Supervised | 0.399 ± 0.0016 | 0.627 ± 0.0009 | 0.712 ± 0.0007 | 4.565 ± 0.0211 |
| | bestfitting | **0.428 ± 0.0021** | **0.642 ± 0.0019** | **0.725 ± 0.0015** | **4.364 ± 0.0278** |

**Table S9**: Evaluating the performance of the models on a broader range of localization categories. Classification results of the models when evaluating the features on the 31 categories present in the HPA dataset.

| Model | Variant | Channels | Normalization | Accuracy | Macro F1 | Micro F1 |
|---|---|---|---|---|---|---|
| SubCell | DNA-Protein | DNA, Structure | Per Channel | *0.989 ± 0.0001* | 0.871 ± 0.0027 | 0.989 ± 0.0001 |
| | | | Per Image | *0.989 ± 0.0002* | 0.867 ± 0.0026 | 0.989 ± 0.0002 |
| | DNA-Protein-Concat | DNA, Structure, Plasma | Per Channel | **0.990 ± 0.0001** | **0.884 ± 0.0020** | **0.990 ± 0.0001** |
| | | | Per Image | *0.989 ± 0.0001* | 0.861 ± 0.0020 | *0.989 ± 0.0001* |
| | ER-DNA-Protein | DNA, Structure, Plasma | Per Channel | 0.988 ± 0.0001 | 0.852 ± 0.0035 | 0.988 ± 0.0001 |
| | | | Per Image | 0.987 ± 0.0002 | 0.824 ± 0.0036 | 0.987 ± 0.0002 |
| SubCell-P | DNA-Protein | DNA, Structure | Per Channel | 0.981 ± 0.0007 | 0.685 ± 0.0306 | 0.981 ± 0.0007 |
| | | | Per Image | 0.980 ± 0.0006 | 0.662 ± 0.0245 | 0.980 ± 0.0006 |
| | DNA-Protein-Concat | DNA, Structure, Plasma | Per Channel | 0.985 ± 0.0010 | 0.755 ± 0.0291 | 0.985 ± 0.0010 |
| | | | Per Image | 0.984 ± 0.0010 | 0.740 ± 0.0306 | 0.984 ± 0.0010 |
| | ER-DNA-Protein | DNA, Structure, Plasma | Per Channel | 0.981 ± 0.0007 | 0.659 ± 0.0283 | 0.981 ± 0.0007 |
| | | | Per Image | 0.979 ± 0.0005 | 0.629 ± 0.0191 | 0.979 ± 0.0005 |
| DINO4Cells-WTC-11 | - | - | - | 0.988 ± 0.0002 | 0.818 ± 0.0027 | 0.988 ± 0.0002 |
| DINO4Cells-ImageNet | - | - | - | **0.990 ± 0.0002** | *0.882 ± 0.0036* | **0.990 ± 0.0002** |

**Table S10**: Evaluation of SubCell Models with different channel combinations and normalizations on the AllenCell dataset.

| Model | Aggregation | Feature Selection | Norm 1 | Norm 2 | Cell Type | Replicate mAP | MoA mAP |
|---|---|---|---|---|---|---|---|
| CellProfiler | median | True | MAD Robustize | None | A549 | 0.4824 | 0.1740 |
| | | | | | U20S | 0.4329 | 0.1573 |
| DeeProfiler | mean | True | PCAcor | MAD Robustize | A549 | 0.4470 | 0.1697 |
| | | | | | U20S | 0.4164 | 0.1651 |
| DINO4Cells-CP | mean | True | PCA | MAD Robustize | A549 | 0.4218 | 0.1737 |
| | | | | | U20S | 0.3827 | 0.1603 |
| SubCell | median | False | PCA | standardize | A549 | **0.5196** | **0.2267** |
| | | | | | U20S | **0.4671** | **0.2009** |

**Table S11**: Post-processing pipelines chosen for each model in the JUMP analysis.

### 2 Supplementary Figures

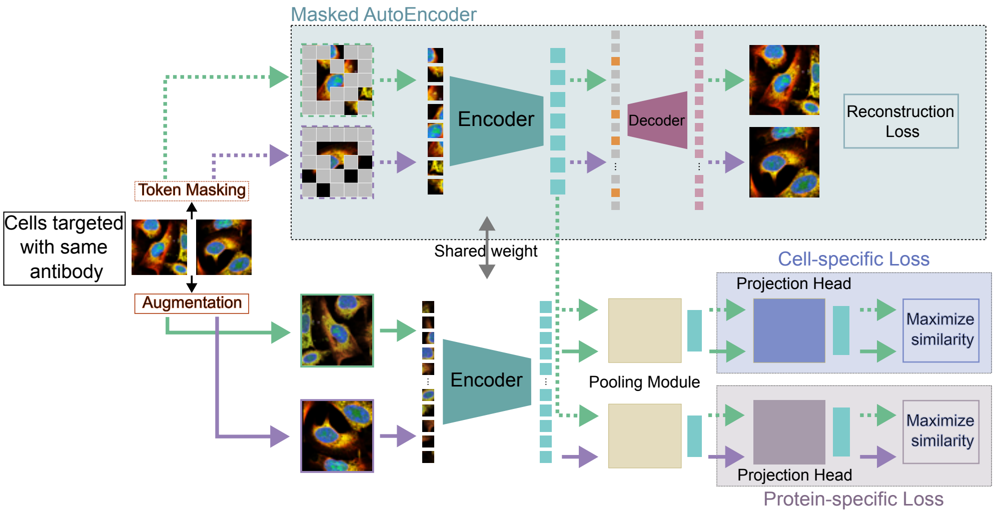


**Fig. S1**: A detailed overview of the multi-task learning framework used in the paper. A batch consisting of multiple images from different antibodies is constructed. One of the views is augmented with geometric augmentations and token masking and fed into the MAE branch. The other view is augmented with geometric and color augmentations and passed through a cell- and protein-specific branch. Finally, the cell-specific and protein-specific loss is calculated over the embeddings from both branches.

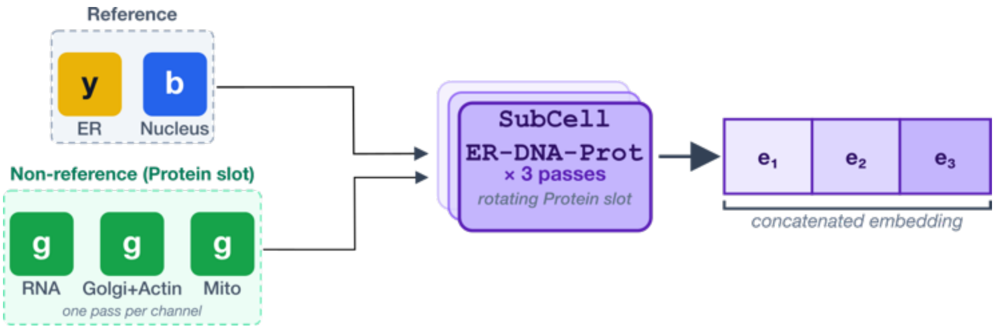


**Fig. S2**: Example illustrating how SubCell extracts features from the JUMP dataset based on the available channels. JUMP samples have two common channels with HPA, nucleus and ER. Three non-reference channels go in the Protein slot. SubCell ER-DNA-Prot model is applied once per non-reference channel. The three embeddings are concatenated into a single representation.

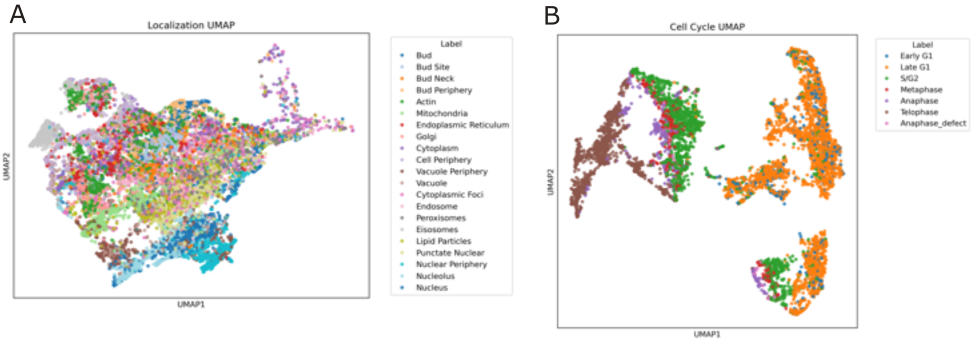


**Fig. S3**: UMAP of SubCell embeddings for yeast images resized to 512x512 (A) colored by protein localization and (B) colored by cell-cycle stage

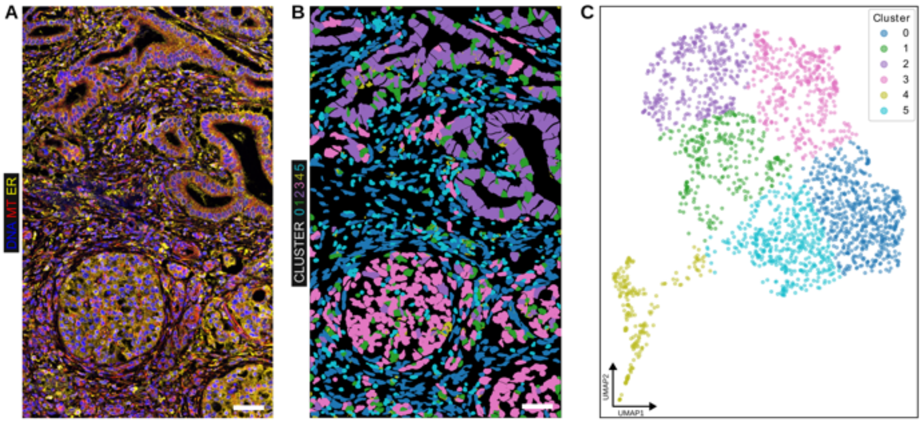


**Fig. S4**: SubCell distinguishes cell types in a tissue section of a patient with pancreatic ductal adenocarcinoma. (A) Stitched confocal fluorescence microscopy images of a pancreatic ductal adenocarcinoma tissue section stained for DNA, MT, and ER. (B) Individual detected cell masks that are used to create single-cell crops, colored by cluster assignment from k-means clustering (k=6) of SubCell embeddings, showing separation of ductal, stromal, and islet cell populations. (C) UMAP visualization of SubCell single-cell embeddings colored by cluster identity. The tissue images were not part of SubCell's training data. Scale bar indicates 50 $\mu m$.

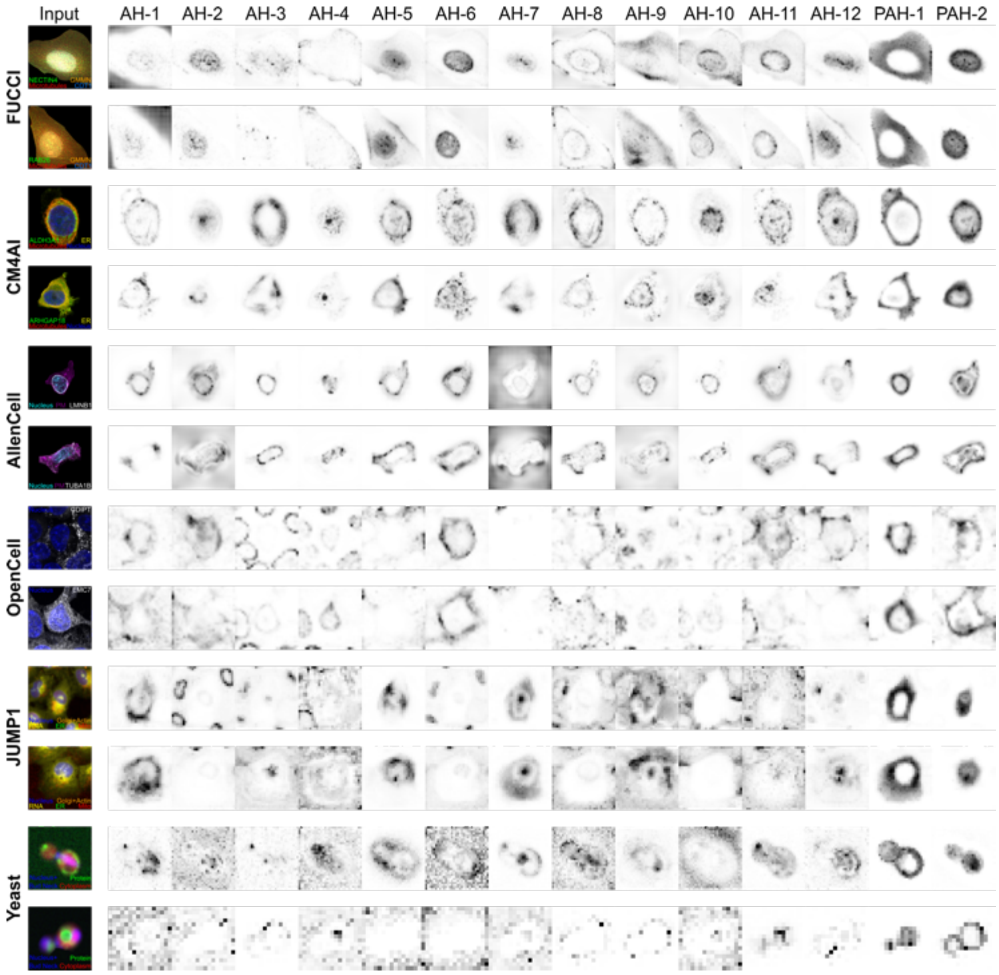


**Fig. S5**: Examples of fluorescent images and the attention captured by the attention heads and pooled attention heads in the SubCell for different datasets. The twelve attention heads of the vision transformers are marked by the prefix AH-, and the pooled attention heads by the prefix PAH-.

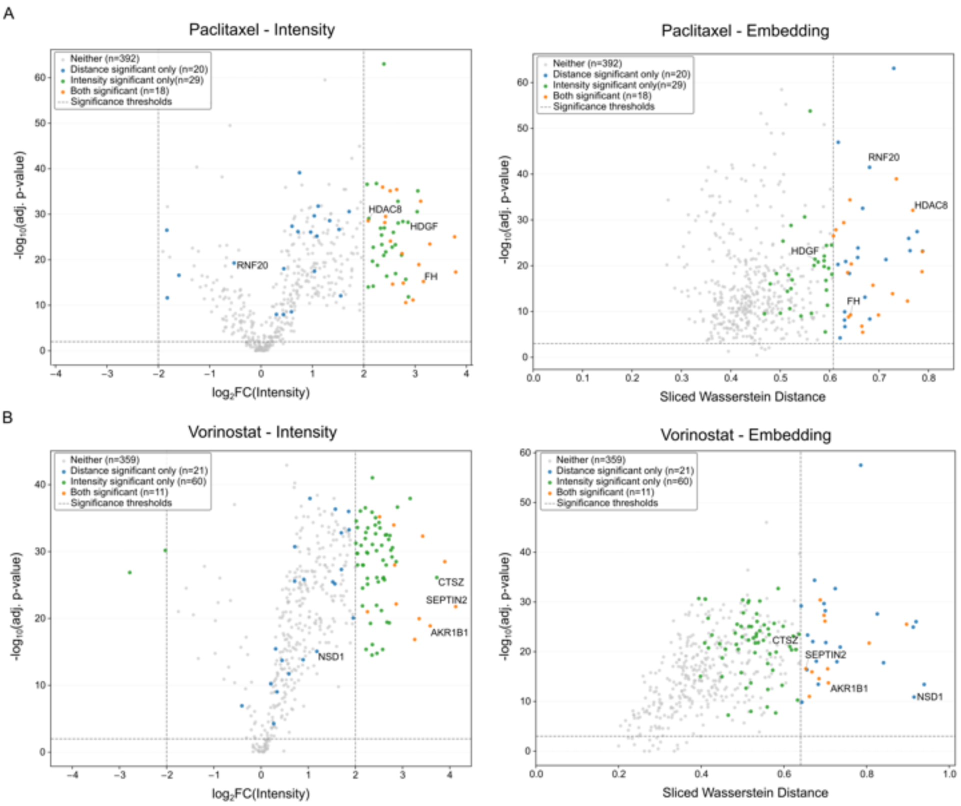


**Fig. S6**: **Volcano plots comparing image intensity and SubCell embedding perturbation signatures for Paclitaxel (A) and Vorinostat (B).** For each drug, proteins are plotted by two complementary measures of drug-induced phenotypic change. **Left:** Intensity volcano: $x$-axis, $\log_2$ fold change in 90th-percentile single-cell image intensity between treated and untreated cells; $y$-axis, $-\log_{10}$ BH-adjusted $p$-value from a two-sided Mann–Whitney U test per protein, with FDR correction across proteins. **Right:** Embedding volcano: $x$-axis, Sliced Wasserstein Distance (SWD) between treated and untreated cell distributions in SubCell embedding space, estimated from 1,000 random projections; $y$-axis, $-\log_{10}$ Bonferroni-adjusted $p$-value, derived by applying a two-sided Mann–Whitney U test per embedding dimension, combining per-dimension $p$-values via Simes' method, and correcting across proteins using the Bonferroni procedure. Points are colored by joint significance (as in Fig. 4C): grey, neither; blue, embedding only; green, intensity only; orange, both. Dashed lines indicate significance thresholds: $|\log_2 \text{FC}| > 2$ and BH-adjusted $p < 0.01$ for intensity; SWD $> \bar{d} + 1.5\sigma$ and Bonferroni-adjusted $p < 0.001$ for embedding. Selected proteins of interest are labeled.

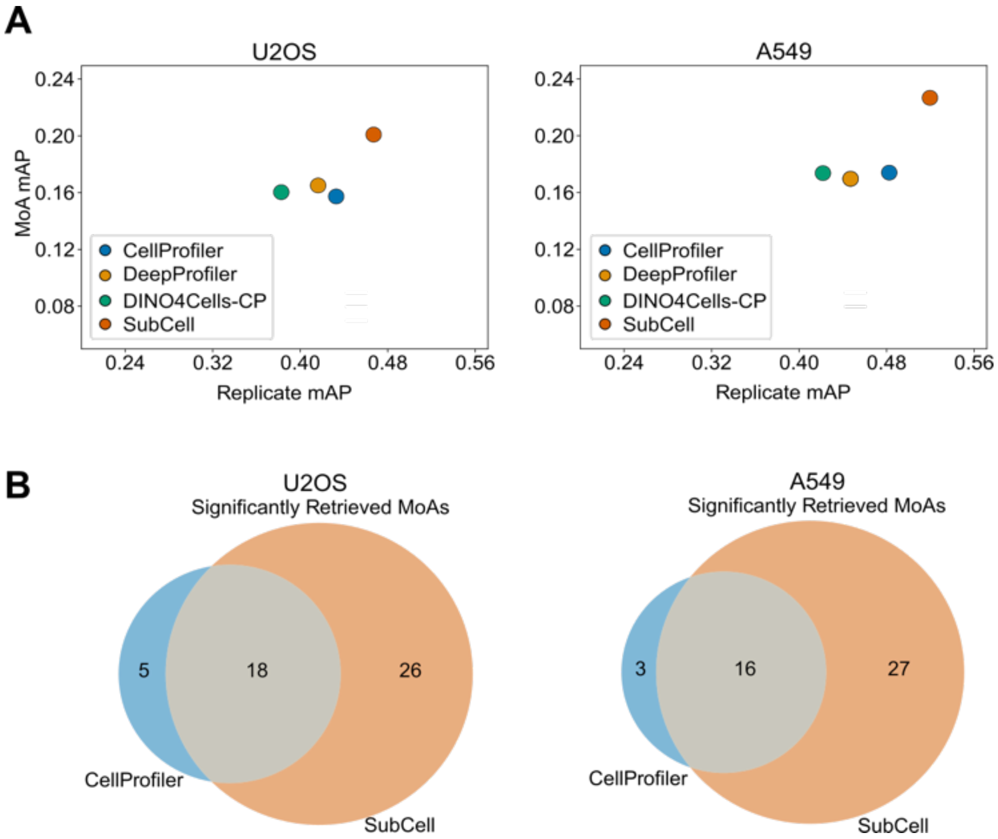


**Fig. S7**: Replicate and MOA retrieval performance on the JUMP1 data. (A) Scatter plots of replicate and MOA retrieval for both U2OS and A549 cells. The copairs package provides a p-values for the mAP values for each MoA. (B) Venn diagram of MoAs significantly retrieved ($p < 0.05$) by SubCell and CellProfiler for both U2OS and A549 cells.

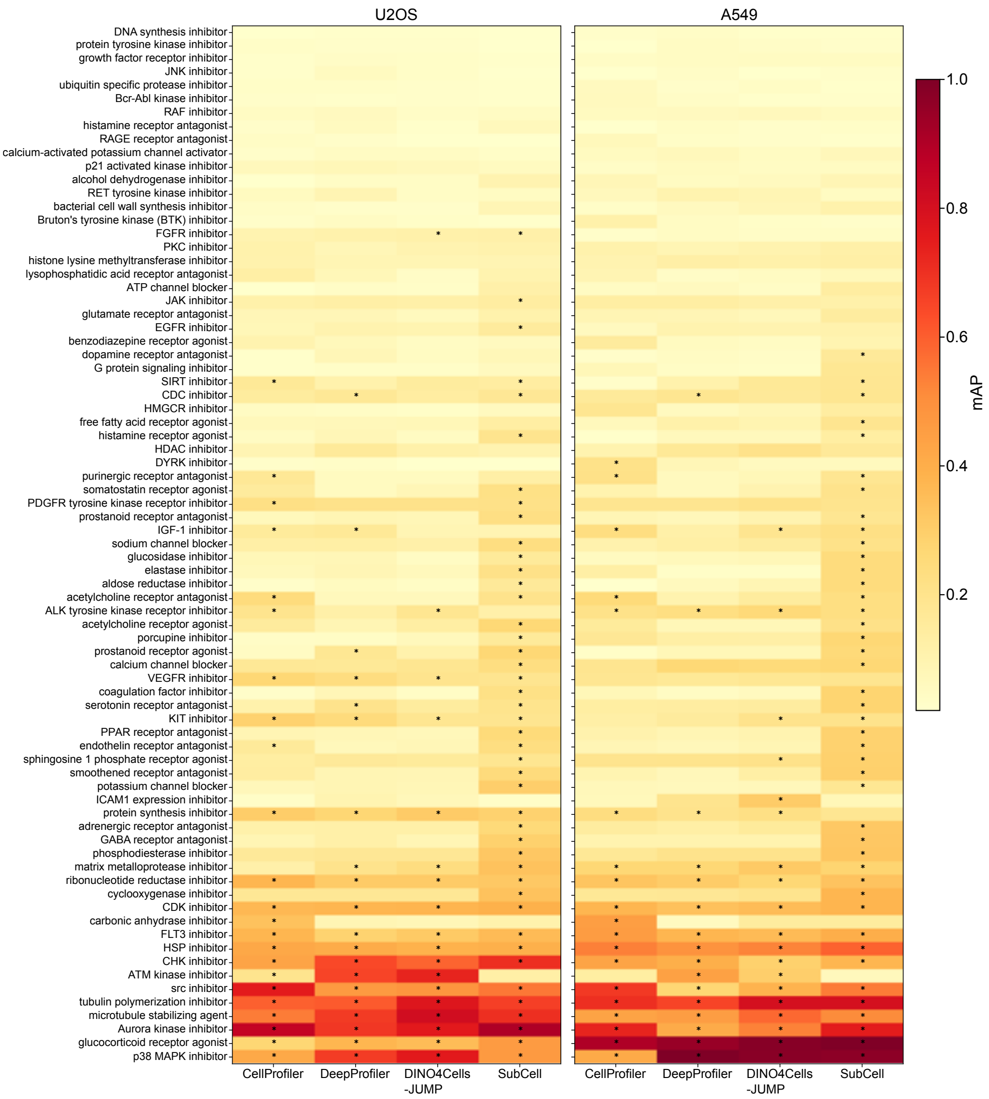


**Fig. S8**: Heatmap of mAP values for MoA retrieval of each model on both cell types. Asterisks indicate mAP values that are statistically significant ($p < 0.05$).

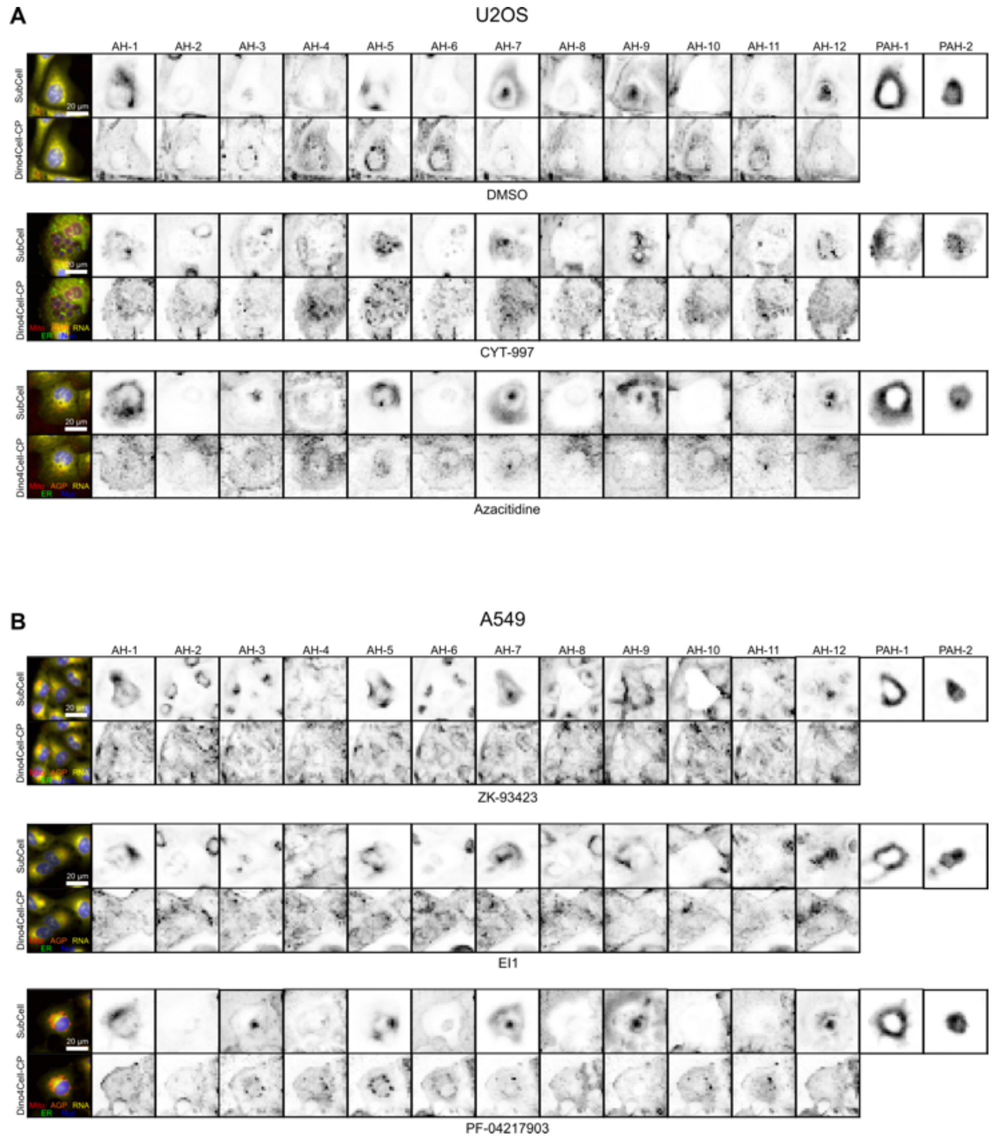


**Fig. S9**: Self-attention maps for SubCell and DINO4Cells-CP on (A) U2OS and (B) A549 cells. One cell crop was randomly selected for 3 random positive-control compounds for each cell-type. Columns show the 12 ViT attention heads for the ViT backbone of each model as well as SubCell's Pooled Attention heads. JUMP cell crops were upscaled to x3.74 to match HPA physical pixel size. Channels are colored as follows: Mitochondria, red; AGP, orange; RNA, yellow; ER, green; Nucleus, blue.

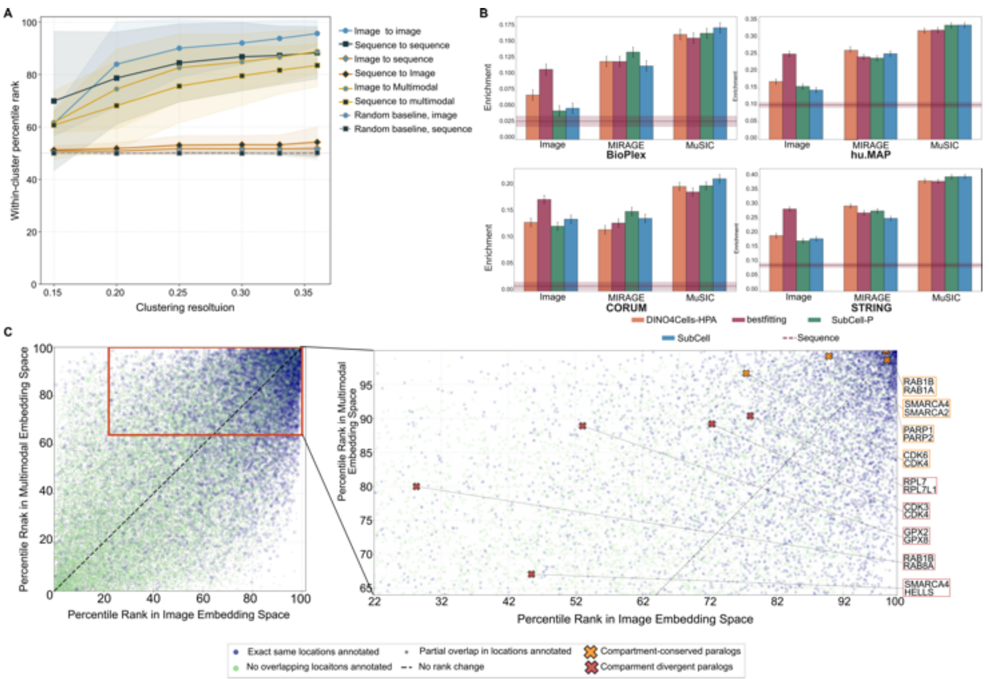


**Fig. S10**: Additional multimodal integration performance. A) Average-ranked similarities of image- and sequence-based cluster labels respectively across different spaces, as a measure of how well different embedding spaces carry information from each. B) Performance of different image models across interaction tasks, using different integration methods (image alone, co-embedded using the MIRAGE method, and co-embedding using the MuSIC method). C) (Right) Scatter plot comparing percentile rank of protein pairs in image vs. multimodal embedding space. Proteins that have the exact same localization pattern are colored in blue, and those that have non-overlapping localization patterns are colored in green. (Left) Examples of redundant (red) and divergent (orange) paralogs highlighted on the percentile rank comparison plot.

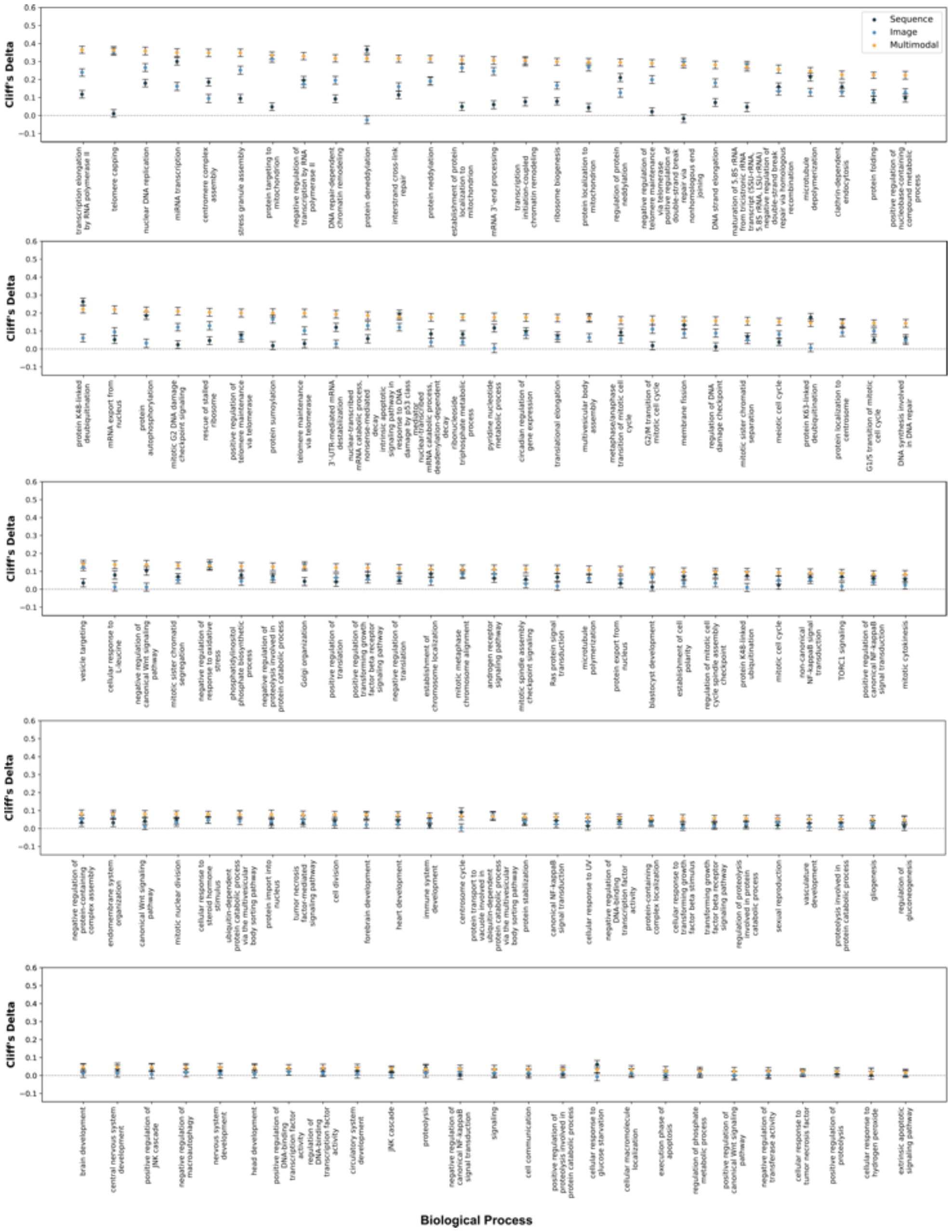


**Fig. S11**: Average embedding similarity (percentile of ranked similarity) of gene members across biological processes. Multimodal space values are shown in yellow, sequence space values are shown in dark blue, and image space values are shown in light blue. The biological processes here are the set of those enriched in our gene set (described in the main section), excluding those shown in Figure 6. They are ordered by the multimodal embedding's value, in decreasing order.

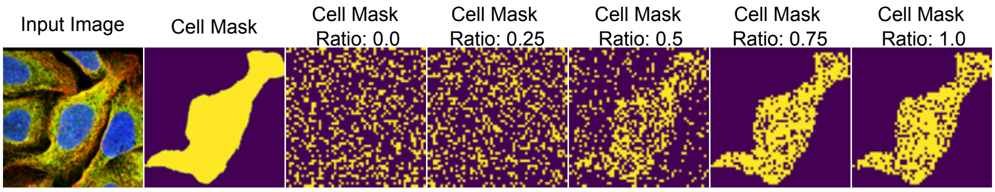


**Fig. S12**: Masking strategy used for removing the tokens in the MAEs using the cell masks. When the cell masking ratio is set to zero, the strategy follows random masking. As the ratio increases, more patches within the cell mask are randomly removed. The overall masking ratio was set to 0.25 for this image.

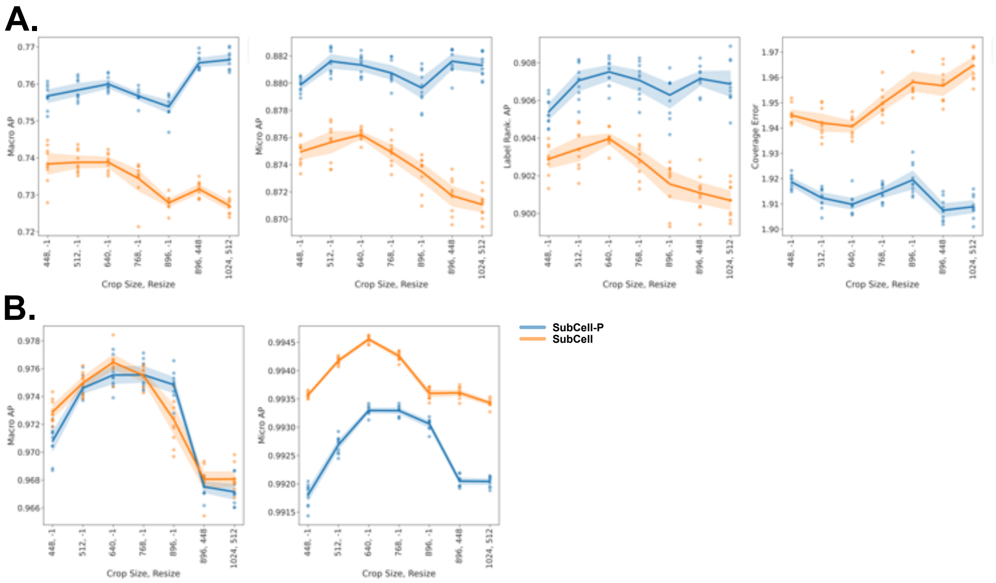


**Fig. S13**: The results of the classifiers trained on the features extracted by the models at different crop sizes and resolutions for A.) localization classification and B.) cell-line prediction tasks.

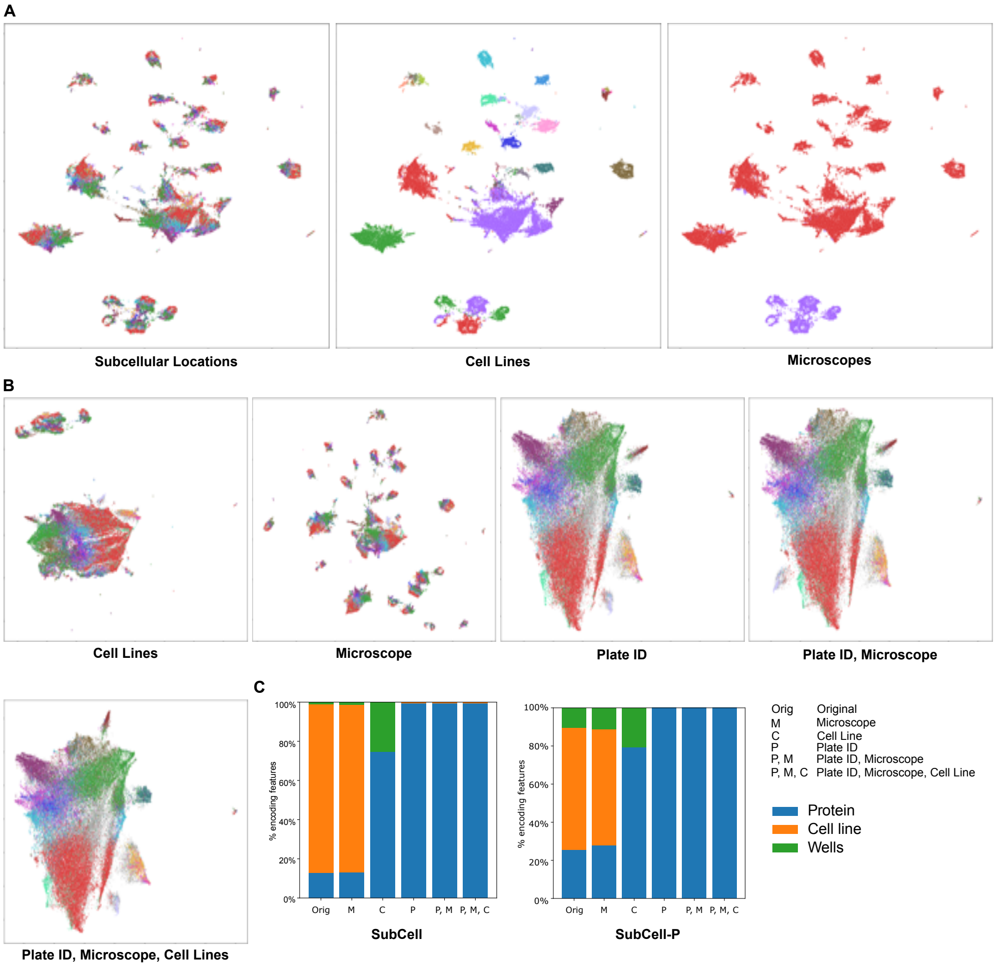


**Fig. S14**: **Visualization of the feature space discovered by the SubCell models**. A) UMAPs of the single-cell features extracted by the SubCell model aggregated by the FOVs colored by subcellular localizations (leftmost), cell lines (middle), and microscope (right). B) The FOV UMAPs showing the resulting features after integrating with Harmony over different factors of variations, namely, cell lines, microscopes, plate IDs, plate IDs and microscopes, and finally with plate IDs, microscopes, and cell lines, with colors representing the subcellular localization categories. C) Stacked-bar plot displaying the fraction of features that are strongly associated with three factors of variation annotated in the HPA dataset for SubCell and SubCell-P.

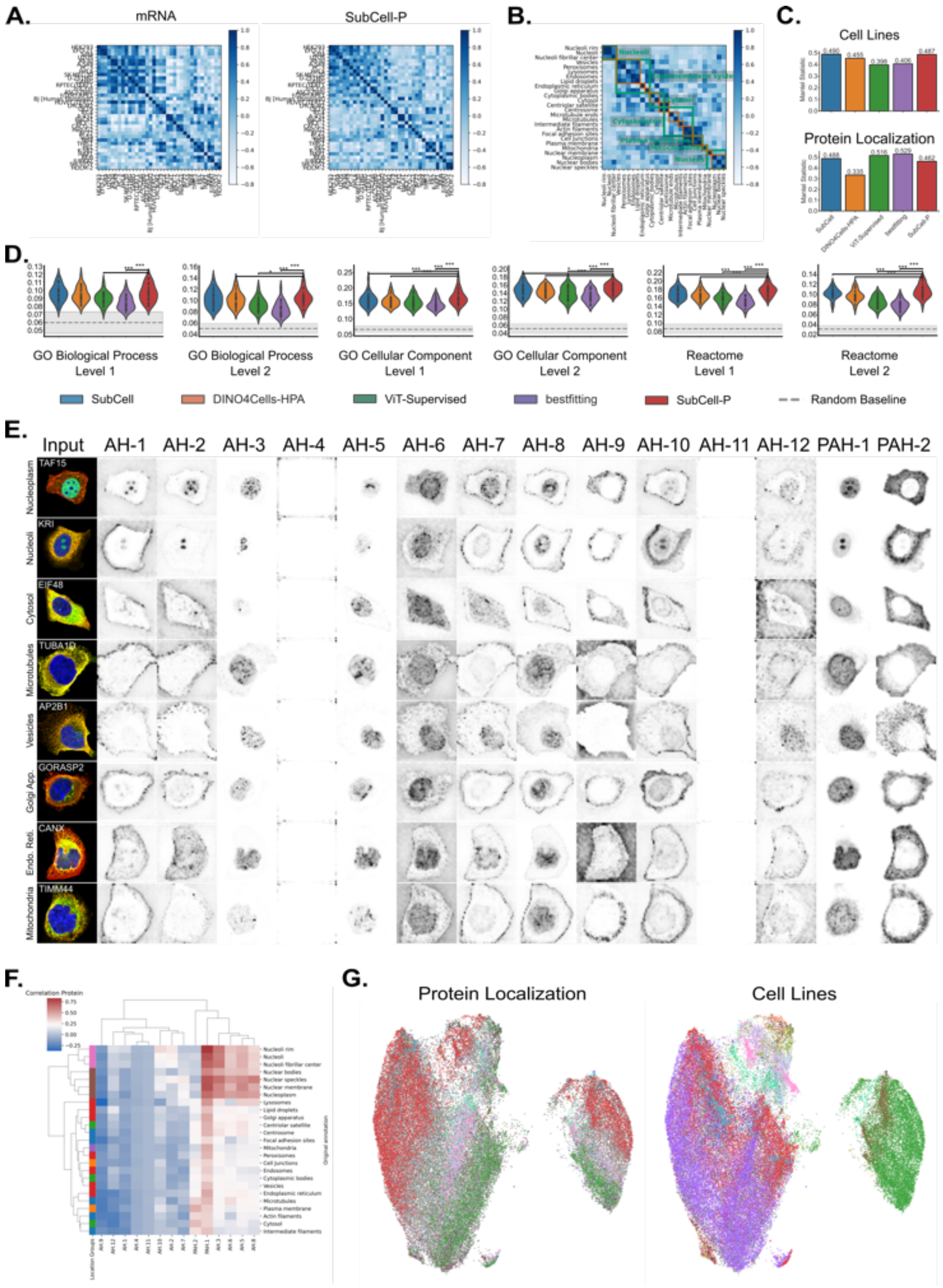


**Fig. S15**: **Feature validation of SubCell-P** (A) Profiling of the cell lines using bulk mRNA levels (left) and aggregated SubCell-P features (right). (B) Cosine similarity among the SubCell-P features aggregated by localization categories. (C) Mantel statistic for the correlation between cell-line-aggregated feature profiles from SubCell-P embeddings and the associated bulk mRNA sequencing profiles (top), and between subcellular localization patterns from SubCell-P and the hierarchical profiles annotated by experts (bottom). (D) Violin plots showing the macro-AP score comparison of the classification performance of the models across different known Level 1 and Level 2 designations denote the hierarchical granularity of the dataset annotations, with Level 1 corresponding to higher-level groupings and Level 2 being more granular (see Methods). (E) Examples of fluorescent images and the attention captured by the attention heads and pooled attention heads in the SubCell-P model for different protein localizations in HPA. (F) A cluster map showing the aggregated Spearman's correlation profiles of the attention heads with the protein channel for different localization categories as defined by expert annotations in C. The order of rows and columns is set by hierarchical clustering of the correlation profiles. The positions of major location groups are shown in different colors on the left side of the plot. (G) Correlation profiles aggregated across all cells within each FOV and plotted using UMAP. The colors represent and annotated protein localization groups (left) and cell lines (right).

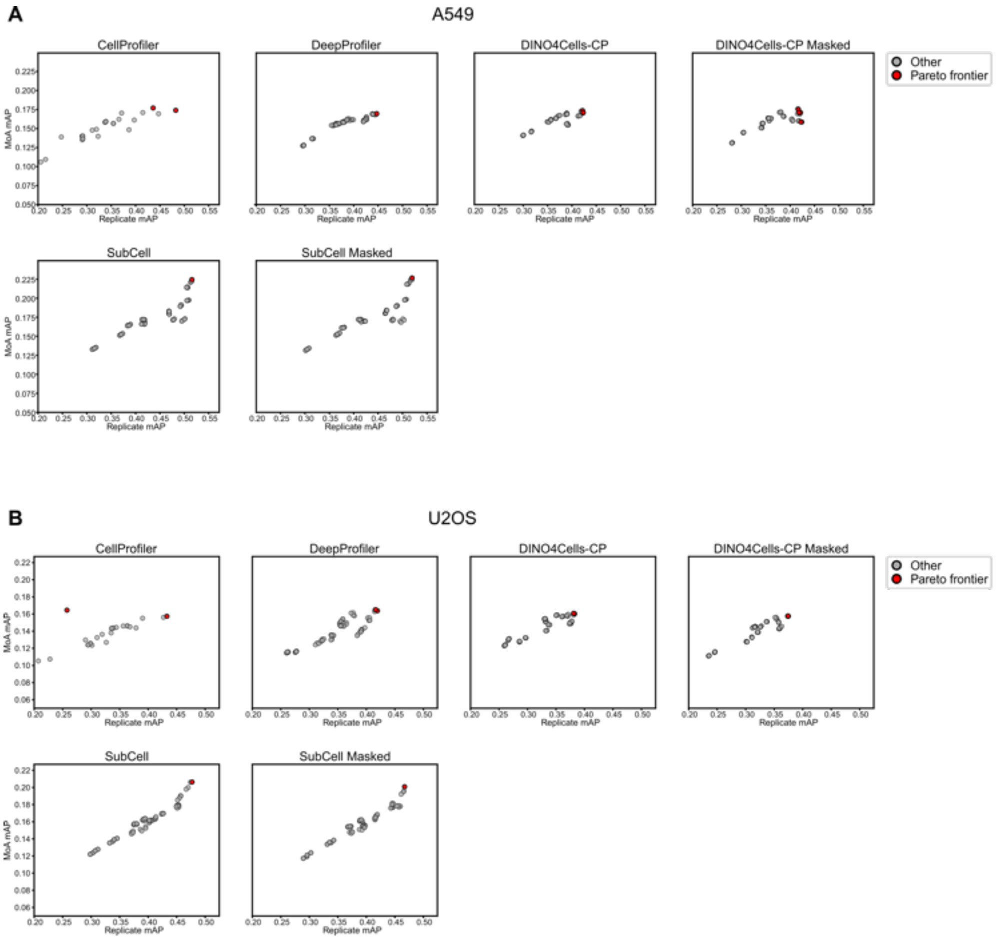


**Fig. S16**: The post-processing configurations for profiles obtained from each model evaluated on the JUMP1 dataset was selected by choosing a configuration whose replicate and MoA mAP value lay on the pareto frontier. Scatter plots of mAP values for all configurations shown for U2OS (A) and A549 (B) cells. Configurations on the pareto frontier shown in red and all other are shown in gray.

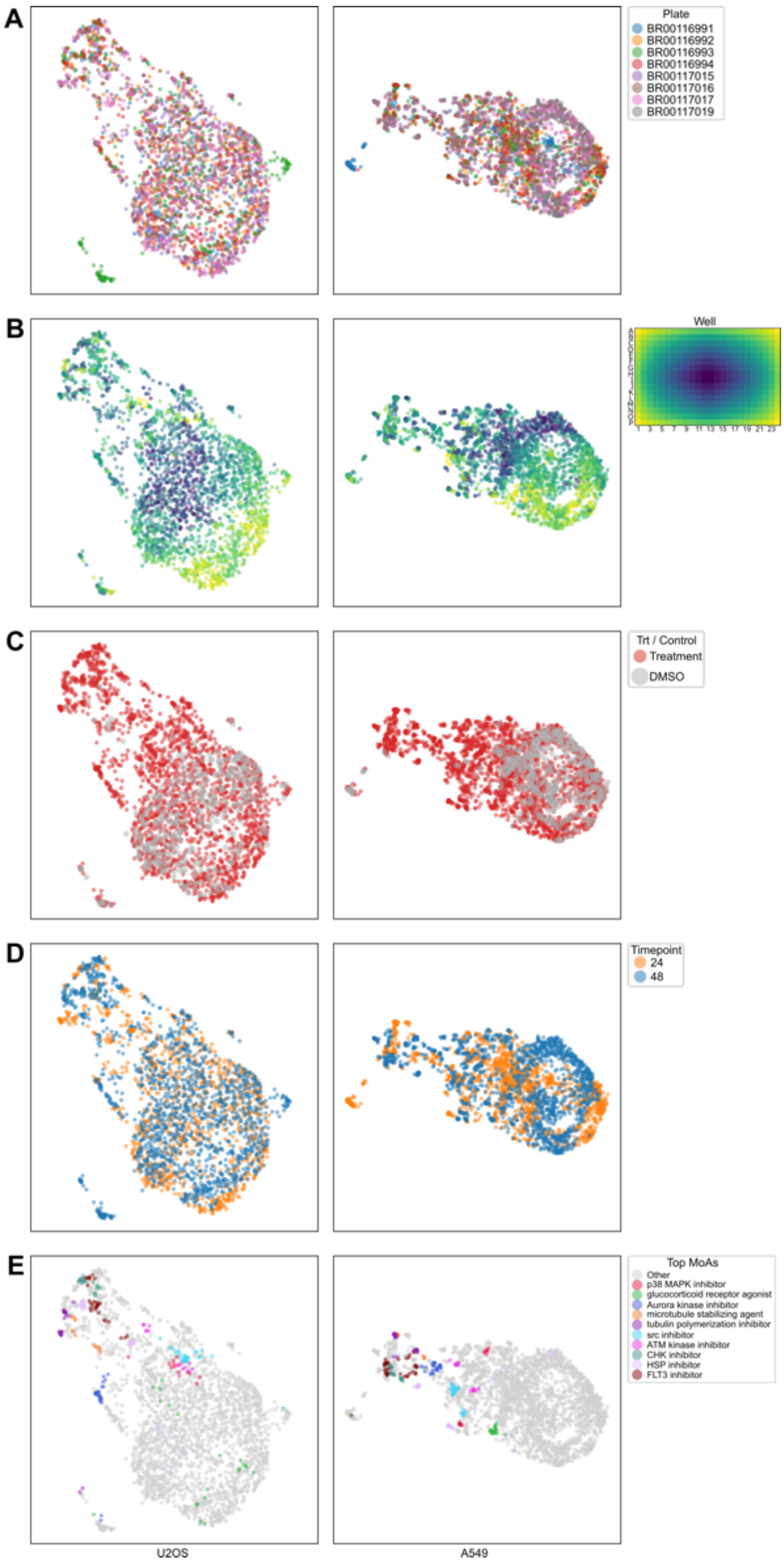


**Fig. S17**: UMAP embeddings of well-level CellProfiler profiles for U2OS and A549 cells from JUMP data. Each column shows one cell type; rows show wells colored by (A) plate, (B) well position, (C) treatment vs. DMSO control, (D) timepoint, and (E) top 10 mechanisms of action, ranked by highest significant mAP across all models and cell types.

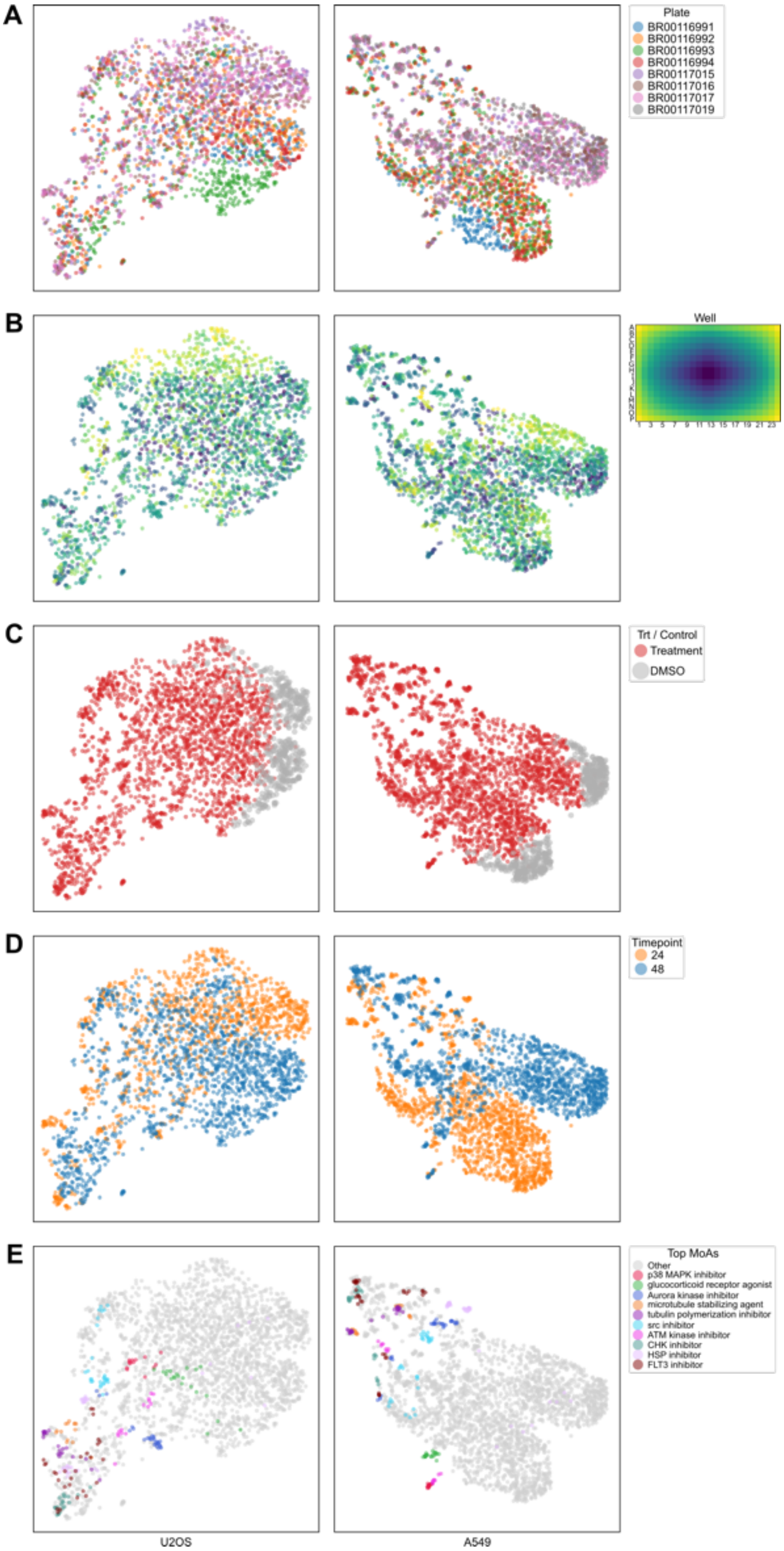


**Fig. S18**: UMAP embeddings of well-level DeepProfiler profiles for U2OS and A549 cells from JUMP data. Each column shows one cell type; rows show wells colored by (A) plate, (B) well position, (C) treatment vs. DMSO control, (D) timepoint, and (E) top 10 mechanisms of action, ranked by highest significant mAP across all models and cell types.

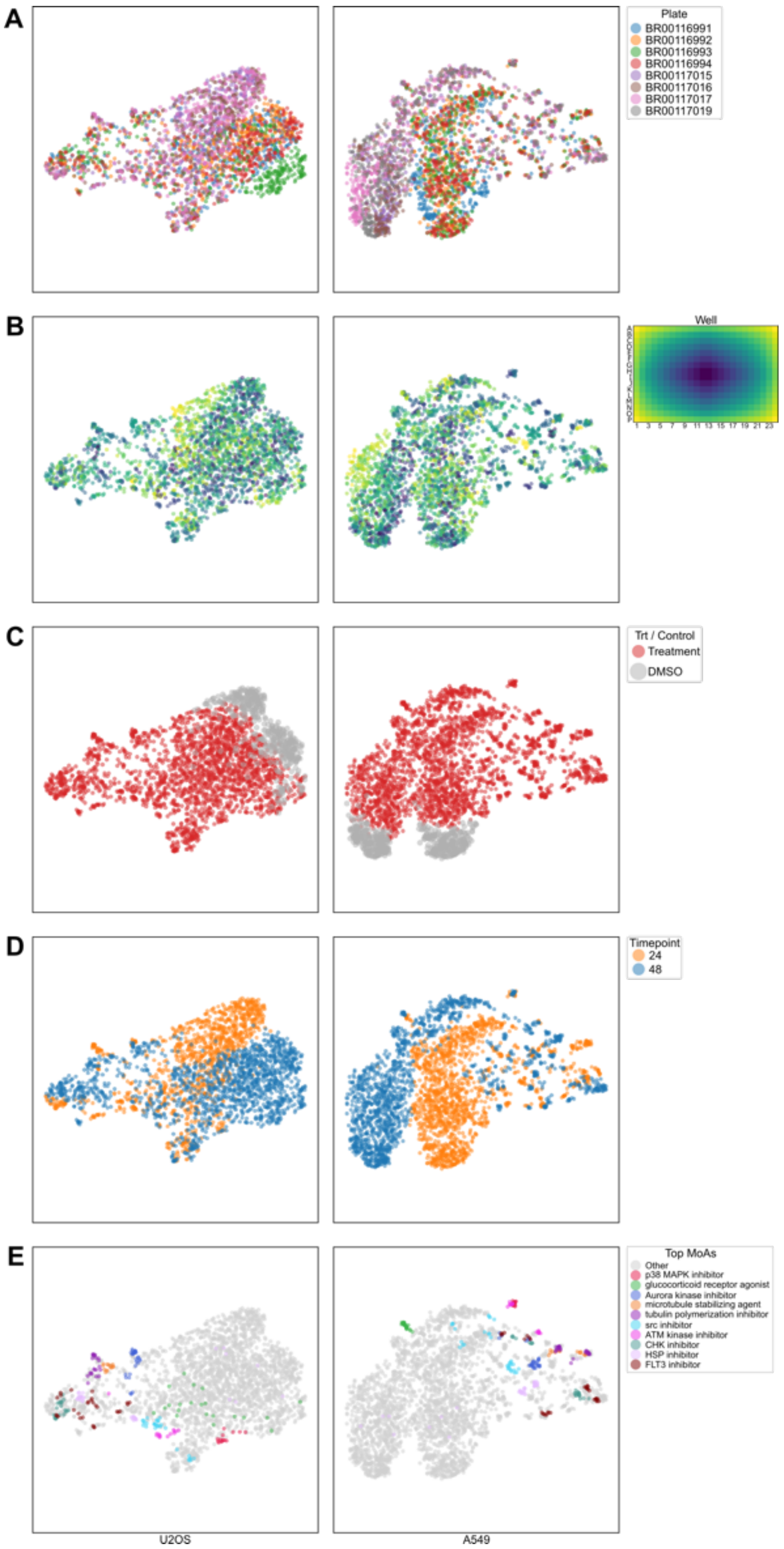


**Fig. S19**: UMAP embeddings of well-level DINO4Cells-CP profiles for U2OS and A549 cells from JUMP data. Each column shows one cell type; rows show wells colored by (A) plate, (B) well position, (C) treatment vs. DMSO control, (D) timepoint, and (E) top 10 mechanisms of action, ranked by highest significant mAP across all models and cell types.

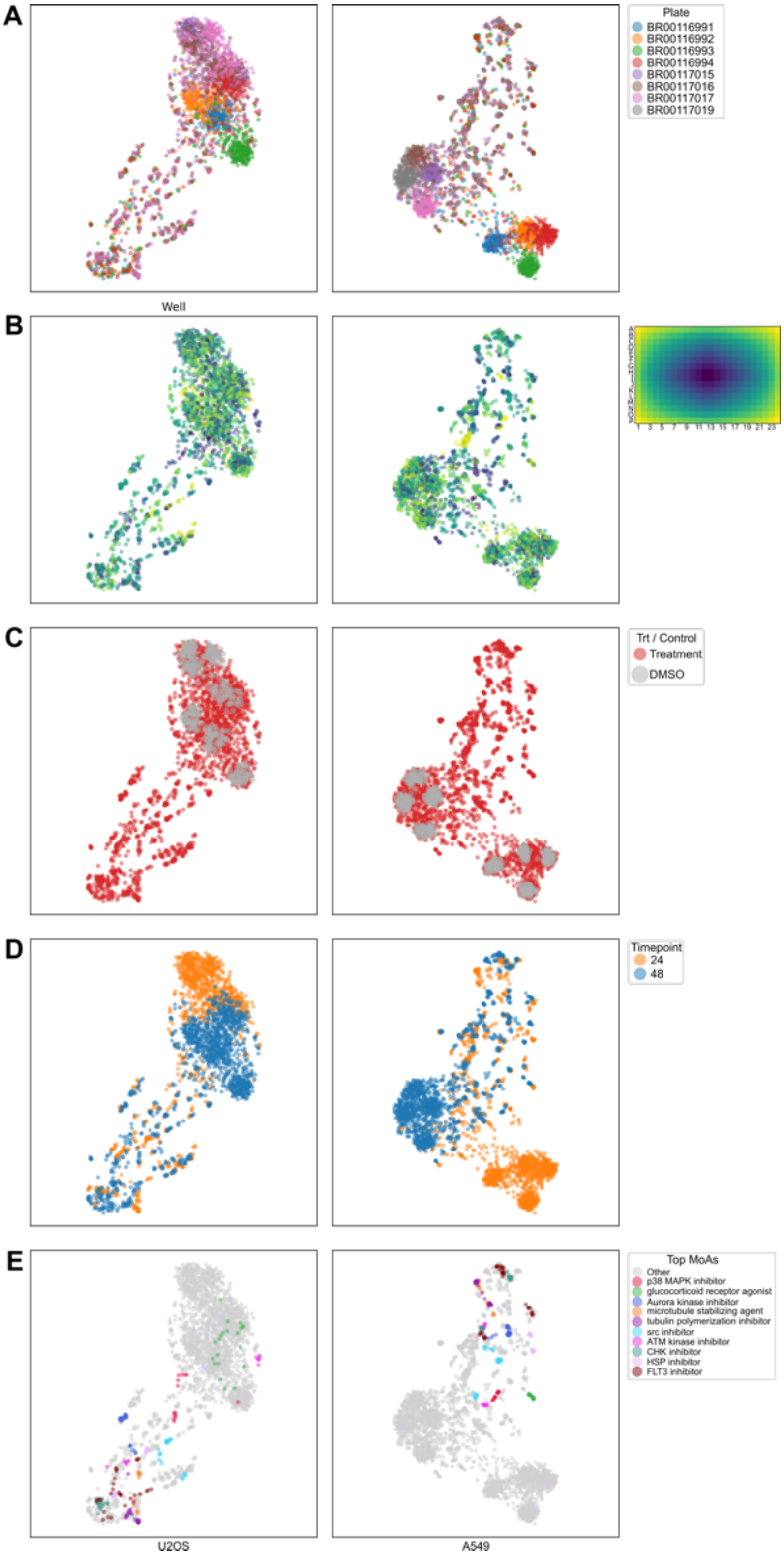


**Fig. S20**: UMAP embeddings of well-level SubCell profiles for U2OS and A549 cells from JUMP data. Each column shows one cell type; rows show wells colored by (A) plate, (B) well position, (C) treatment vs. DMSO control, (D) timepoint, and (E) top 10 mechanisms of action, ranked by highest significant mAP across all models and cell types.
